# Surface Functionality and pH Govern Structural Dynamics and Drug Binding in PETIM and PAMAM Dendrimers

**DOI:** 10.64898/2026.08.04.742721

**Authors:** Anuj Garg, Santosh Mogurampelly, Subbarao Kanchi

## Abstract

Surface functionality and pH play a decisive role in governing the structural dynamics, hydration, and drug-binding behaviour of dendrimers. Here, all-atom molecular dynamics (MD) simulations were performed on five generations of PAMAM (G1-G5) and PETIM (G2-G6) dendrimers with O-core and N-core architectures, functionalized with amine, carboxylic acid, or sugar terminal groups under different protonation states. Protonation of the tertiary branch-point amines expands the dendrimer structure, increases internal porosity and hydration, and enhances structural fluctuations across both families. In contrast, non-protonated amine -NH_2_ (NP) and carboxylic acid -COOH (NP) terminated dendrimers, together with deprotonated carboxylate-COO^-^ (DeP) systems, retain comparatively compact conformations. Sugar-functionalized dendrimers (β-galactose-terminated PETIM and D-glucose-terminated PAMAM) are most hydrated and structurally rigid, whereas amine-terminated dendrimers exhibit the greatest conformational dynamics. PAMAM dendrimers with -NH_2_, -NH_3_^+^, and -COO^-^ terminal groups are generally more hydrated than their PETIM counterparts. However, β-galactose-terminated PETIM dendrimers are more hydrophilic than D-glucose-terminated PAMAM dendrimers. N-core PETIM dendrimers also adopt more compact and spherical conformations than equivalent O-core PETIM dendrimers. Drug-binding MD simulations show that curcumin binding is dominated by van der Waals (vdW) interactions, whereas doxorubicin complexation is primarily driven by electrostatic interactions. Among the investigated surface functionalities, -NH_2_ (NP), -NH_3_^+^ (P), -COOH (NP), and -COO^-^ (DeP) terminations exhibit the most favourable drug-binding characteristics. Except for deprotonated carboxylate systems, curcumin binds more strongly than doxorubicin. Further, drug clustering and spatial organization are governed by dendrimer surface functionality, protonation state, drug loading, and drug-specific interactions. Protonation and higher loading promote drug-drug association, with curcumin and doxorubicin preferentially localizing toward the inner and outer dendrimer regions, respectively. Overall, these findings establish molecular-level relationships between surface functionality, protonation state, dendrimer architecture, and drug-binding behaviour, providing design principles for pH-responsive dendrimer nanocarriers with enhanced drug-loading and controlled-release performance.

## 2. Introduction

Dendrimers are monodisperse, highly branched macromolecules possessing a well-defined three-dimensional architecture, internal cavities, and a large number of terminal functional groups. These structural characteristics have made them attractive nanocarriers for diverse applications, including drug delivery, catalysis, sensing, and nanotechnology ^1–5^. Their physicochemical behaviour is governed by both the dendrimer architecture and the chemistry of the surface functional groups. Recent experimental studies demonstrate that the internal branching pattern of dendrimers determines their molecular size, flexibility, and internal porosity, whereas the terminal functional groups regulate their interactions with the surrounding environment, thereby influencing solubility, hydration, reactivity, and molecular recognition ^6^. Consequently, tailoring the dendrimer surface provides an effective strategy for designing nanocarriers with properties suited to specific biomedical and industrial applications ^7–11^.

Recent studies demonstrate that the internal branching pattern of dendrimers determines their molecular size, flexibility, and internal porosity, whereas the terminal functional groups govern their interactions with the surrounding environment, thereby influencing solubility, hydration, reactivity, and molecular recognition. Consequently, tailoring the dendrimer surface provides an effective strategy for designing nanocarriers with properties suited to specific biomedical and industrial applications

Among the various surface chemistries, amine (-NH₂), carboxylic acid (-COOH), and carbohydrate-derived sugar moieties have attracted considerable attention because they impart distinct physicochemical and biological characteristics. Amine-terminated dendrimers possess positively charged surfaces under physiological conditions, enabling strong electrostatic interactions with negatively charged biomolecules such as nucleic acids and proteins. This property has led to their extensive use in gene delivery, drug encapsulation, biosensing, and nanoparticle functionalization ^12–14^. However, their high cationic charge density may also induce membrane disruption and cytotoxicity, motivating the development of alternative surface chemistries or controlled surface modifications to improve biocompatibility ^15^. Carboxylic acid-terminated dendrimers exhibit anionic or neutral surface characteristics depending on the solution pH, generally resulting in lower cytotoxicity and enhanced aqueous stability ^16^. Their terminal carboxyl groups readily participate in hydrogen bonding and ionic interactions, facilitating the formation of stable drug complexes while simultaneously providing reactive sites for further chemical functionalization. These properties have enabled their application in antimicrobial systems, metal nanoparticle stabilization, and polymer-drug conjugates, including poly (ethylene glycol)-modified dendrimers for carboplatin delivery ^16,17^. Sugar-functionalized dendrimers represent another important class of biomimetic nanocarriers. Terminal carbohydrates such as β-galactose and D-glucose resemble naturally occurring glycoconjugates, promoting favourable interactions with cell membranes, serum proteins, and carbohydrate-binding receptors ^18^. Their highly hydrophilic surfaces improve water solubility, biocompatibility, and colloidal stability while enabling receptor-mediated cellular recognition. Consequently, glycodendrimers have been explored for nucleic acid delivery, antiviral and antibacterial therapies, inhibition of urinary tract infections, and targeted delivery of small interfering RNA (siRNA) ^9,19–22^.

Besides surface chemistry, the protonation state of dendrimers plays a crucial role in determining their structural and functional properties. Protonation of internal tertiary amines and terminal functional groups alters electrostatic interactions within the dendrimer, leading to changes in molecular size, shape, flexibility, internal hydration, and cavity accessibility. Since these protonation states are governed by environmental pH, dendrimers exhibit pH-responsive behaviour that directly influences their ability to encapsulate and release therapeutic molecules. A molecular-level understanding of how surface functionality and protonation collectively regulate dendrimer conformation is therefore essential for the rational design of efficient drug delivery systems.

Curcumin and doxorubicin are two widely studied anticancer agents whose therapeutic performance is limited by poor pharmacokinetic properties. Curcumin possesses broad-spectrum anticancer activity but suffers from extremely low aqueous solubility and limited bioavailability, whereas doxorubicin, although highly effective against a wide range of cancers, is associated with systemic toxicity and dose-dependent side effects. Encapsulation within dendrimers offers an effective strategy to overcome these limitations by improving drug solubility, stability, circulation time, and controlled release ^23,24^. Owing to its largely hydrophobic and neutral character under physiological conditions, curcumin interacts predominantly through hydrophobic and van der Waals interactions within the dendrimer interior ^25^. In contrast, doxorubicin contains ionisable functional groups, and its complexation strongly depends on electrostatic interactions that vary with the protonation state of both the drug and the dendrimer ^26,27^. Moreover, incorporation of pH-sensitive linkers or pH-responsive dendrimer architectures enables preferential drug release within the acidic tumour microenvironment, thereby enhancing therapeutic efficacy while reducing toxicity toward healthy tissues ^23,25,27^.

Despite extensive studies on dendrimer–drug interactions, several aspects remain insufficiently understood, including (i) the influence of dendrimer generation on structural organization, size, internal accessibility, and drug interactions; (ii) the role of surface functional groups and their protonation states in regulating dendrimer conformation, hydration, and drug binding; (iii) the effects of dendrimer structure and surface chemistry on the stability and energetics of drug–dendrimer complexes; and (iv) the contributions of direct molecular contacts and solvent-mediated interactions to drug association. The present study addresses these gaps through a systematic investigation of O-core and N-core PETIM dendrimers alongside PAMAM dendrimers bearing amine, carboxylic acid, and sugar terminal groups under different protonation states. Their interactions with neutral trans-curcumin and positively charged doxorubicin were subsequently analyzed to elucidate the molecular mechanisms governing drug binding. By correlating dendrimer architecture, surface chemistry, protonation state, and drug-binding energetics, this study provides molecular design principles for optimizing dendrimer-based nanocarriers for pH-responsive drug delivery.

## 3. Simulation Methodology

The initial molecular structures of amine-terminated PAMAM and PETIM dendrimers were generated using the Dendrimer Building Toolkit (DBT) ^28^. The geometries of the PETIM repeating unit together with the terminal carboxylic acid and sugar (D-glucose and β-galactose) residues were optimized for their respective protonation states using Gaussian09 ^29^ shown in **Figure S1**. Quantum mechanical calculations were performed at the MP2/6-31+G(d) level of theory to obtain the minimum-energy geometries. Atomic partial charges were derived from the electrostatic potential (ESP) of the optimized structures and subsequently fitted using the restrained electrostatic potential (RESP) procedure implemented in the Antechamber module of the AMBER package ^30^. The capped repeating units and terminal functional groups were assigned net charges of 0, +1, and −1 corresponding to their non-protonated (NP), protonated (P), and deprotonated (DeP) states, respectively. Using these optimized building blocks, PAMAM (G1–G5) and PETIM (G2–G6) dendrimers were constructed with different surface functionalities. The amine-functionalized dendrimers were considered in NH₂ (NP), NH₃⁺ (P), and fully protonated (DP) forms, whereas the carboxyl-functionalized dendrimers were modelled as COOH (NP and P) and COO⁻ (DeP). Sugar-functionalized dendrimers comprised β-galactose-terminated PETIM and D-glucose-terminated PAMAM in both non-protonated and protonated states. The two-dimensional representations of O-core PETIM dendrimers with different terminal functional groups, including NH₂-terminated G3, COOH-terminated G3.5, and β-galactose-terminated G3.5 dendrimers, are shown in **Figure S2** to illustrate their differences in molecular architecture and surface functionality. For the dendrimer– drug systems, RESP-derived atomic partial charges were similarly obtained for neutral trans-curcumin and positively charged doxorubicin, corresponding to net molecular charges of 0 and +1, respectively. The General AMBER Force Field (GAFF) parameters were assigned to both drug molecules ^31^.

All-atom molecular dynamics simulations were carried out using the GROMACS simulation package ^32^ with the AMBER14SB force field ^33^. Each dendrimer was solvated in an explicit TIP3P water box ^34^, ensuring a minimum distance of 1.0 nm between the solute and the simulation box boundaries. To reproduce physiological ionic conditions, Na⁺ and Cl⁻ ions were introduced to neutralize the systems and achieve a salt concentration of 0.15 M. Before the production simulations, each system was energy minimized using the steepest-descent algorithm for 50,000 steps to eliminate unfavourable steric contacts. This was followed by successive equilibration under the canonical (NVT) and isothermal–isobaric (NPT) ensembles for 100ps each with an integration time step of 2 fs. Throughout the simulations, the temperature and pressure were maintained at 300 K and 1 atm using the Berendsen thermostat ^35^ and the Parrinello–Rahman barostat ^36^, respectively to allow physically realistic volume and pressure fluctuations during the NPT production runs. Long-range electrostatic interactions were evaluated using the Particle Mesh Ewald (PME) method ^37^, while all covalent bonds involving hydrogen atoms were constrained using the LINCS algorithm^38^. During equilibration, positional restraints of 1000 kcal mol⁻¹ Å⁻² were applied to the dendrimer atoms to preserve their structural integrity. These restraints were subsequently released before the production simulations.

Following equilibration, each dendrimer system was subjected to a 100 ns production molecular dynamics simulation to investigate its structural and dynamical properties. Dendrimer– drug complexes were prepared by placing five drug molecules around a single dendrimer, followed by 200 ns production simulations to examine the complexation behaviour. The radius of gyration (Rg) time profiles for the higher-generation G6 O-core PETIM and G5 PAMAM dendrimers are presented in **Figure S3**. The Rg profiles indicate that both dendrimer structures reach a relatively stable and equilibrated state within approximately 40 ns, confirming adequate structural relaxation before the subsequent analyses. Structural analyses together with the molecular mechanics Poisson– Boltzmann surface area (MM/PBSA) binding free-energy calculations were performed using the final 10 ns of the equilibrated trajectories, ensuring that all reported properties correspond to well-converged simulation ensembles. The simulation methodology detailed previously successfully generated stable trajectories in thermodynamic equilibrium for systems including dendrimers, polymer nanoparticles, proteins, and lipid membranes ^39–44^. The structural properties of dendrimers, including the radius of gyration (Rg), asphericity (δ), bound water molecules (n_inner_, n_surf_, n_bulk_), and root-mean-square-fluctuations (RMSF), were analysed using the methodology described in our previous study ^40^.

## 4. Results and discussion

In this study, the structures of poly-ether imine (PETIM) dendrimers with oxygen-core (O-core) and nitrogen-core (N-core) architectures were examined as a function of their size, pH, and surface functionalization (amine, carboxylic, and sugar groups). These dendrimers were compared with the well-known poly-amido amine (PAMAM) dendrimers synthesized with an ethylene-diamine (EDA) core. For amine-terminated dendrimers, primary amines remain non-protonated (charge = 0) at basic pH (≥10.0) but become protonated (charge = +1.0e) at neutral pH (∼7.0). The pKa values of the protonated primary amine groups in PAMAM dendrimers typically range from approximately 7 to 9. In addition, the internal tertiary amine groups can become protonated under acidic conditions (pH ≤ 5), with reported pKa values generally ranging from approximately 3 to 6^45^. Dendrimers with carboxylic terminal groups are predominantly deprotonated (charge = -1.0e) at pH ≥ 5.0, consistent with reported pKa values in the range of 3-4.4 ^46^. In contrast, the terminal groups of sugar-functionalized dendrimers, such as β-galactose and D-glucose, remain essentially neutral over the physiologically relevant pH range. However, the tertiary amines of both carboxylic and sugar terminal groups will be protonated at pH≤5.0. Further, the protonation (net positive charge) or deprotonation (net negative charge) of amine, carboxylic or sugar terminal groups significantly modifies the drug release profiles of dendrimers compared to their non-protonated (net zero charge) states. Furthermore, drug-binding to dendrimers may alter the local chemical environment of the ionisable groups and, consequently, influence their effective pKa values in dendrimer-drug complex ^45^. To elucidate these effects, various structural parameters including the radius of gyration (Rg), aspect ratios, asphericity, radial density distributions, hydration number, and root-mean-square-fluctuations (RMSFs) were analysed across different protonation states. This comprehensive analysis highlights the impact of pH on the structural characteristics and surface functionality of PETIM and PAMAM dendrimers, offering valuable insights into their potential for drug delivery applications.

### 4.1. Structure of PETIM dendrimers

Poly-ether imine (PETIM) dendrimers are classified into two types based on the central core atom: oxygen-core (O-core) and nitrogen-core (N-core). The O-core PETIM dendrimers are synthesized by branching into two arms, while the N-core PETIM dendrimers grow into three arms, corresponding to the multiplicity of their central oxygen and nitrogen atoms, respectively. In this study, the structures of O-core and N-core PETIM dendrimers across generations 2 to 6 were analysed with surface functionalization of amine (-NH_2_), carboxylic (-COOH), and β-galactose (-C_6_H_12_O_6_) at different protonation states (pH) using all-atom molecular dynamics (MD) simulations.

The final thermodynamically equilibrated simulation snapshots of dendrimers with various terminations are shown in **Figures 1a** and **1b** for O-core and N-core PETIM dendrimers, respectively. As the generation number increases, the number of branched layers grows, leading to a corresponding increase in dendrimer size across all surface terminations. Protonated dendrimers (net positive charge) with amine or β-galactose surface groups adopt an open confirmation, stretching out their exterior branch units. In contrast, deprotonated dendrimers (net negative charge) with carboxylic terminations exhibit a more compact structure compared to their neutral counterparts (net charge = 0) for respective functionalization. This behaviour can be attributed to differences in the screening of the charged surface terminal groups.

**Figure 1.**
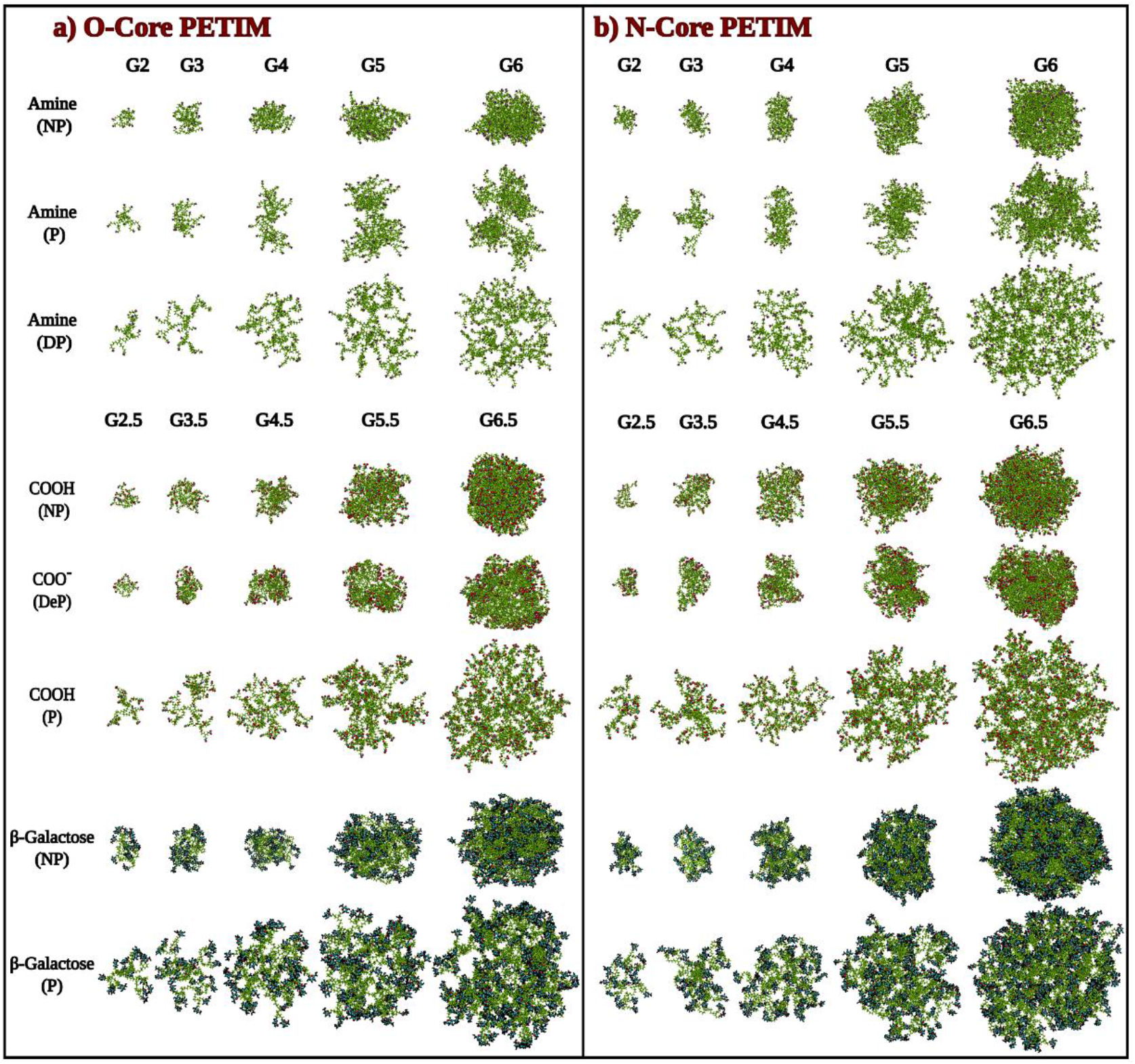
Final simulation snapshots of NH₂ (NP), NH₃⁺ (P), NH₃⁺ (DP), COOH (NP), COO⁻ (DeP), COOH (P), β-galactose (NP), and β-galactose (P)-terminated (G2–G6) (a) oxygen-core (O-core) and (b) nitrogen-core (N-core) PETIM dendrimers after 100 ns of molecular dynamics (MD) simulations are presented. Dendrimers functionalized with NH₃⁺ (DP), COOH (P), and β-galactose (P) adopt more expanded conformations and exhibit the highest internal porosity, primarily due to strong electrostatic repulsions between the positively charged protonated tertiary amines located at the branch points. In contrast, electrostatic repulsion among the negatively charged COO⁻ (DeP) terminal groups is effectively screened by Na⁺ counter ions, resulting in more compact structures than those observed for dendrimers with COOH (NP) termination.

#### 4.1.1. Size of PETIM dendrimers

The R_g_ profiles were computed for O-core and N-core PETIM dendrimers (**Figures 2a** and **2b**) with varying surface functionalisation and protonation states across different generations. This analysis explores how surface functionalities influence dendrimer size. As the number of branching layers increases with generation, the Rg values grow correspondingly. The comparison of Rg profiles reveals that the impact of protonation levels on size is more pronounced in amine terminated dendrimers than in carboxylic and β-galactose terminated dendrimers. Among all variations, the non-protonated (NP) amine terminated dendrimers exhibit smallest size. The sizes of both non-protonated (-COOH) and deprotonated (-COO^-^) carboxylic groups dendrimers are comparable to those of NP amine terminated dendrimers. Additionally, β-galactose terminated dendrimers are larger than their amine and carboxylic terminated counterparts in all corresponding protonation states.

**Figure 2.**
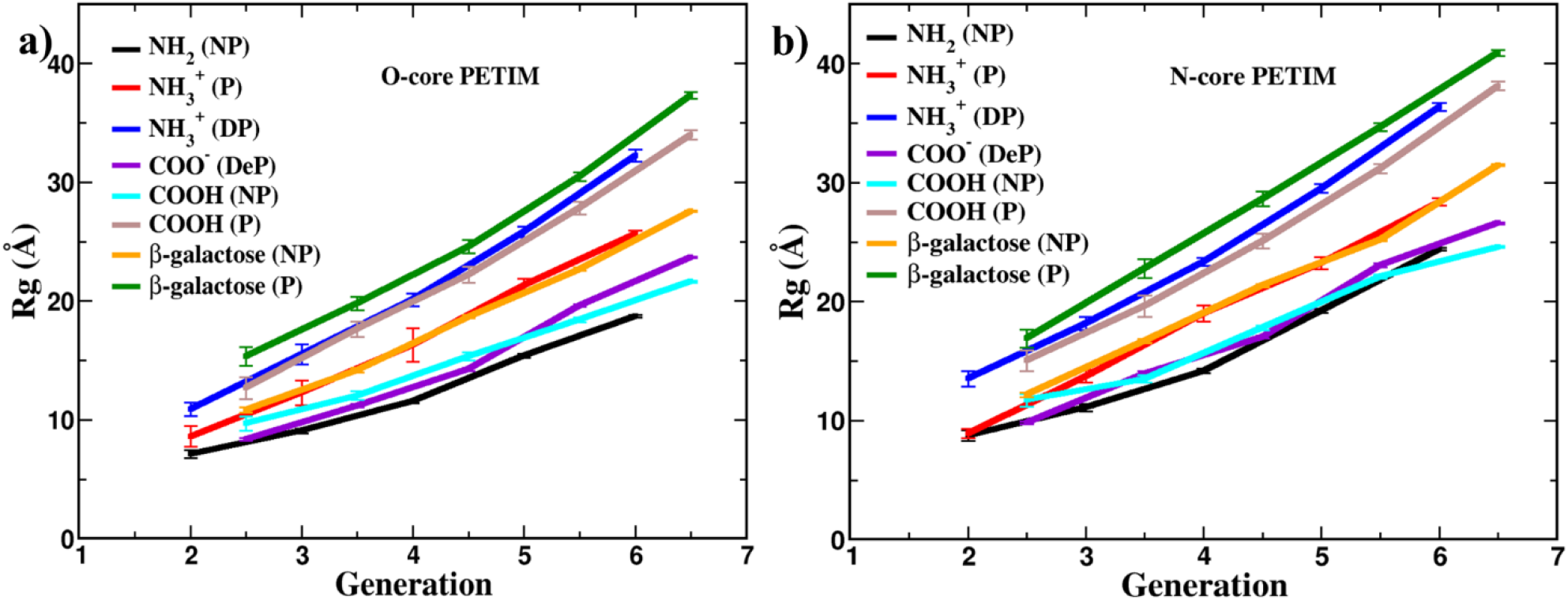
The radius of gyration (Rg) profiles of (a) O-core and (b) N-core PETIM dendrimers are presented as a function of dendrimer generation for different surface functionalizations. Comparison of the Rg values indicates that protonation has a substantially greater impact on the size of amine-terminated dendrimers than on carboxylic acid- or β-galactose-terminated dendrimers. In contrast, deprotonation of the carboxylic acid terminal groups results in only a marginal change in dendrimer size.

#### 4.1.2. Shape anisotropy of PETIM dendrimers

The asphericity and aspect ratios profiles of O-core and N-core PETIM dendrimers with various surface functionalisation were examined across generations (**Figures S4a** to **S4f**) to investigate the influence of surface chemistry on shape anisotropy. Comparative analysis highlights that the impact of protonation or charge on shape anisotropy is more pronounced for amine terminated dendrimers than for carboxylic or β-galactose terminated ones. Notably, dendrimers with a net neutral charge (-NH_2_/-COOH) display a more spherical morphology, characterized by lower asphericity and aspect ratios, relative to their protonated (-NH_3_^+^) or deprotonated (-COO^-^) counterparts, which exhibit higher asphericity and aspect ratios across generations. In protonated amine terminated dendrimers, the dendritic units elongate by clustering into two or three sub-clusters, thereby minimizing strong electrostatic repulsions between the positively charged terminal groups. This structural adaptation results in increased shape anisotropy compared to their non-protonation states. Furthermore, protonation of the tertiary amines causes the dendrimers to swell and become more porous because of electrostatic repulsions at the branch points. As a result, the double-protonated amine terminated dendrimers and the protonated β-galactose or carboxylic terminated dendrimers become more symmetric and exhibit higher radius of gyration (Rg) values than their non-protonated counterparts. Interestingly, the deviation from a spherical shape is more pronounced in intermediate generations (G3/G3.5 to G5/G5.5) than in smaller (G2/G2.5) or larger (G6/G6.5) dendrimers. In contrast, for deprotonated carboxylic dendrimers, the electrostatic repulsions between negatively charged -COO^-^ terminal groups are effectively mitigated by the screening effect of positively charged Na^+^ counter ions. Consequently, the impact of surface charge on shape anisotropy and size becomes less significant for carboxylic terminated dendrimers. Moreover, both protonated amine (-NH_3_^+^(P)) and non-protonated (NP) β-galactose terminated dendrimers exhibit similar Rg values; however, the NP β-galactose terminated dendrimers achieve a more symmetric and spherical shape compared to their -NH_3_^+^(P) terminated counterparts across the respective generations.

#### 4.1.3. Atomic arrangement inside PETIM dendrimers

The density profiles of O-core and N-core PETIM dendrimers (**Figures 3a** – **3p**) were computed as a function of radial distance from their centre of mass for various surface functionalizations to examine differences in their internal atomic arrangements. For all surface functionalizations, the terminal branch units of non-protonated (NP) dendrimers fold back into the interior counterparts due to favourable van der Waals (vdW) interactions. As a result, these dendrimers adopt more compact structures than their protonated or deprotonated counterparts. Consequently, the density profiles of NP dendrimers exhibit broader high-density regions near the centre of mass along with sharper tail regions. In contrast, the protonated (P) dendrimers display increased porosity and open conformations for amine, carboxylic and β-galactose terminated systems. This behaviour arises from strong electrostatic repulsions between the positively charged primary and tertiary amines. As a result, the width of the high-density regions decreases, leading to longer tail regions compared to the corresponding NP dendrimers. For deprotonated PETIM dendrimers terminated with COO^-^ groups, the electrostatic repulsions between negatively charged terminal groups are screened by oppositely charged Na^+^ counter ions. Consequently, deprotonation has only a minimal effect on the density distribution profiles of carboxylic terminated dendrimers compared to the neutral COOH-terminated (NP) dendrimers. This observation is consistent with the relatively small changes observed in the Rg, asphericity and density profiles of carboxylic terminated dendrimers in their COOH (NP) and COO^-^ (DeP) forms. Additionally, the width of the high-density and tail regions in the density distributions of protonated carboxylic (**Figures 3f** & **3n**) or β-galactose terminated dendrimers (**Figures 3h** & **3p**) are similar to those observed for double protonated amine-terminated dendrimers (**Figures 3c** & **3k**) of corresponding generations. This similarity arises because the density profiles are predominantly governed by the protonation of tertiary amines at the branch points, irrespective of the terminal functional groups. Furthermore, these observations suggest that the dendrimers possess comparable interior porosity and structural flexibility.

**Figure 3.**
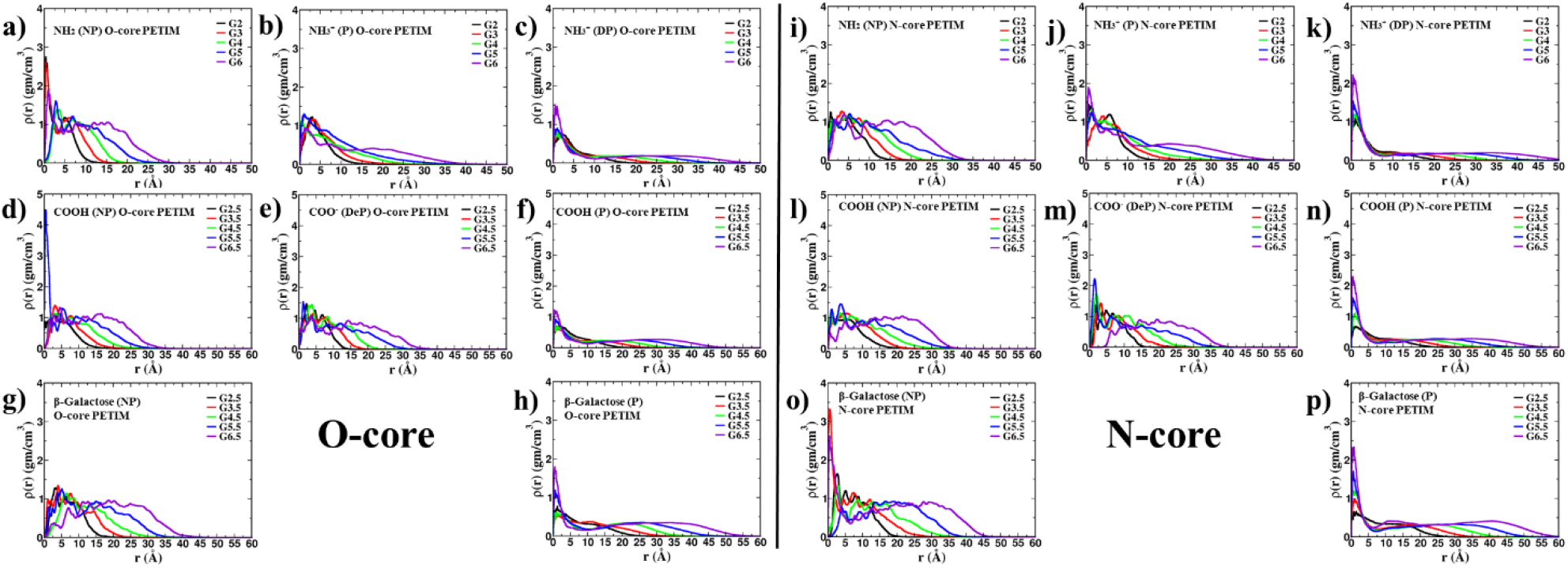
Radial density profiles are shown as a function of the distance from the centre of mass for PETIM dendrimers with an oxygen core (O-core; a–h) and a nitrogen core (N-core; i–p) bearing different surface functionalities. For amine-, carboxylic acid-, and β-galactose-terminated dendrimers, the protonated (P) forms exhibit lower central atomic densities and more extended tail regions than their non-protonated (NP) counterparts, indicating more expanded conformations. In contrast, deprotonation produces only minor changes in the radial density distributions of carboxylic acid-terminated dendrimers.

#### 4.1.4. Hydration of PETIM dendrimers

The quantities of inner, surface, and bulk bound water molecules (n_inner_, n_surf_, and n_bulk_) were calculated for both O-core (**Figures 4a** – **4c**) and N-core (**Figure 4d** – **4f**) PETIM dendrimers to investigate the hydration differences associated with various surface functionalizations. The hydration of the dendrimers increases monotonically with generation for all terminal groups. While the hydration differences are minimal in lower generation dendrimers, they become increasingly pronounced with increasing generation number. The bound water analysis reveals that β-galactose terminated dendrimers exhibit significantly higher hydration levels (n_inner_, n_surf_, and n_bulk_) compared to carboxylic and amine terminated dendrimers. This behaviour can be attributed to the larger size of the β-galactose functional groups and the presence of multiple hydroxyl groups, which facilitate the formation of a greater number of hydrogen bonds with surrounding water molecules. At low acidic pH, the tertiary amines of the dendrimer become protonated, leading to more open conformations with enhanced porosity for double protonated amine and protonated carboxylic and β-galactose terminated dendrimers (-NH_3_^+^(DP), -COOH (P), & β-galactose (P)). Consequently, these systems exhibit higher surface and bulk hydration (n_surf_, and n_bulk_) compared to the non-protonated dendrimers corresponding to basic pH conditions. In contrast, the non-protonated amine and carboxylic dendrimers (-NH_2_(NP) & -COOH (P)) exhibit the lowest hydration levels due to their comparatively compact conformations. Additionally, although electrostatic repulsions between the negative charged deprotonated carboxylic groups (-COO^-^ (DeP)) would generally favour expanded structures, these repulsions are effectively screened by Na^+^ counter ions. The stronger screening of COO⁻ groups can be attributed to favourable Na⁺-COO⁻ association arising from their matching hydration characteristics and localized electrostatic interactions ^47^. Molecular dynamics studies have reported pronounced Na⁺ association with carboxylate oxygens, whereas Cl⁻ association with NH₃⁺ groups is comparatively more diffuse ^48^. This localized Na⁺ association more effectively reduces COO⁻-COO⁻ electrostatic repulsion, thereby favouring a more compact dendrimer conformation. As a result, the hydration levels and compactness of deprotonated carboxylic dendrimer become comparable to those of the neutral non-protonated amine and protonated carboxylic-terminated dendrimers. Furthermore, hydration of charged biomolecules, such as DNA and dendrimers, can significantly influence their binding interactions, as demonstrated by previous experimental and computational studies ^49–52^.

**Figure 4.**
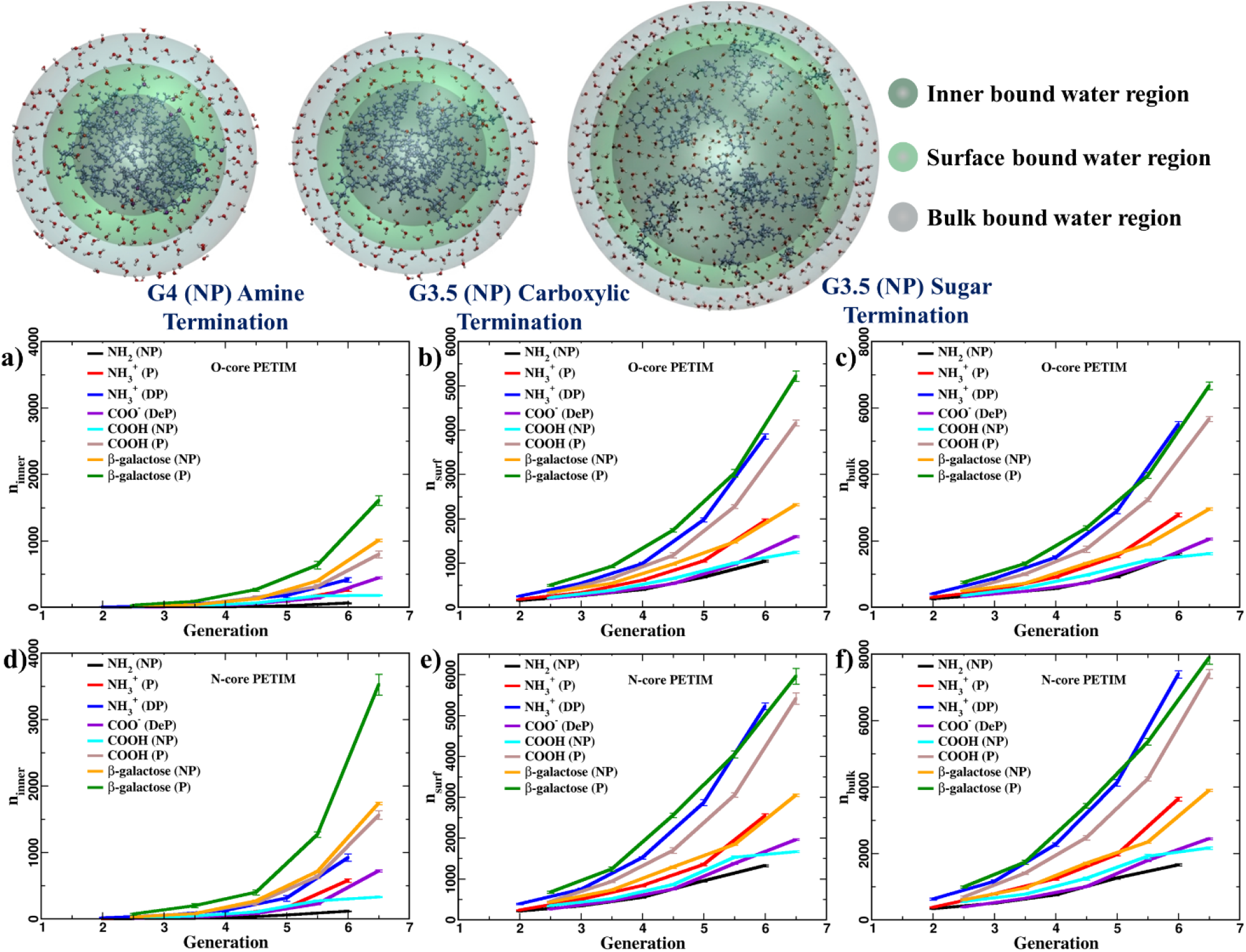
The numbers of inner (a, d), surface (b, e), and bulk (c, f) bound water (ninner, nsurf, & nbulk) are presented for O-core and N-core PETIM dendrimers with amine-, carboxylic acid-, and β-galactose-terminated surfaces as a function of dendrimer generation. The bound water analysis shows that β-galactose-terminated dendrimers consistently accommodate greater numbers of inner, surface, and bulk bound water molecules than their amine- and carboxylic acid-terminated counterparts.

#### 4.1.5. Internal flexibility of PETIM dendrimers

The root-mean-square-fluctuations (RMSFs) profiles of dendrimer branching points were analysed as a function of interior sub-generations (internal branching layers) for both O-core (**Figures 5a** – **5h**) and N-core (**Figures 5i** – **5p**) PETIM dendrimers to quantify their structural flexibility. The RMSF analysis reveals that the flexibility of the internal dendrimer layers decreases with increasing generation. Furthermore, dendrimer flexibility is strongly influenced by surface functionalization. For a given generation, amine terminated dendrimers exhibit the highest flexibility, whereas β-galactose terminated dendrimers show the lowest flexibility, with carboxylic terminated dendrimers displaying intermediate flexibility relative to their respective protonation states. The RMSF profiles of non-protonated (NP) dendrimers exhibit lower values than those of protonated (P) dendrimers, indicating greater rigidity and compactness in the NP systems. This behaviour arises from back-folding of dendrimer branching units toward the interior of the dendrimer. In contrast, the flexibility of charged protonated (P) and double protonated (DP) PETIM dendrimers terminated with amine, carboxylic, or β-galactose functional groups is enhanced by stronger electrostatic repulsions among their the positively charged primary and tertiary amines. In addition, the higher flexibility of amine-terminated dendrimers relative to β-galactose functionalized counterparts may be attributed to their comparatively lower water hydration. Conversely, deprotonation of surface carboxylic functional groups (-COO^-^ (DeP)) reduces dendrimer flexibility relative to the corresponding -COOH (NP & P) systems, primarily due to the charge screening effect by positively charged Na^+^ counter ions. Notably, the influence of protonation (P) or deprotonation (DeP) on dendrimer flexibility are more pronounced in lower generations (G2.5 – G4.5) than in higher generations (G5.5 – G6.5).

**Figure 5.**
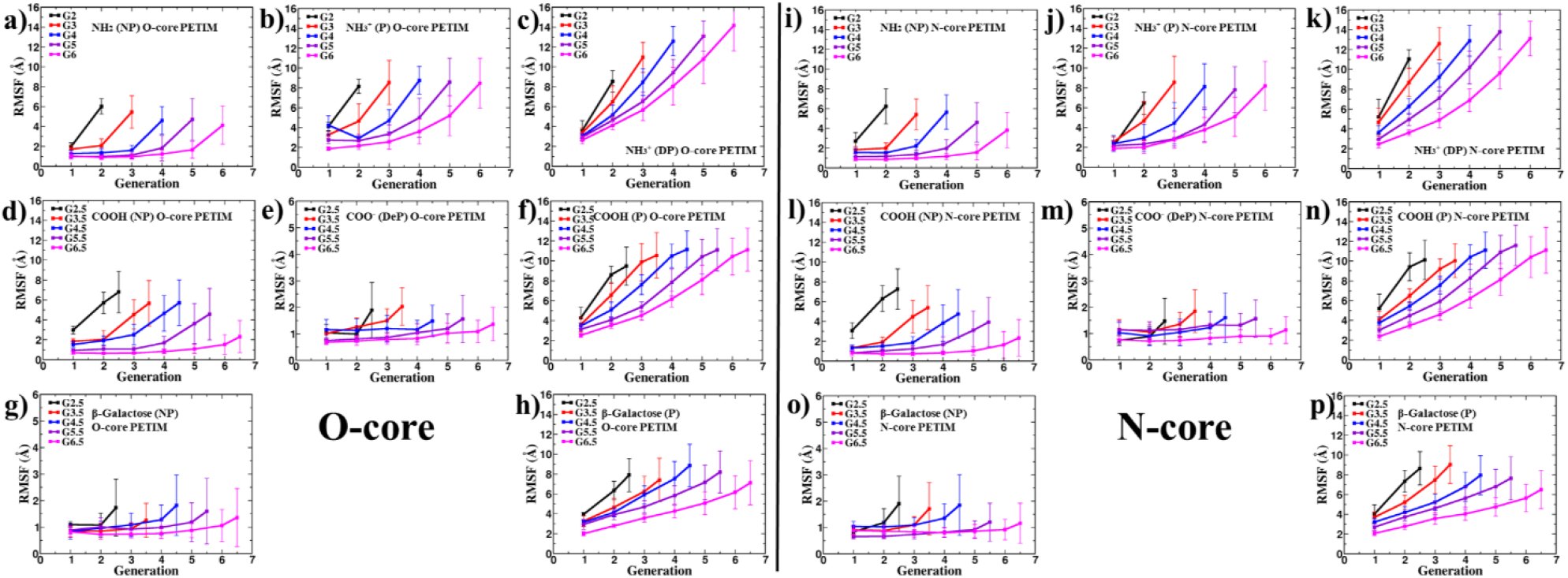
The root-mean-square fluctuation (RMSF) profiles of dendrimer branch points are presented as a function of interior sub-generation (internal branching layer) for O-core (a–h) and N-core (i–p) PETIM dendrimers with amine-, carboxylic acid-, and β-galactose-terminated surfaces. Deprotonation of the carboxylic acid terminal groups decreases the RMSF values, indicating reduced branch-point mobility. In contrast, protonation of the tertiary amines located at the branch points increases the RMSF values across all dendrimers, reflecting enhanced structural flexibility.

### 4.2. Comparison between oxygen core (O-core) and nitrogen core (N-core) PETIM dendrimers

The O-core and N-core PETIM dendrimers possess core multiplicities of two and three, respectively, resulting in architectures with two or three dendritic arms connected to oxygen or nitrogen core atoms. The additional arm in the N-core PETIM dendrimers facilitates a more compact packing of branching units compared to the O-core counterparts. Consequently, N-core dendrimers attain larger overall sizes while maintain greater internal compactness than the corresponding O-core dendrimers. As a result, the Rg values increase by approximately 12-16%, and the widths of the high-density regions in the radial density profiles increased by nearly 25-30%, accompanied by relatively larger tail regions for N-core dendrimers compared to O-core dendrimers of the same generation. Except for the NH_2_ (NP) and COO^-^ (DeP) terminated systems, the additional branch in N-core dendrimers reduces both shape anisotropy and flexibility compared to the corresponding O-core dendrimers. This behaviour is reflected by their lower asphericities and aspect ratios values and smaller RMSF values (**Figures S4** & **5**). In contrast, the NH_2_(NP) and COO^-^ (DeP) terminated N-core dendrimers exhibit enhanced anisotropy and flexibility compared to their O-core counterparts. This change arises from enhanced local domain formation in NH_2_ (NP) systems and the increased spatial separation between negatively charged COO^-^ groups in deprotonated (DeP) systems due to screening by Na^+^ counter ions. Furthermore, because N-core dendrimers possess 1.5 times more terminal groups than O-core dendrimers, they exhibit larger surface areas and greater water hydration capacity. Comparative analysis of bound water molecules (n_inner_, n_surf_, and n_bulk_) shows that N-core dendrimers accommodate approximately 1.5 times higher hydration than corresponding O-core dendrimers.

### 4.3. Structural, hydration, and conformational flexibility of PAMAM dendrimers

Polyamidoamine (PAMAM) dendrimers with an ethylene diamine (EDA) core are well-established for a wide range of applications ^27,53–56^. To elucidate the differences in the structural, hydration and conformational flexibility characteristics of PETIM dendrimers, a comparative study of their structural and dynamical properties relative to PAMAM dendrimers is essential. Accordingly, generations 1 to 5 of EDA-core PAMAM dendrimers were systematically investigated using all-atom molecular dynamics (MD) simulations (**Figure S5**) as a function of surface functionalization with amine (-NH_2_), carboxylic acid (-COOH), and D-glucoamine (-C_6_H_13_O_5_N) terminal groups under different protonation states (pH conditions).

The comparative analysis of the radius of gyration (Rg), asphericity, and aspect ratios reveals that the neutral NH_2_ (NP) and COOH (NP) terminated dendrimers adopt comparatively compact conformations with smaller sizes due to back folding of their dendritic units. In contrast, the positively charged NH_3_^+^ (DP), COOH (P), and D-glucose (P) terminated dendrimers exhibit more extended and porous conformations with larger sizes relative to the corresponding NP dendrimers, primarily due to electrostatic repulsions between the protonated tertiary amines located at the branch points (**Figure S6**). The observed pH-dependent changes in NH_2_ terminated dendrimer size and compactness are consistent with previously reported experimental and computational studies ^57,58^. Among these systems, the NH_3_^+^ (DP) terminated dendrimers display the highest degree of symmetry, with lower asphericity and aspect ratios values compared to the COOH (P) and D-glucose (P) terminated dendrimers (**Figure S7**). Furthermore, the NH_3_^+^ (P) and D-glucose (NP) terminated dendrimers exhibit intermediate sizes while maintaining similar shape anisotropies. This behavior is attributed to the comparatively larger size and higher hydroxyl group content of the D-glucose terminal moieties, which promote enhanced hydrogen bonding interactions with the surrounding water molecules relative to the NH_3_^+^ terminal groups. In contrast, deprotonation of the carboxyl terminal groups to form COO^-^ (DeP) dendrimers exerts only a marginal influence on dendrimer size relative to the corresponding COOH (NP) terminated systems. However, the COO^-^ (DeP) terminated dendrimers exhibit greater shape anisotropy, which arises from the increased spacing between the negatively charged COO^-^ terminal groups due to electrostatic screening by positively charged Na^+^ counter ions.

The effect of terminal functional groups on the internal structure of PAMAM dendrimers was investigated through the radial density distributions shown in **Figures S8a–S8h.** D-glucose terminated dendrimers exhibit a greater internal atom density, indicating a more compact architecture. In contrast, NH_2_-terminated dendrimers display lower internal densities, suggesting a comparatively expanded structure. The COOH terminated systems show intermediate packing characteristics, lying between the D-glucose and NH_2_ terminated dendrimers. Further, the protonation significantly alters the internal organization of PAMAM dendrimers regardless of surface functionalization. The protonated systems exhibit broader density distributions and reduced interior packing, reflecting the adoption of more open conformations. This structural expansion can be attributed to the electrostatic repulsion between positively charged primary and tertiary amine groups within the dendrimer framework, which drives the branches apart and decreases core compactness. For the COO⁻-terminated dendrimers, however, the associated Na⁺ counter ions screen these electrostatic repulsions. As a result, their density distributions remain comparable to those of the corresponding neutral COOH (NP) terminated dendrimers, with only minor differences in packing density.

To examine the effect of surface functionality on PAMAM dendrimer hydration, the inner, bound, or bulk bound water molecules were calculated with different terminal groups, as shown in **Figures S9a–S9c**. The highly porous NH_3_^+^ (DP), COOH (P) and D-glucose (P) terminated dendrimers accommodate a larger number of surface and bulk bound water molecules (**Figures 9b and 9c**), indicating enhanced solvent accessibility. Among them, D-glucose (P) terminated dendrimers exhibit the highest number of inner bound water molecules. The larger D-glucose terminal groups contain multiple hydroxyl functionalities that promote hydrogen bond formation with water molecules. In addition, the pronounced tail regions and deeper minima in their density profiles suggest enhanced surface hydration and the presence of internal cavities, facilitating water penetration into the dendrimer interior. As a result, a greater number of water molecules remains confined within the dendrimer structure. In contract, NH_3_^+^ (DP) terminated dendrimers adopt more expanded conformations with extended branches, as reflected by the broader low density tail regions in their density profiles. While this increases surface accessibility to water, it reduces the formation of the internal pockets, resulting in fewer inner bound water molecules (**Figure 9a**) compared with D-glucose terminated dendrimers.

To further assess the role of surface functionality in determining PAMAM dendrimer flexibility, RMSF profiles were evaluated for individual branch-point generations of dendrimers, as shown in **Figures S10a–S10h**. Among all systems, both NP and P of D-glucose terminated dendrimers display the smallest RMSF values, indicating a lower degree of structural fluctuation than the corresponding COOH and NH_2_ terminated dendrimers. The reduced fluctuations are consistent with the greater population of inner-bound water molecules, which provides additional stabilization to the dendrimer framework and enhances its structural rigidity. A significant increase in flexibility is observed upon protonation for all three surface functionalities. This behaviour arises from the repulsive interactions between positively charged amine groups, which promote branch expansion and increase local segmental motion. In contrast, the deprotonated COO^-^ terminated dendrimers exhibit comparatively lower flexibility. The presence of Na^+^ counter ions effectively moderates the electrostatic interactions associated with the COO^-^ terminal groups, resulting in a more constrained structure with reduced fluctuations.

### 4.4. Comparison between O-core PETIM and PAMAM dendrimers

Owing to the distinct core multiplicities of two dendrimer architectures, with the oxygen core PETIM exhibiting a functionality of two and the EDA core of PAMAM exhibiting a functionality of four, their branching growth follows different generation scaling. As a result, a PETIM dendrimer at generation *(G+1)* contains a comparable number of branch units to a PAMAM dendrimer at generation *G*. This equivalence provides a rational basis for comparing their structural and dynamical characteristics and for examining the influence of surface terminations across the two dendrimer families.

A comparison of global structural properties indicates that PETIM dendrimers generally adopt more expanded conformations than PAMAM dendrimers with a similar branching architecture, as reflected by their larger radius of gyration (Rg). The distinction is especially evident for dendrimers functionalized with β-galactose and D-glucose, whereas NH_2_ and COOH terminated systems exhibit comparatively smaller differences (**Figures S11a-S11c**). Moreover, the divergence between the two dendrimer families becomes more pronounced under protonated (P) conditions, while only modest variations are observed for the non-protonated (NP) and deprotonated (DeP) states. The trends in molecular shape, characterized through the asphericity and aspect ratio parameters (**Figures 6a-6i**), show that dendrimers of lower generations deviate more strongly form spherical symmetry. As generation number increases, both PETIM and PAMAM architectures gradually approach spherical symmetry. A notable exception is observed for NH_3_^+^ terminated protonated (P) systems, where the intermediate PETIM generations (G4 and G5) display markedly enhanced anisotropy relative to both the lower (G1-G2) and higher (G6) generations (**Figures 6a, 6d, and 6g**). This non-monotonic behaviour may originate from the uneven spatial distribution of internal branching segments, resulting in a less symmetric arrangement of subdomains within the PETIM framework compared with PAMAM dendrimers of analogous generation.

**Figure 6.**
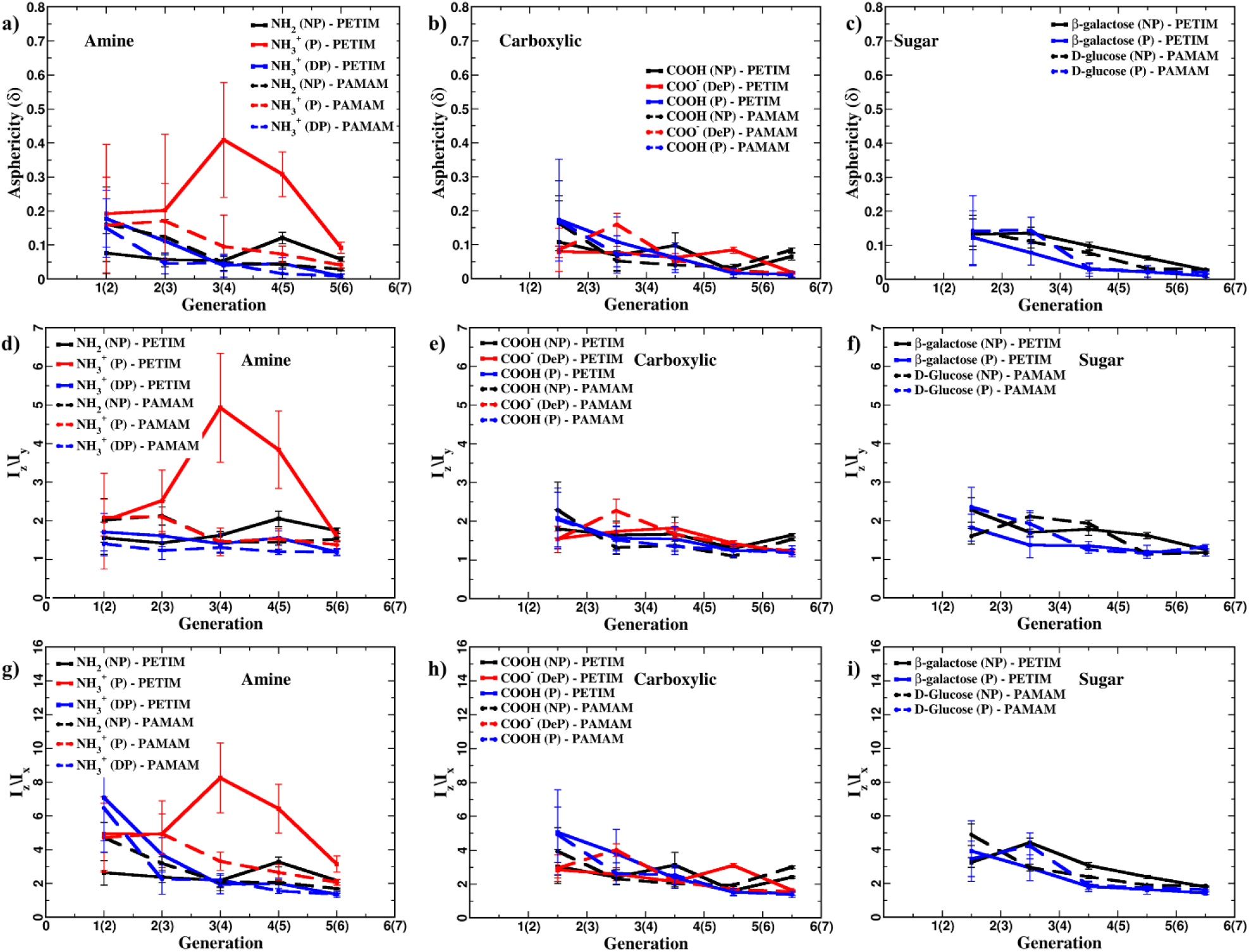
The asphericity and aspect ratio of O-core PETIM (solid lines) and PAMAM (dashed lines) dendrimers with (a, d, g) amine-, (b, e, h) carboxylic acid-, and (c, f, i) sugar (β-galactose/D-glucose)-terminated surfaces are compared to evaluate differences in shape anisotropy. The analysis demonstrates that NH₃⁺ (P)-terminated PAMAM dendrimers adopt more spherical conformations than the corresponding PETIM dendrimers with the same surface functionality, as reflected by their lower asphericity and aspect ratio values.

**Figure 7.**
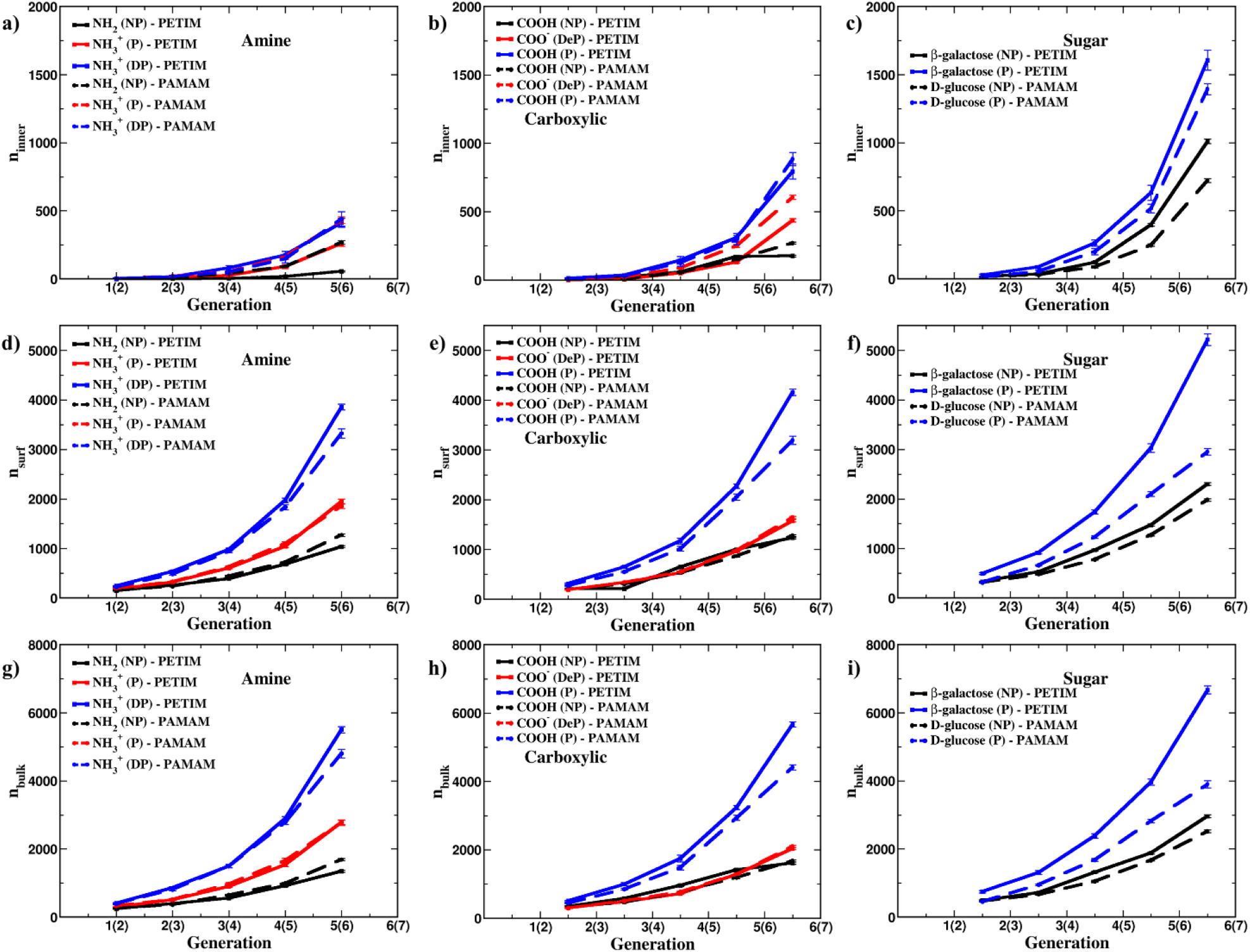
The numbers of inner-, surface-, and bulk-bound water molecules in O-core PETIM (solid lines) and PAMAM (dashed lines) dendrimers with (a, d, g) amine-, (b, e, h) carboxylic acid-, and (c, f, i) sugar (β-galactose/D-glucose)-terminated surfaces are compared to evaluate differences in their hydration behaviour. The results show that NH₂ (NP)-, NH₃⁺ (P)-, and COO⁻ (DeP)-terminated PAMAM dendrimers exhibit greater interior hydration than the corresponding PETIM dendrimers. In contrast, D-glucose-terminated PAMAM dendrimers display lower interior hydration than β-galactose-terminated PETIM dendrimers.

The internal organization of NH_2_ terminated dendrimers differs significantly between the two (PETIM and PAMAM) architectures. In the non-protonated state, PETIM dendrimers exhibit higher interior densities and fewer bound water molecules than equivalent PAMAM dendrimers, indicating a more compact structure **(Figures S12a, and S12i**). These differences, however, become less pronounced upon protonation. For COOH terminated systems, PETIM and PAMAM dendrimers display comparable internal densities and hydration levels under neutral conditions. Upon deprotonation (COO^-^), PETIM dendrimers become more internally accessible, resulting in lower packing density and greater water uptake than their PAMAM counterparts (**Figures S12e, and S12m**). Sugar-functionalized dendrimers exhibit a contrasting behaviour. The β-galactose terminal groups promote a more open peripheral arrangement in PETIM dendrimers than the D-glucose groups in PAMAM, leading to reduced interior density and enhanced hydration (**Figures S12g, S12h, S12o, and S12p, and 7c, 7f, 7i**). Consequently, PETIM dendrimers accommodate a larger number of bound water molecules. The RMSF analysis further reveals that NH_2_ and COOH terminated PETIM dendrimers are generally more flexible than the corresponding PAMAM dendrimers (**Figures S13a-S13f, and S13i-S13n**), although these differences diminish with protonation or deprotonation. In contrast, β-galactose terminated protonated PETIM dendrimers retain greater flexibility despite their higher hydration levels compared to D-glucose terminated PAMAM dendrimers (**Figures S13g, S13h, S13o, and S13p**). Furthermore, protonation exerts a stronger influence on both hydration and conformational dynamics in sugar terminated PETIM dendrimers, highlighting the greater sensitivity of this architecture to changes in the charge state.

### 4.5. Drug binding with PETIM dendrimers

The study was further extended to investigated the binding of drug molecules, namely neutral trans curcumin and positively charged doxorubicin, with O-core PETIM dendrimers possessing different surface functionalities (NH_2_, COOH, Sugar). The thermodynamically equilibrated structures of the drug-dendrimer complexes are shown in **Figures 8a and 8b**. Owing to its neutral nature, curcumin interacts with dendrimers primarily through van-der-Waal interactions, whereas the binding of positively charged doxorubicin is largely governed by electrostatic interactions. The MD simulations reveal that both curcumin and doxorubicin preferentially associate near the surface of NH_2_, COOH and β-galactose terminated dendrimers in their non-protonated (NP) states (**Figures S14a, S14d, S14i, and S14l**), reflecting the compact conformations adopted by these dendrimers. Upon protonation of amine groups or deprotonation of carboxylic groups, the dendrimer architectures become more open and porous, enabling curcumin molecules to penetrate deeper into the dendritic interior and enhance their van-der-Waal interactions (**Figure 9a**). In contrast, doxorubicin exhibits a distinct binding behaviour. The positively charged drug preferentially accumulates near the negatively COO^-^ groups of deprotonated dendrimers because of favourable electrostatic attractions (**Figure 9b**). However, protonation of primary and tertiary amines in NH_2_, COOH and β-galactose terminated dendrimers generates a net positive dendrimer charge, leading to strong electrostatic repulsion with doxorubicin and consequently reducing drug binding affinity. Furthermore, although protonated dendrimers adopt more expanded conformations, their lower internal densities provide fewer favourable interaction sites for curcumin, resulting in comparatively weaker binding than that observed for the corresponding NP dendrimers. These pronation dependent differences in dendrimer conformation and surface charge, and their impact on drug encapsulation, were quantitatively assessed using MMPBSA calculations (**Figures 9a and 9b**). The computed binding energies are consistent with the distinct binding modes and affinities observed for curcumin and doxorubicin across the various dendrimer systems.

**Figure 8:**
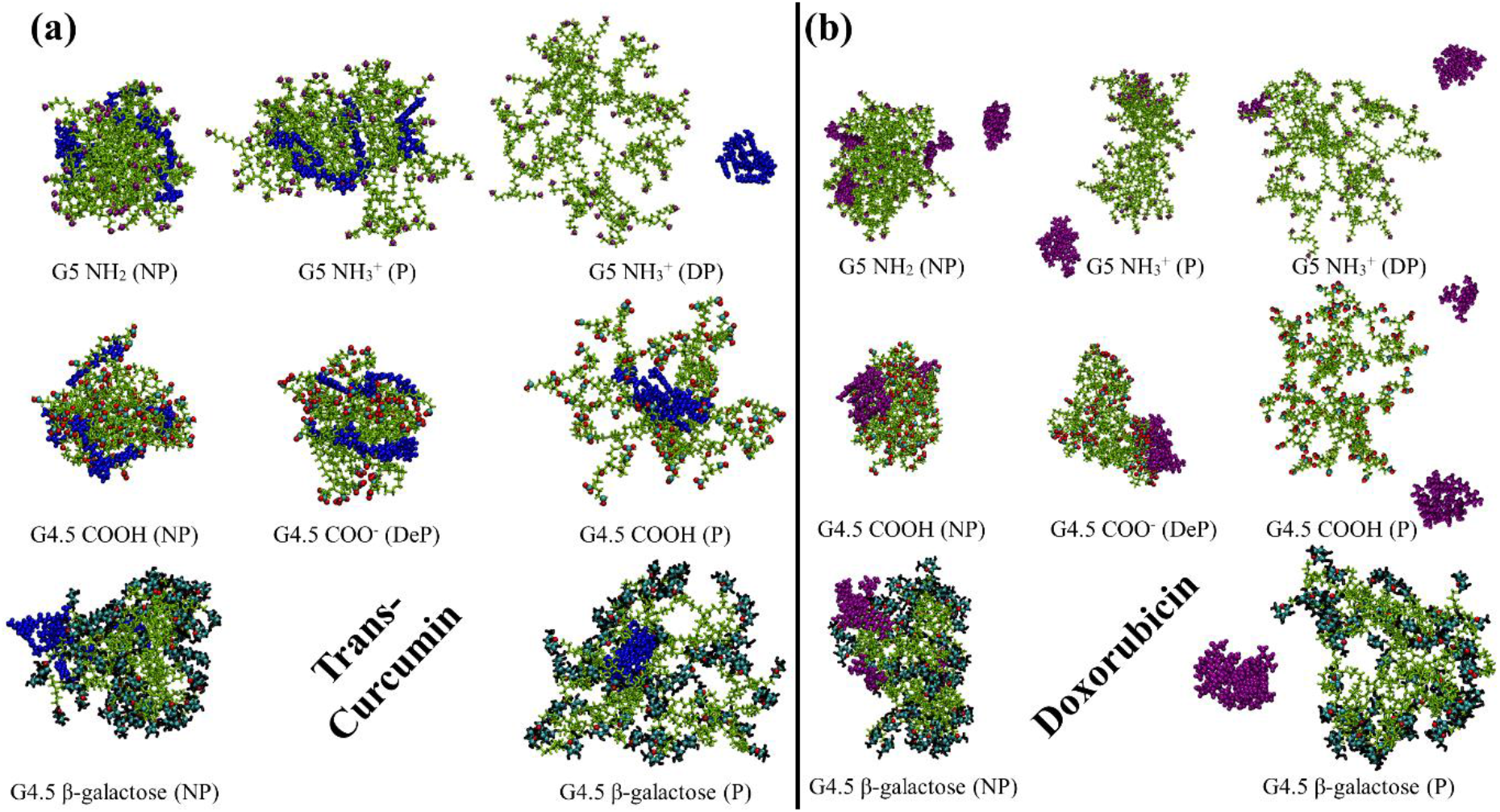
Final simulation snapshots of trans-curcumin (a) and doxorubicin (b) interacting with amine-terminated G5 and carboxylic-/sugar-terminated G4.5 O-core PETIM dendrimers under different pH conditions after 200 ns of molecular dynamics (MD) simulations. Except for the deprotonated carboxylic-terminated dendrimer, neutral trans-curcumin exhibits stronger binding to dendrimers with all terminal groups than doxorubicin. In contrast, the positively charged doxorubicin binds strongly to the negatively charged COO⁻ groups of the deprotonated carboxylic-terminated dendrimer due to favourable electrostatic interactions.

**Figure 9:**
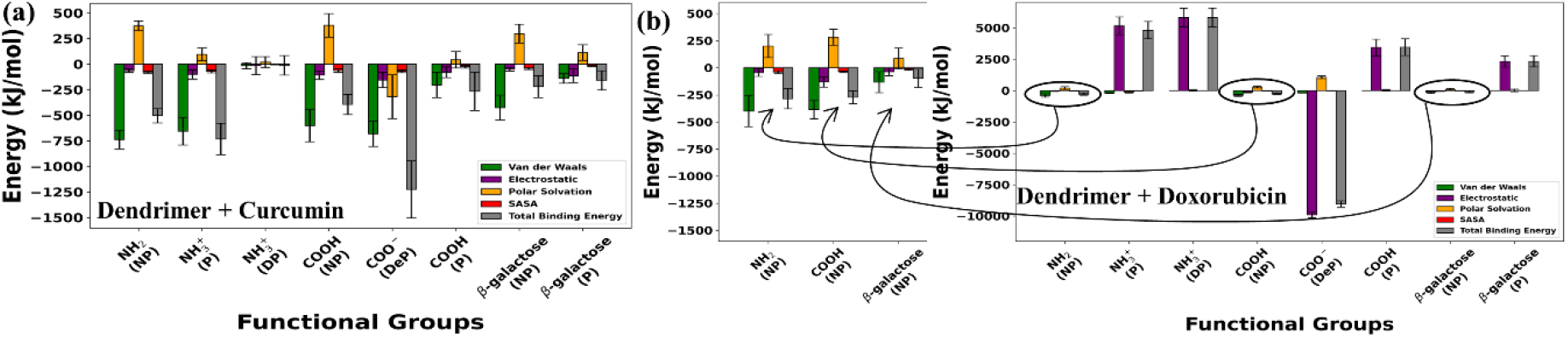
MM-PBSA binding free energies for trans-curcumin (a) and doxorubicin (b) interacting with O-core PETIM dendrimers possessing different terminal functionalities under various pH conditions. The deprotonated carboxylate-terminated dendrimer (DeP) exhibits the most favorable binding free energies for both drug molecules. For doxorubicin, electrostatic interactions dominate the binding, leading to strong affinity for the negatively charged COO⁻-terminated (DeP) dendrimer. In contrast, the binding of neutral trans-curcumin is governed primarily by van der Waals interactions, which constitute the major contribution to its overall binding affinity.

### 4.6. Spatial organization of drug molecules within functionalized PETIM dendrimers

The spatial distribution of drug molecules within the functionalized O-core PETIM dendrimers was examined by analysing the size and location of drug clusters formed during the simulations. The number of clusters containing different numbers of drug molecules was determined for curcumin and doxorubicin in the amine-, carboxylic- and β-galactose terminated dendrimers (**Figure 10 (a, c, e, g, i, k)**). This analysis distinguishes isolated drug molecules from groups of associated molecules and provides a measure of extent of drug clustering within the dendrimer cavity. To further characterize their spatial localization, the distance of each drug cluster from the dendrimer center was calculated (**Figure 10 (b, d, f, h, j, l)**). The resulting radial distribution indicates the preferred regions occupied by the drug molecules within the dendrimer-drug complexes.

**Figure 10:**
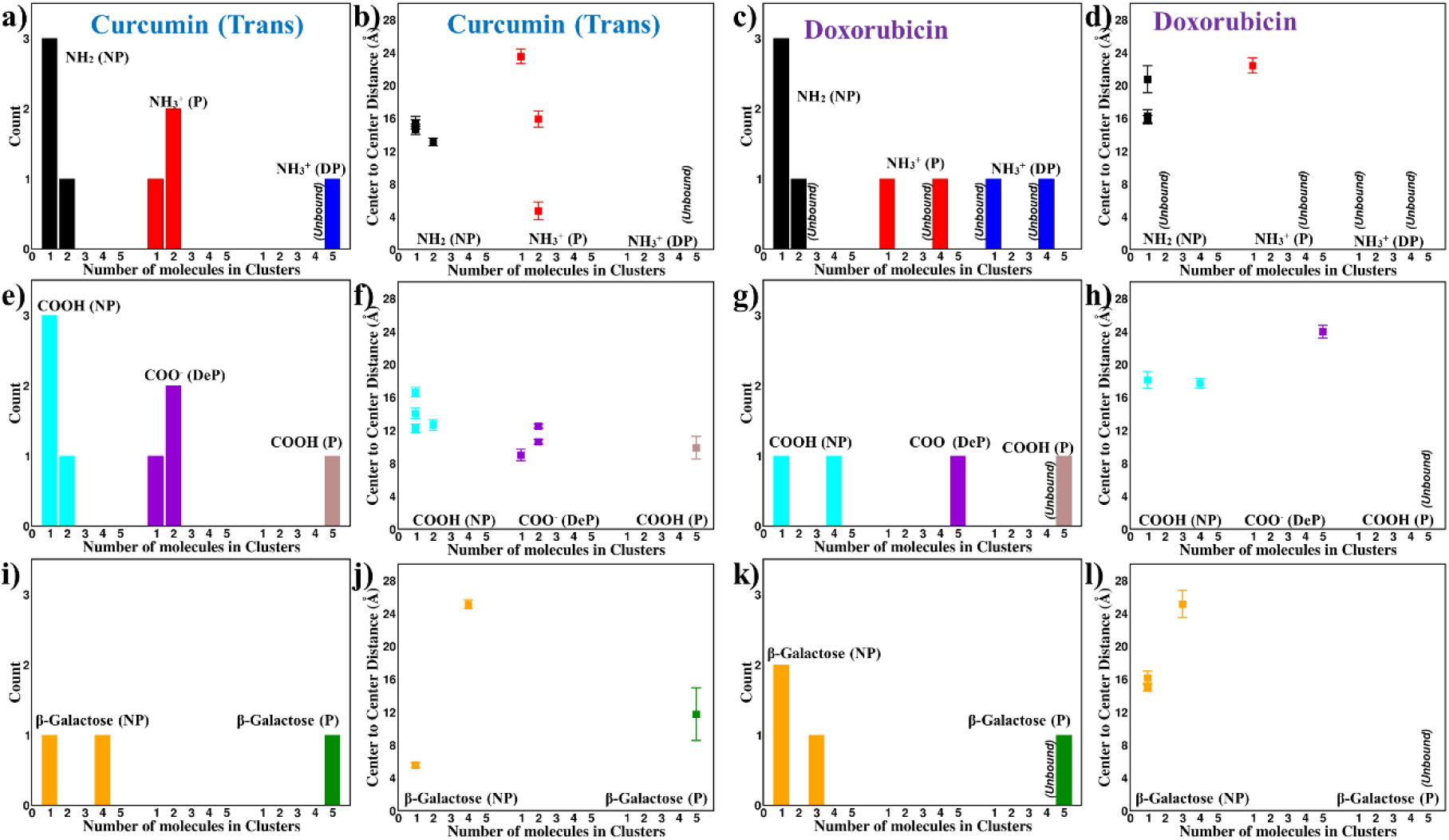
The number of clusters containing different numbers of drug molecules was calculated to characterize the spatial organization of the curcumin & doxorubicin within the (a, c) amine- (e, g) carboxylic- and (I, k) β-galactose-terminated dendrimers, respectively. The distance of each cluster from the dendrimer center was further analysed to determine the preferential binding locations of the drug molecules within the dendrimer-drug complexes (b, d: amine; f, h: carboxylic; and j, l: β-galactose). In the NP NH_2_ and COOH dendrimers, both drugs predominantly occur in a monomeric state, whereas drug clustering increases upon protonation of the primary and tertiary amines of the dendrimers. Curcumin clusters are preferentially located deeper within the dendrimer interior than doxorubicin clusters.

In the NP state, both NH_2_ and COOH terminated dendrimers exhibit a predominance of single-molecule clusters for both curcumin and doxorubicin, indicating that the drugs largely remain spatially separated rather than forming larger assemblies. Upon protonation of the dendrimer amine groups, an increase in the occurrence of the clusters containing multiple drug molecules is observed. This behaviour indicates that the protonation alters the local interaction environment within the dendrimer promotes greater association between the bounded drug molecules. Similar effects of dendrimer protonation on drug binding have also been reported in experimental studies using fluorescence spectroscopy and laser flash photolysis ^59^ and simulations ^60^. The extent of clustering differs with dendrimer surface functionality and drug type, highlighting the role of the dendrimer chemical environment in determining the organization of the encapsulated molecules. The radial distribution further reveals a clear difference in the localization of curcumin and doxorubicin. Curcumin clusters are predominantly located closer to the dendrimer center. They exhibit a larger number of contacts with the dendrimer and its terminal functional groups. In contrast, doxorubicin clusters are distributed at relatively larger radial distances from the dendrimer center. They show fewer contacts with the dendrimer functional groups, as shown in **Figure S15**. Thus, curcumin preferentially occupies the deeper regions of the dendrimer, while doxorubicin is more frequently distributed toward the outer regions. This difference in localization reflects the distinct molecular characteristics and interaction patterns of the two drugs within the heterogeneous dendrimer environment. Furthermore, the relatively lower SASA values of the unbound drug clusters compared with those of the dendrimer-bound drug clusters, as shown in **Figure S15 (c),** indicate increased exposure of the drug molecules to the solvent upon dendrimer binding. This suggests that dendrimer-drug association is accompanied by enhanced drug hydration. The effect of drug concentration on cluster formation was further examined by comparing the systems containing drug-to-dendrimer ratios of 5:1 and 10:1 **(Figure S16 and S17)**. Increasing the number of drug molecules results in a noticeable increase in drug clustering for both curcumin and doxorubicin. The enhanced clustering at the higher loading suggests that, as the number of drug molecules increases, drug-drug interactions become increasingly important in determining their organization within dendrimer. This effect is particularly evident when the available dendrimer binding environment becomes increasingly occupied, favouring association among neighbouring drug molecules. Overall, the cluster-size and radial-distribution analyses demonstrates that the spatial organization of curcumin and doxorubicin within O-core PETIM dendrimers is influenced by surface functionality, dendrimer protonation, drug loading, and molecular characteristics of the drug. These factors collectively determine both the extent of drug association and their preferred localization within the dendrimer-drug complexes.

### 4.7. Discussion

The structural properties and biomedical applications of dendrimers are strongly governed by pH-dependent variations in their size, shape, internal density, flexibility, and hydration. Under basic pH conditions, amine-terminated NH_2_ (NP) and carboxylic-terminated COO^-^ (DeP) dendrimers adopt compact, nearly spherical conformations characterised by high internal density, low flexibility, and limited hydration ^57,61,62^. These rigid architectures are advantageous for intravenous drug delivery because their compact size reduces hydrodynamic drag and immune clearance, thereby extending blood circulation time ^63–66^. Their dense interiors with limited flexibility also promote sustained drug release by increasing the diffusion path while protecting encapsulated therapeutics from enzymatic degradation and premature leakage ^53,54,67,68^. In addition, their compact structures can inhibit viral attachment through steric shielding of viral surface proteins ^69,70^. For carboxylic-terminated dendrimers, the combination of high rigidity and low water content further makes them attractive for nanoscale catalysis and organic electronic applications, where structural stability and minimal hydration are desirable ^65,71,72^.

At physiological (neutral pH) conditions, protonation of terminal amines to NH_3_^+^ (P) produces moderately flexible and hydrated dendrimers with increased asphericity arising from the formation of internal subdomains. These characteristics are favourable for oral drug delivery, where enhanced flexibility promotes mucoadhesion and improves intestinal permeation ^68,73,74^. The resulting architectures also suitable for tissue engineering owing to their improved structural adaptability ^75,76^. In contrast, carboxylic-terminated (COOH (NP)) dendrimers remain relatively compact and rigid at neutral pH, making them promising blood pool MRI contrast agents because of their stable size and low shape anisotropy. Their well-defined surface chemistry also supports bio-sensing and diagnostic applications ^62,71,77^. Sugar-terminated dendrimers, including β-galactose (NP) terminated PETIM and D-glucose (NP) terminated PAMAM, retain comparatively rigid structures with moderate hydration under both neutral and basic conditions. These properties are advantageous for MRI contrast enhancement by increasing rotational correlation time, while also supporting sustained drug release, nanoscale catalysis, and vaccine antigen presentation ^65,73,76^.

Under acidic pH conditions, protonation of the tertiary amines located at the dendrimer branch points induces substantial conformational expansion, resulting in increased size, internal porosity, flexibility, and hydration irrespective of surface functionality. These structural changes enhance the suitability of dendrimers for gene and siRNA delivery by facilitating nucleic acid encapsulation and intracellular transport through flexible frameworks and porous interiors ^53,54,67,68^.

The enlarged size and pH-responsive behaviour also favour controlled drug release in acidic tumour microenvironments, promote enhanced permeability and retention (EPR)-mediated tumour accumulation, and improve performance in hydrogel and tissue engineering applications ^27,64,78^. Furthermore, the highly hydrated and flexible conformations are beneficial for wound healing, pulmonary drug delivery, and antimicrobial therapies that rely on efficient interactions with biological membranes ^68,69,75,76^.

Further, the drug-loading performance of dendrimers is strongly governed by both their surface functionality and the physicochemical characteristics of the encapsulated drug. Among the systems investigated, neutral trans-curcumin and positively charged doxorubicin exhibit the strongest binding affinities toward deprotonated carboxylic-terminated (COO^-^) dendrimers under basic pH conditions, indicating that this protonation state is the most favourable for drug loading. Upon exposure to acidic environments, such as tumour microenvironment or intracellular endosomal compartments, protonation of the dendrimer weakens the drug-dendrimer interactions, thereby facilitating pH-responsive and sustained drug release. This behaviour is particularly advantageous for targeted drug-delivery applications, as it promotes drug retention during circulation while enabling controlled release at the desired site of action. The superior binding exhibited by carboxylic-terminated dendrimers toward both trans-curcumin and doxorubicin further underscores the critical role of surface charge engineering in optimizing dendrimer-based nano-carriers. Overall, these findings provide molecular-level design principles for tailoring PETIM and PAMAM dendrimers to achieve enhanced drug encapsulation efficiency and controlled, stimulus-responsive drug release.

## 5. Conclusions

In this study, we systematically investigated the structural properties of O-core and N-core PETIM dendrimers together with PAMAM dendrimers terminated with amine, carboxylic and sugar surface functional groups under different protonation states corresponding to varying pH conditions. Their drug encapsulation behaviour toward neutral trans-curcumin and positively charged doxorubicin was also examined. The molecular dynamics (MD) simulations demonstrate that protonation of the tertiary amines located at the dendrimer branch points results in expanded conformations with increased size and internal porosity, and hydration irrespective of surface functionality. In contrast, the neutral non-protonated NH_2_ and COOH terminated dendrimers and the negatively charged deprotonated COO^-^ terminated dendrimers adopt more compact conformations with reduced dimensions. Protonation of terminal amines (NH_3_^+^), corresponding to neutral pH, produces the greatest shape anisotropy in both PETIM and PAMAM dendrimers. The influence of protonation on dendrimer size, shape, and hydration is considerably stronger for amine- and sugar-terminated dendrimers, whereas deprotonation has only a modest effect on carboxylic-terminated (COO^-^) systems. Hydration analysis further reveals that sugar terminated dendrimers (β-galactose terminated PETIM and D-glucose terminated PAMAM) are the most hydrophilic among all surface functionalities. In addition, PAMAM dendrimers terminated with NH_2_ (NP), NH_3_^+^ (P), and COO^-^ (DeP) exhibit greater hydration than the corresponding PETIM dendrimers, whereas D-glucose terminated PAMAM dendrimers are comparatively less hydrophilic than β-galactose terminated PETIM dendrimers. Flexibility (RMSF) analysis shows that amine terminated dendrimers possess the highest structural mobility, sugar terminated dendrimers are the most rigid, and carboxylic-terminated dendrimers exhibit intermediate flexibility. Protonation consistently enhances dendrimer flexibility, whereas deprotonation of carboxylic-terminal groups reduces the structural fluctuations.

Molecular dynamics simulations of dendrimer-drug complexes indicate that, except for the COO^-^ terminated systems, neutral trans-curcumin exhibits stronger binding affinity than positively charged doxorubicin. MMPBSA free-energy calculations reveal that dendrimer-curcumin complexation is dominated by van-der-Waal (vdW) interactions, whereas dendrimer-doxorubicin binding is primarily driven by electrostatic interactions. Among the different surface functionalities, NH_2_ (NP), NH_3_^+^ (P), COOH (NP) and COO^-^ (DeP) terminations provide the most favourable drug-binding characteristics. Furthermore, the spatial organization of curcumin and doxorubicin within O-core PETIM dendrimers is governed by dendrimer surface functionality, protonation state, drug loading, and drug-specific interactions. Protonation and higher drug loading promote drug-drug clustering, while curcumin preferentially localizes toward the dendrimer interior and doxorubicin toward the outer regions. Overall, this work establishes clear relationships between dendrimer architecture, surface functionality, protonation state, and drug-binding behaviour. These molecular-level insights provide useful design principles for selecting appropriate dendrimer surface functionalities and pH conditions to optimize drug loading, stability, and delivery performance in dendrimer-based nano-carrier systems.

**Table 1:**
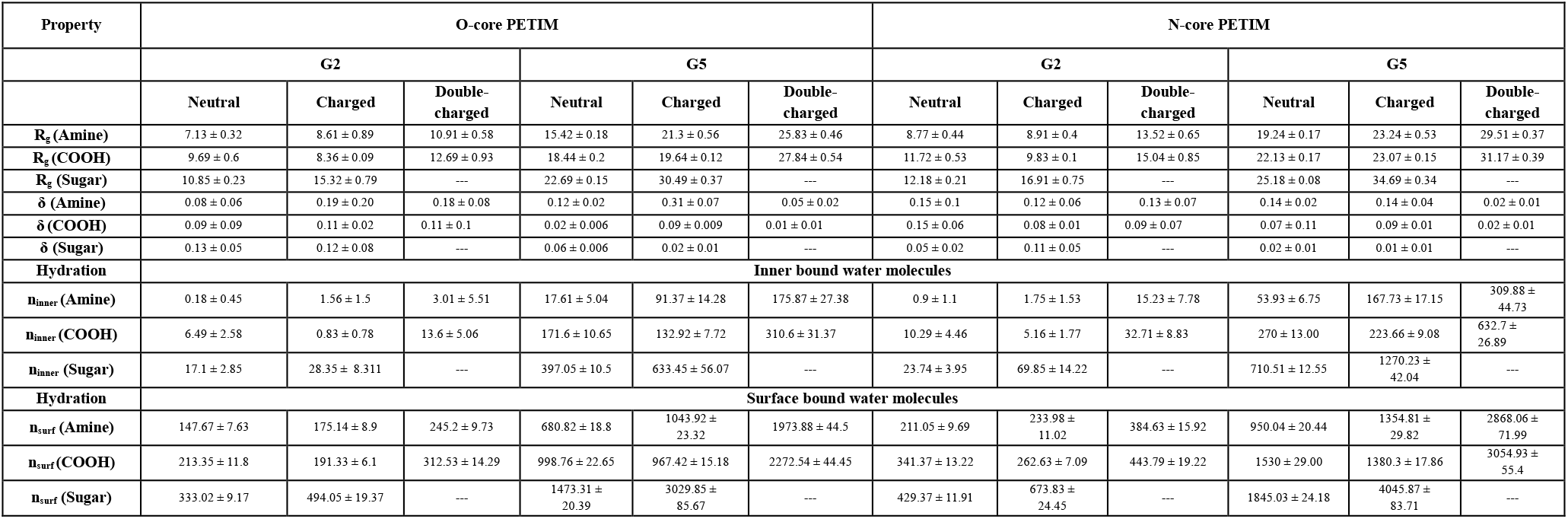
Comparison of structural parameters of oxygen core (O-core) and nitrogen core (N-core) PETIM dendrimers.

**Table 2:**
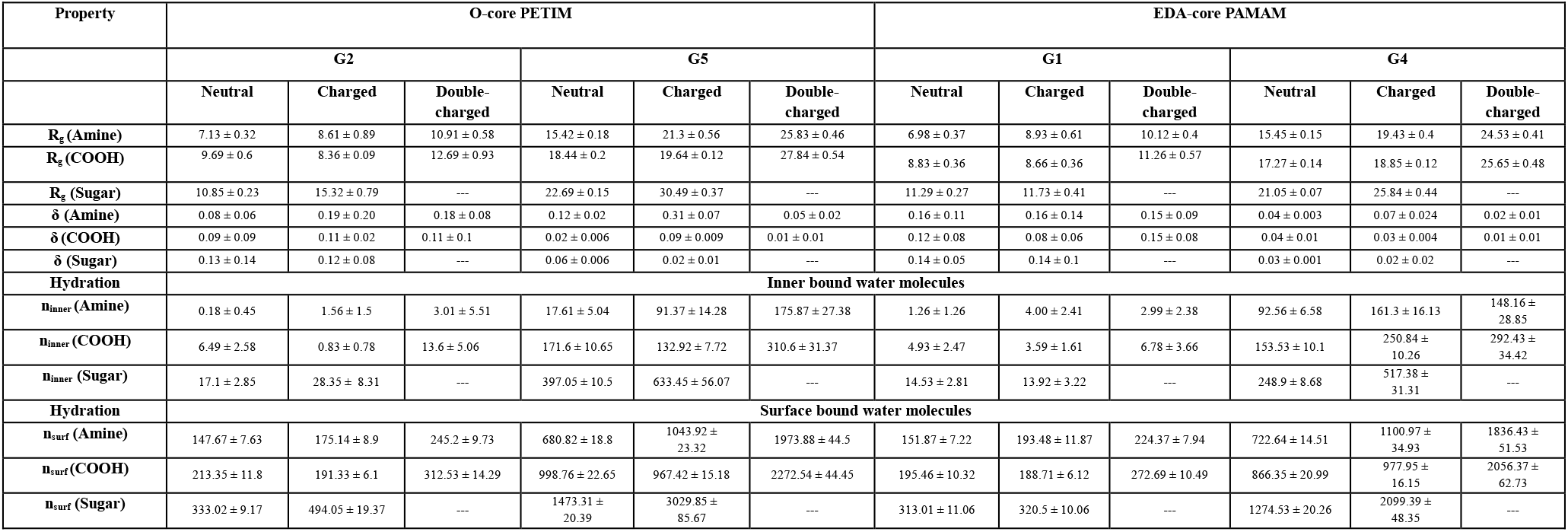
Comparison of structural parameters of PETIM and PAMAM dendrimers.

**Table 3:** Structural properties comparison for PETIM (O-core and N-core) dendrimers.

| Properties | NH<sub>2</sub> (NP)<br>Basic | NH<sub>3</sub><sup>+</sup> (P)<br>Neutral | NH<sub>3</sub><sup>+</sup> (DP)<br>Acidic | COO<sup>-</sup> (DeP)<br>Basic | COOH (NP)<br>Neutral | COOH (P)<br>Acidic | Galac (NP)<br>Neutral/Basic | Galac (P)<br>Acidic |
| --- | --- | --- | --- | --- | --- | --- | --- | --- |
| Size | Low | Medium | High | Low | Low | High | Medium | High |
| Spherical | High | Low | Medium | High | High | Medium | Medium | High |
| Density | High | Medium | Low | High | High | Low | High | Low |
| Flexibility | Low | Medium | High | Very Low | Low | High | Very Low | Medium |
| Bound water | Low | Medium | High | Low | Low | High | Medium | High |

## Supporting information

supporting_information_dend_paper

## Author contributions

A.G. conceived the study, performed the MD simulations, and analysed the data. S.K. conceptualized and designed the research. A.G., S.M., and S.K. contributed to discussions, data compilation, and the writing and revision of the manuscript. All authors reviewed and approved the final version of the manuscript.

## Supporting Information

The Supporting Information including residue information, 2D-representations of dendrimer architecture, Rg time profiles, asphericities of PETIM, MD snapshots, Rg, asphericities, density profiles, bound water, and RMSFs of PAMAM, comparison of Rg, density, RMSF profile of PAMAM & PETIM, density profiles and analysis of dendrimer-drug interactions, Effect of concentration on curcumin/doxorubicin binding with dendrimers, Tables containing atomic charges, force field parameters of dendrimers with various functionalization are available free of charge on the ACS Publications website at DOI: XXXXXX. Further, the 100 ns equilibrated MD structures of the dendrimers with all investigated surface functional groups are provided in the accompanying data repository link https://github.com/anujgarg939/petim-pamam-surface-functionality-ph-md.

## Acknowledgements

The authors thank the SAI-HPC, COSMOS Lab (CRIF) and DMACS at Sri Sathya Sai Institute of Higher Learning for providing the necessary computational resources. The authors also acknowledge the assistance of AI large-language-model tools in improving the grammar, clarity, and readability of the manuscript. All scientific content, analysis, and conclusions are solely the responsibility of the authors. The authors are grateful to Bhagawan Sri Sathya Sai Baba, Founder Chancellor, SSSIHL for his constant inspiration.

## TOC Graphic

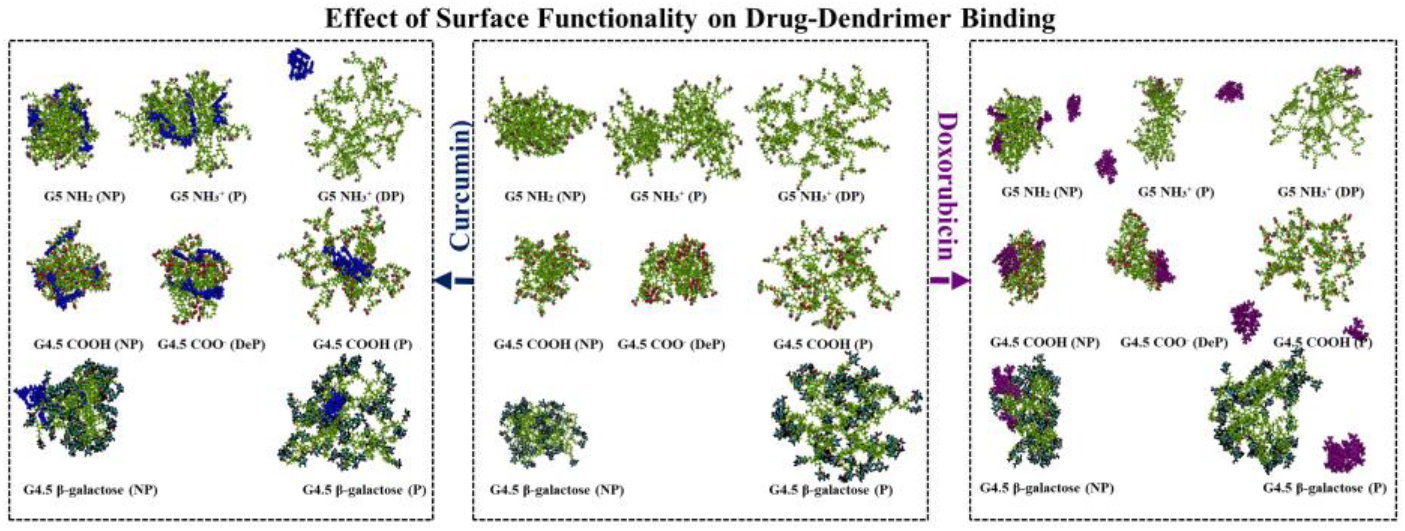

