## supporting_information_dend_paper for "Surface Functionality and pH Govern Structural Dynamics and Drug Binding in PETIM and PAMAM Dendrimers"

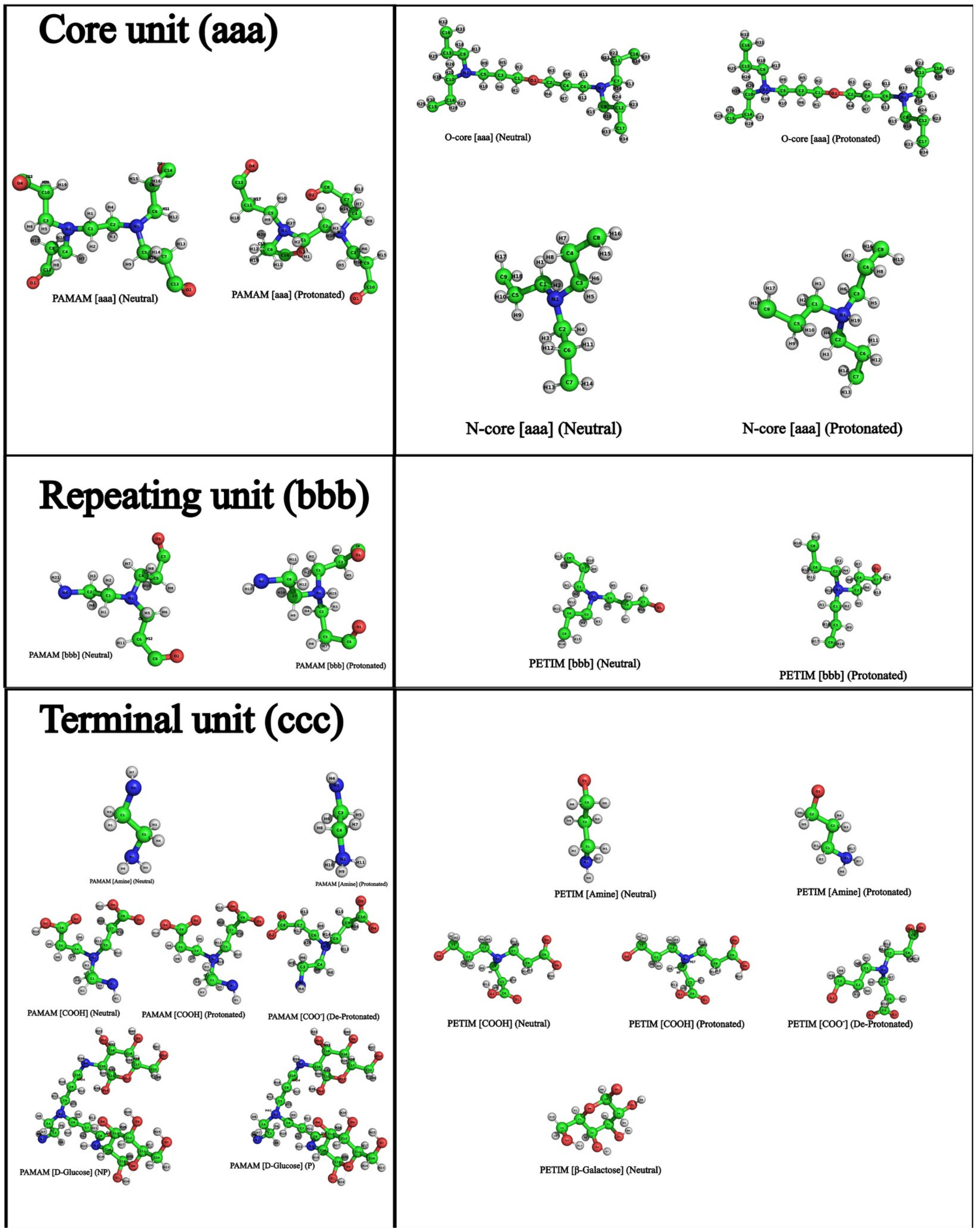

**Figure S1:** Residue information involved in dendrimer building.

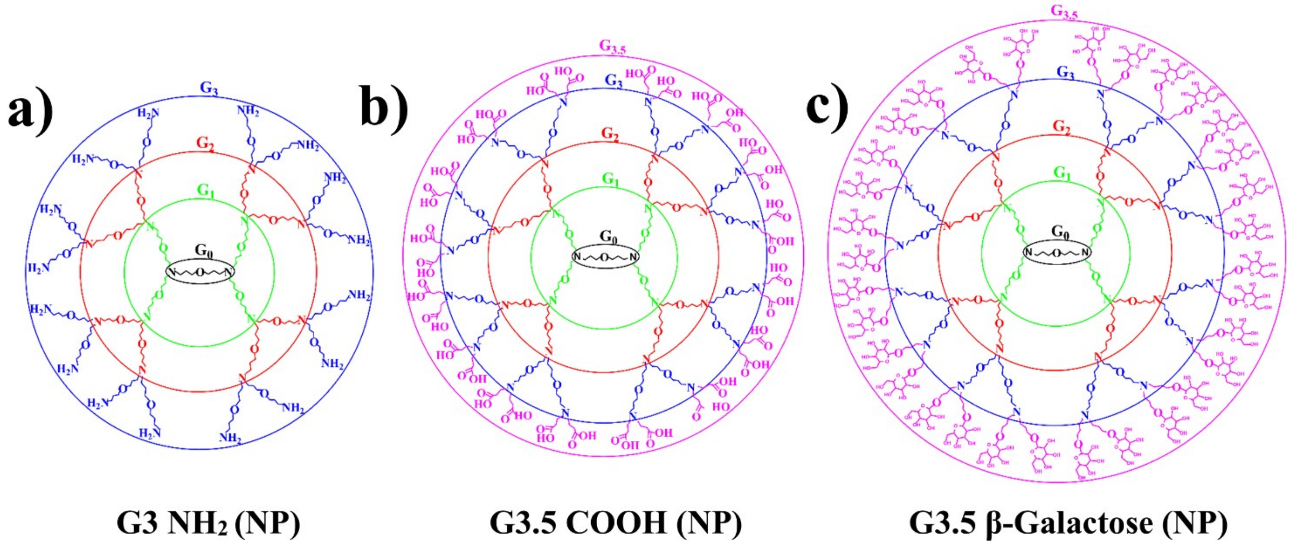

**Figure S2:** 2-D representations of dendrimer architectures with different surface functionalizations: (a) G3 $NH_2$, (b) G3.5 COOH, and (c) G3.5 β-galactose terminated dendrimers, illustrating their differences in surface chemistry and terminal functional groups. The G3 $NH_2$ dendrimer contains 16 terminal $NH_2$ groups, whereas the half-generation G3.5 COOH and G3.5 β-galactose terminated dendrimers each contain 32 terminal functional groups.

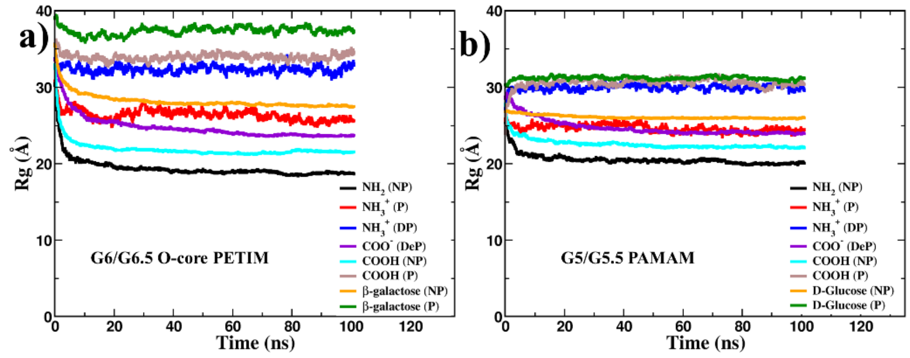

**Figure S3:** Time evolution of the radius of gyration ($R_g$) for (a) G6/G6.5 O-core PETIM dendrimers, and (b) G5/G5.5 PAMAM dendrimers over a 100 ns MD simulation. In the initial phase, dendrimers rearrange to the solvent environment and equilibrate by about 40 ns, after which $R_g$ values saturate.

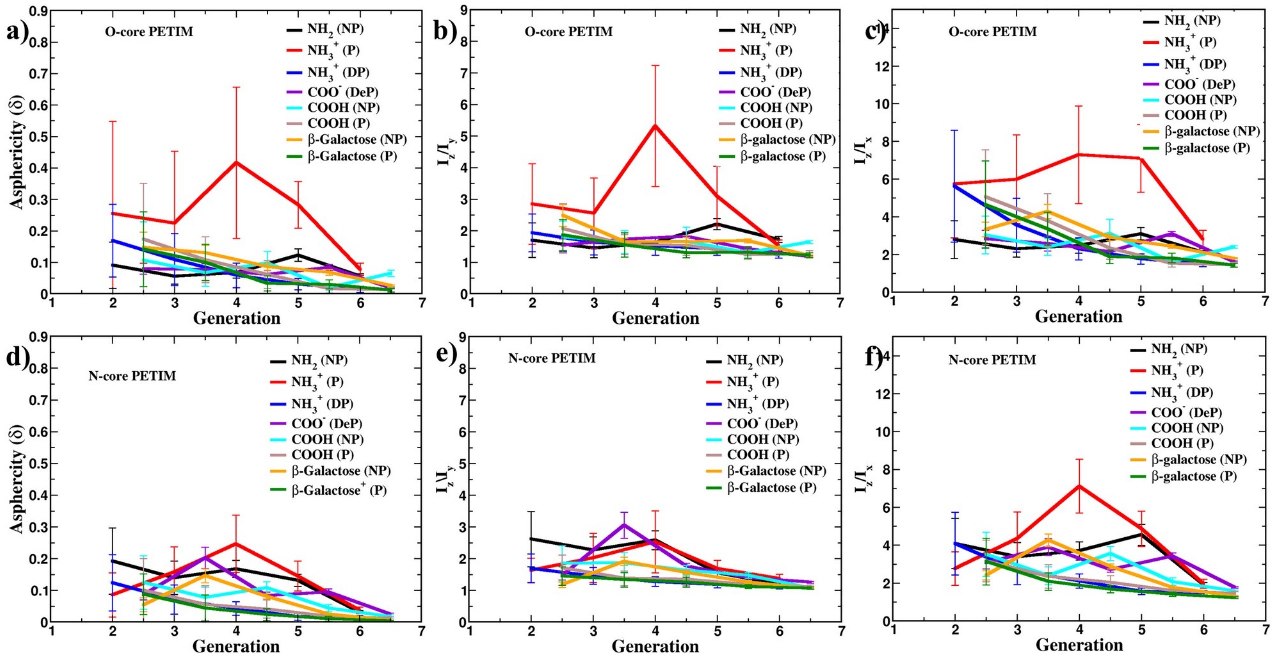

**Figure S4.** The profiles of asphericities and aspect ratios for oxygen core (O-core) (a-c) and nitrogen core (N-core) (d-f) dendrimers are computed across generations for dendrimers with amine, carboxylic and β-galactose terminal functional groups. The effect of protonation of the surface primary amines ($-NH_3^+$, P) on dendrimer shape anisotropy is more pronounced than that of carboxylic or β-galactose terminated dendrimers.

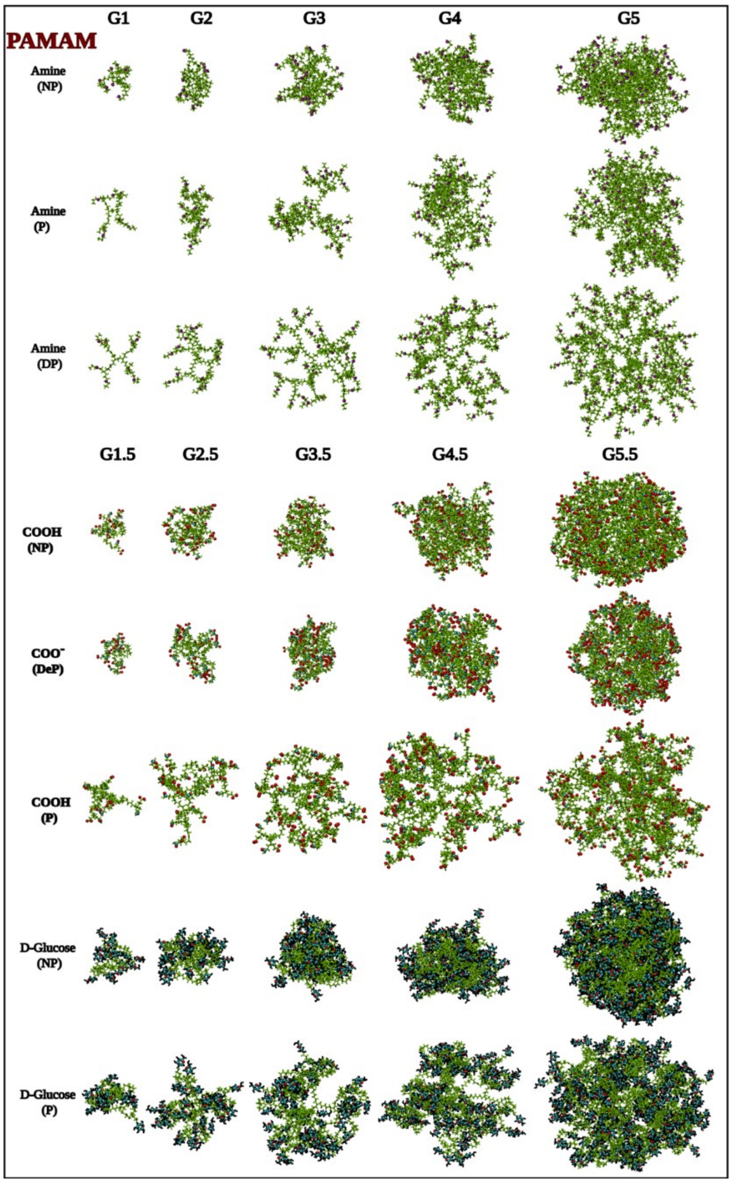

**Figure S5.** Final simulation snapshots of $NH_2$ (NP), $NH_3^+$ (P), $NH_3^+$ (DP), COOH (NP), $COO^-$ (DeP), COOH (P), D-Glucose (P) and D-Glucose (NP) terminated (G1-G5) PAMAM dendrimers after 100ns MD. Protonation of the tertiary amines located at the dendrimer branch points results in more extended conformations and greater internal porosity for $NH_3^+$ (DP), COOH (P), and D-Glucose (P) surface functionalised dendrimers than for the corresponding NP and DeP dendrimers. This behaviour arises from the strong electrostatic repulsion between the positively charged tertiary amines.

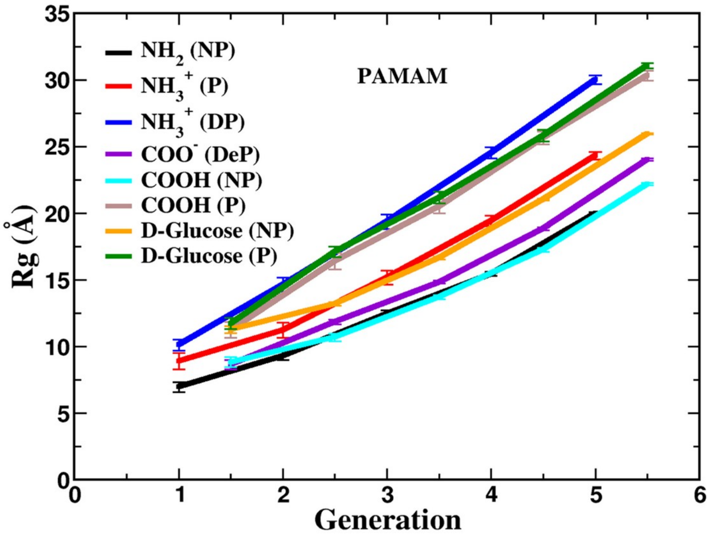

**Figure S6.** The radius of gyration (Rg) profiles of the PAMAM dendrimers, as a function of generation, are compared to understand the effect of surface functionalization on their size. The results reveal that the $NH_3^+$ (DP), COOH (P) and D-Glucose (P) terminated PAMAM dendrimers exhibit larger sizes than their $NH_2$ (NP), COOH (NP), and $COO^-$ (DeP) counterparts.

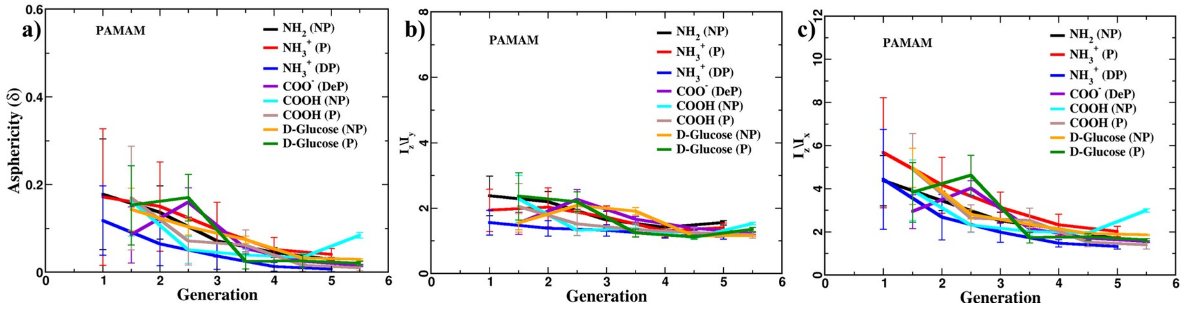

**Figure S7.** The asphericity and aspect ratios of PAMAM dendrimers with various surface terminations are plotted as a function of generation. The results reveal that the influence of protonation is substantially more pronounced in amine terminated PAMAM dendrimers than in carboxylic or D-Glucose terminated dendrimers. Furthermore, $NH_3^+$ (DP) and COOH (NP) terminated dendrimers exhibit more isotropic conformations than their respective counterparts.

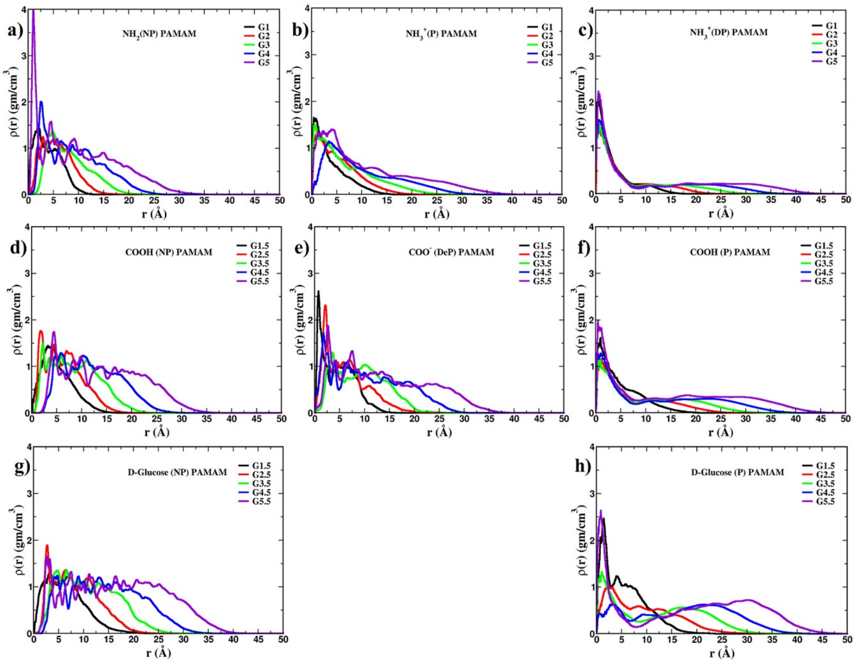

**Figure S8.** The radial density profile of PAMAM dendrimers are plotted as a function of distance from the centre of mass for various surface functional terminations. The analysis demonstrates that the non-protonated carboxylic/D-Glucose terminated dendrimers exhibit broader high-density regions near the centre of mass than similarly sized amine terminated dendrimers. In contrast, the reduced high-density regions observed in P and DP amine terminated PAMAM dendrimers indicate increased internal porosity relative to their NP counterparts.

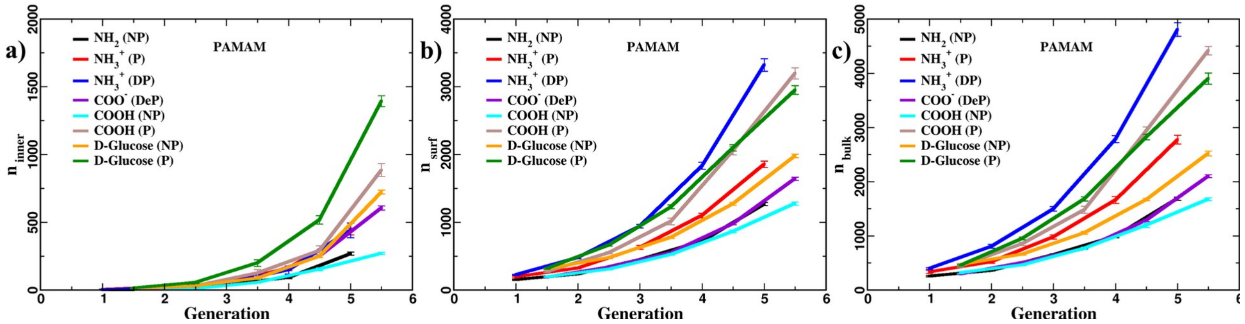

**Figure S9.** The inner ($n_{inner}$), surface ($n_{surf}$) and bulk ($n_{bulk}$) bound water content of PAMAM dendrimers, with various terminal groups ($NH_2$, COOH, and D-Glucose) and corresponding to different protonation states, is calculated as a function of dendrimer size (Generation). The analysis reveals that $COO^-$ (DeP), COOH (NP), and $NH_2$ (NP) terminated dendrimers accommodate fewer bound water molecules than D-Glucose (P), $NH_3^+$ (P), and $NH_3^+$ (DP) terminal dendrimers. This difference arises from the back-folding of the terminal branches in $COO^-$ (DeP), COOH (NP), and $NH_2$ (NP) terminated dendrimers, which restricts water penetration, whereas the outward extension of the terminal braches in D-Glucose, $NH3^+$ (P), and $NH_3^+$ (DP) terminated dendrimers facilitates greater water binding.

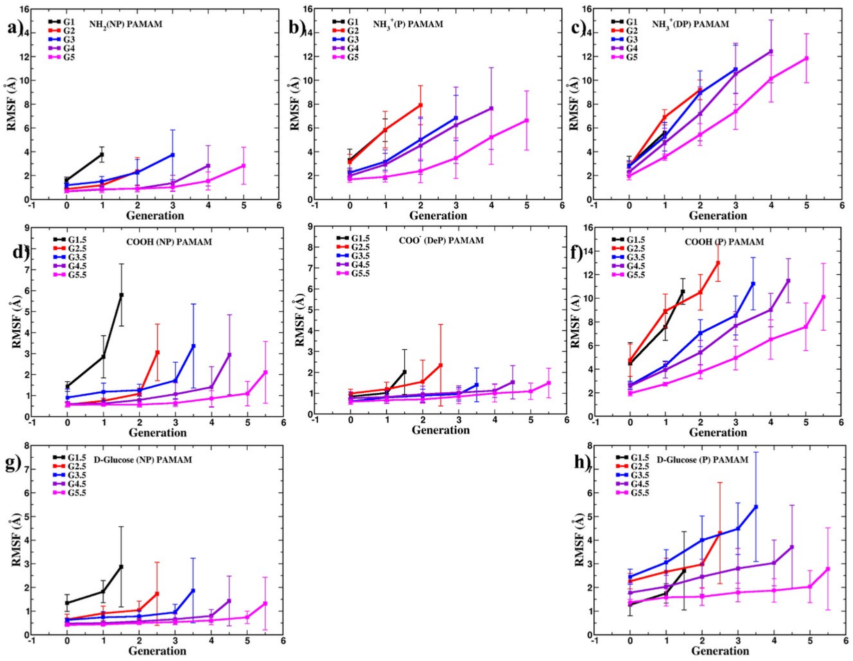

**Figure S10.** The RMSF profiles of branching points in PAMAM dendrimers are calculated for each generation, considering their internal sub-generations. The analysis reveals that amine terminated dendrimers exhibit higher RMSF values than carboxylic/D-Glucose terminated dendrimers, indicating greater structural flexibility. Additionally, the RMSF of amine terminated dendrimers increases with protonation ($RMSF_{DP}$ > $RMSF_{P}$ > $RMSF_{NP}$), owning to progressively stronger electrostatic repulsion among the positively charged primary and tertiary amines in the P and DP states, respectively. In contrast, deprotonation of carboxylic terminal groups reduces the RMSF because the electrostatic repulsion between negatively charged $COO^-$ groups is effectively screened by the associated positively charged counterions.

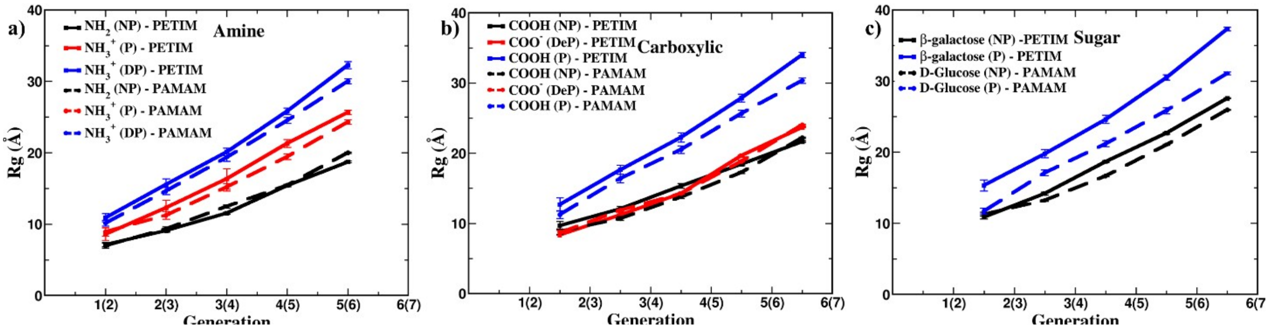

**Figure S11.** The $R_g$ values of O-core PETIM (solid lines) and PAMAM (dashed lines) dendrimers with (a) amine, (b) carboxylic, and (c) sugar (β-galactose/D-glucose) terminations were compared to analyse differences in their sizes. The results indicate that PETIM dendrimers are larger than PAMAM dendrimers with an equivalent number of surface functional groups, with the size difference being most pronounced for sugar terminated dendrimers.

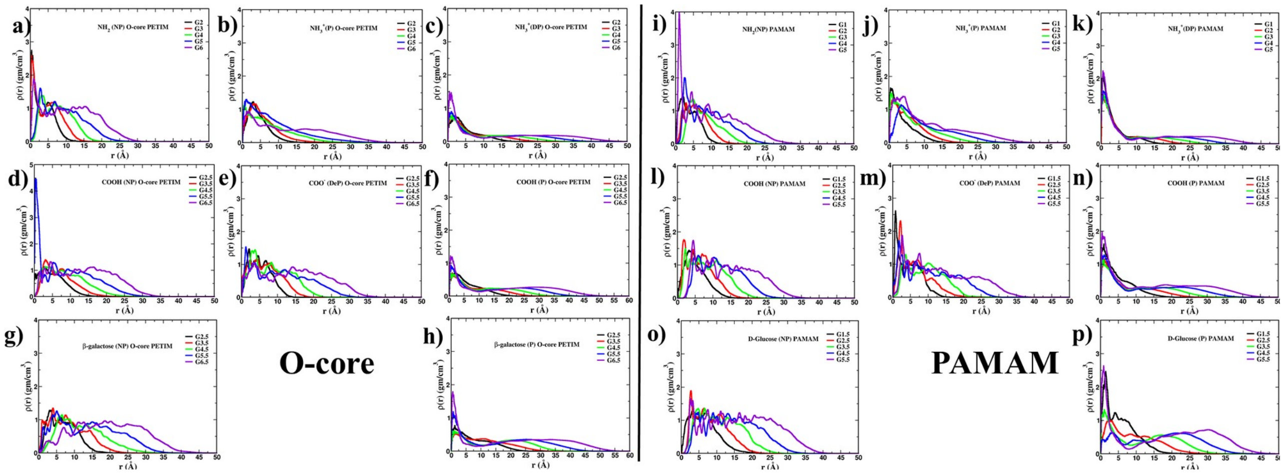

**Figure S12.** The density profiles of O-core PETIM and PAMAM dendrimers with (a-c & i-k) amine, (c-d & l-n) carboxylic, and (g, h & o, p) sugar (β-galactose/D-glucose) terminations were compared to examine differences in their atomic arrangements. The analysis reveals that PETIM dendrimers exhibit broader high-density regions arising from branch back folding than equivalently sized PAMAM dendrimers.

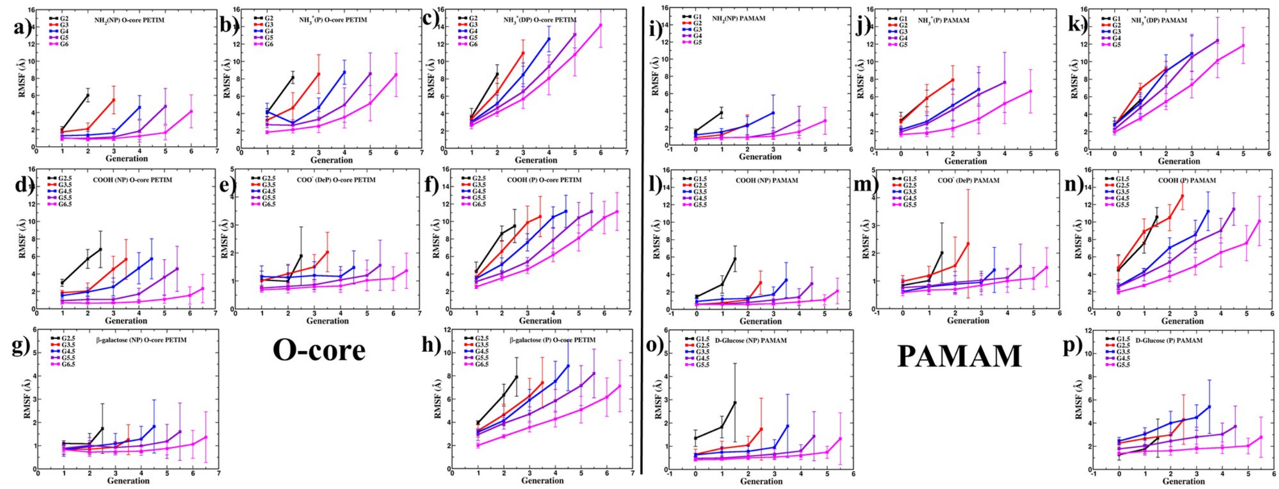

**Figure S13.** The RMSF profiles of O-core PETIM and PAMAM dendrimers with (a, b & f, g) amine, (c, d & h, i) carboxylic, and (e & j) sugar (β-galactose/D-glucose) terminations were compared to assess differences in their flexibilities. The analysis reveals that PETIM dendrimers exhibit greater conformational flexibility than PAMAM dendrimers with the same surface functionalization.

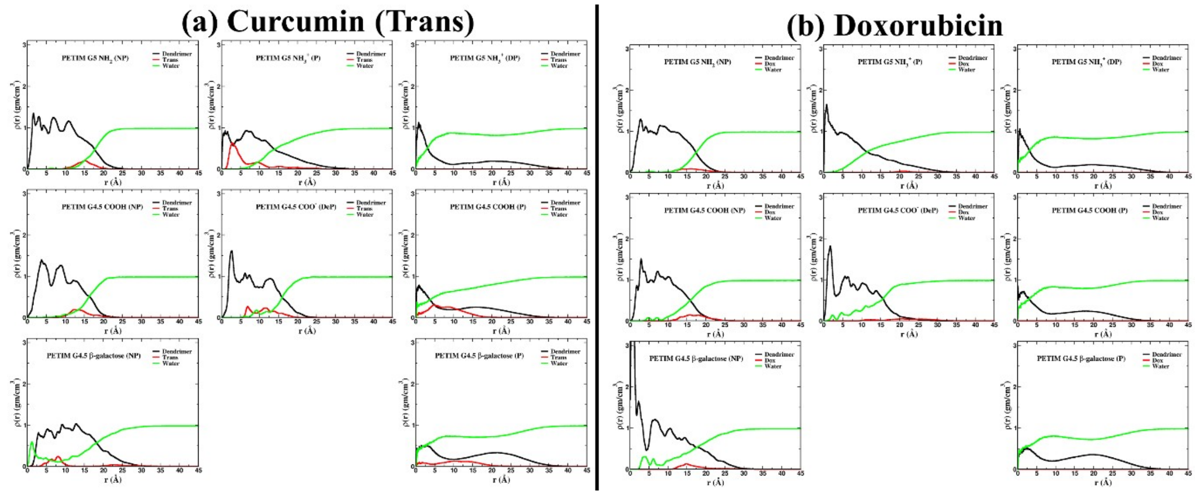

**Figure S14:** Radial density profiles as a function of distance from the centre of mass were computed for O-core PETIM dendrimers with different terminal functional groups in the presence of curcumin (a) and doxorubicin (b). The density profiles reveal that curcumin preferentially binds within the dendrimer interior, indicating stronger encapsulation than doxorubicin.

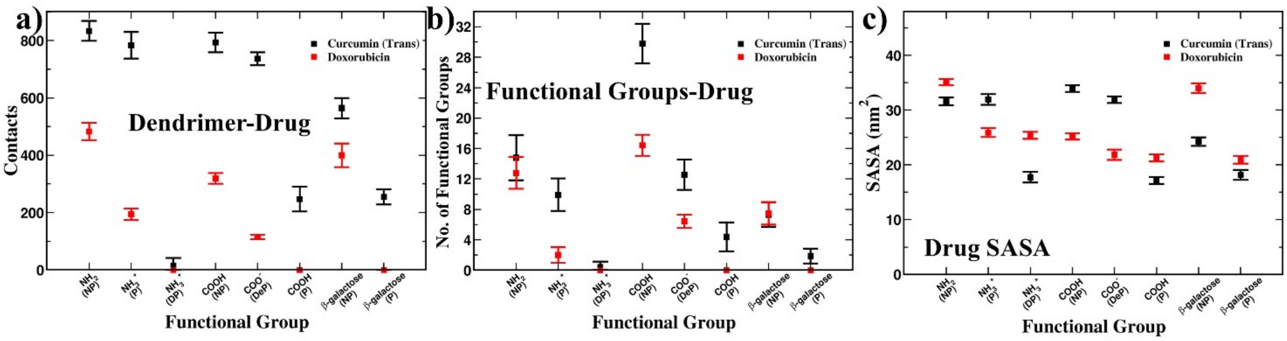

**Figure S15:** Interaction analysis between dendrimers and drug molecules over the simulation trajectory: (a) number of contacts between the dendrimers and drug molecules, (b) number of dendrimer functional groups in close proximity to the drug molecules, and (c) solvent-accessible surface area (SASA) of the drug molecules in various dendrimer-drug complexes. Curcumin maintains a larger number of contacts with the dendrimers than doxorubicin. However, unbound drug clusters (curcumin/doxorubicin) exhibit lower SASA values than when the drugs are bound to dendrimers.

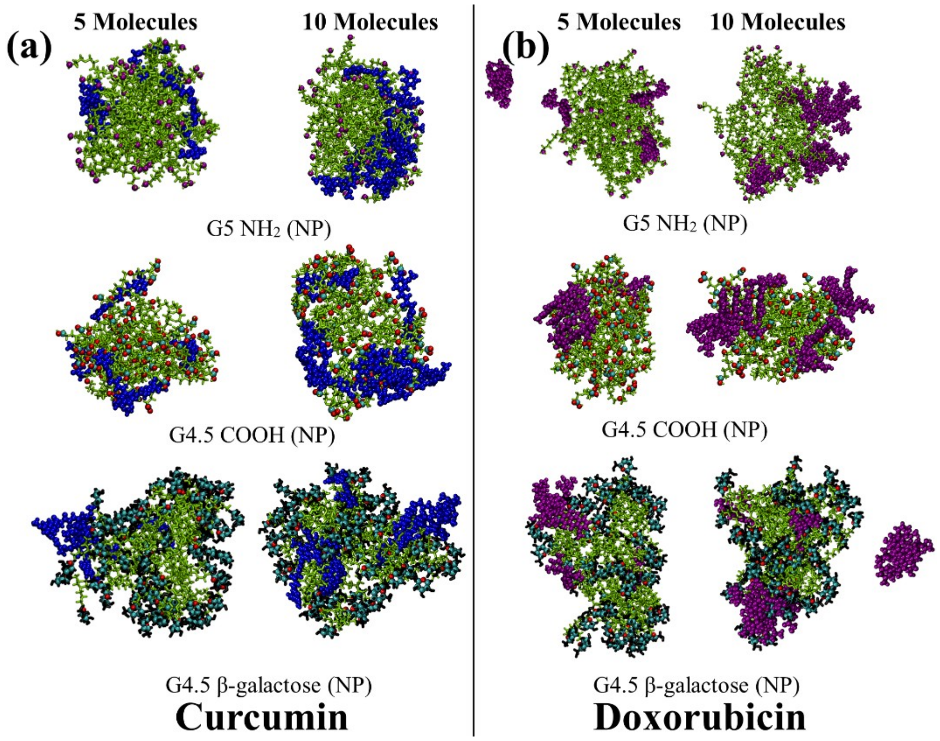

**Figure S16:** Comparison of the final MD snapshots of (a) dendrimer-curcumin and (b) dendrimer-doxorubicin complexes after 200 ns to examine the effect of drug concentration on dendrimer-drug binding. Larger drug clusters are observed in the 1:10 systems compared with the corresponding 1:5 systems, indicating enhanced drug aggregation at higher drug concentrations.

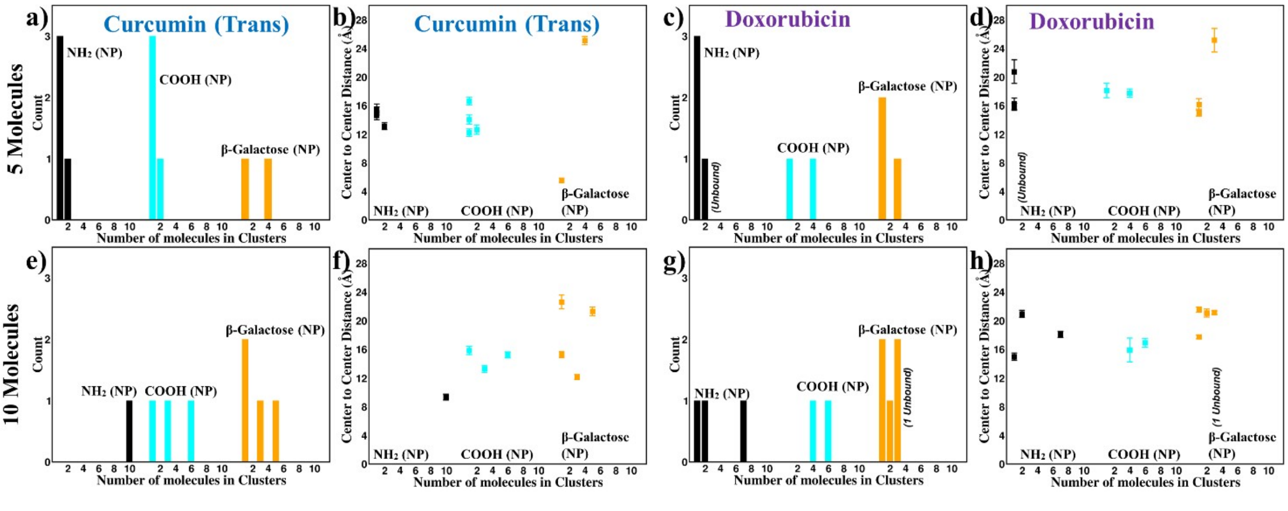

**Figure S17:** The number of clusters containing different numbers of drug molecules was calculated for the (a, c) 1:5 and (e, g) 1:10 dendrimer-drug systems, along with the corresponding center to center distances between clusters, to characterise their spatial organization within the dendrimer. Comparative analysis reveals enhanced clustering at the higher drug loading (1:10), suggesting that increasing the number of drug molecules promotes drug-drug interactions, which become increasingly important in determining their spatial organization within the dendrimer.

| PAMAM | | | | | |
|---|---|---|---|---|---|
| | **Dendrimer_atoms** | **Water_atoms** | **$Na^+$ ion** | **$Cl^-$ ion** | **Total** |
| G1 | | | | | |
| Amine (NP) | 228 | 11637 | 11 | 11 | 11887 |
| Amine (P) | 236 | 11799 | 11 | 19 | 12065 |
| Amine (DP) | 242 | 10881 | 10 | 24 | 11157 |
| $COO^-$ (DeP) | 356 | 16689 | 32 | 16 | 17093 |
| COOH (NP) | 372 | 16701 | 16 | 16 | 17105 |
| COOH (P) | 386 | 8037 | 8 | 22 | 8453 |
| Glucose (NP) | 724 | 26496 | 25 | 25 | 27270 |
| Glucose (P) | 738 | 14916 | 15 | 29 | 15698 |
| G2 | | | | | |
| Amine (NP) | 516 | 23385 | 22 | 22 | 23945 |
| Amine (P) | 532 | 22350 | 21 | 37 | 22940 |
| Amine (DP) | 546 | 25200 | 24 | 54 | 25824 |
| $COO^-$ (DeP) | 772 | 31560 | 62 | 30 | 32424 |
| COOH (NP) | 804 | 29928 | 28 | 28 | 30788 |
| COOH (P) | 834 | 24147 | 23 | 53 | 25057 |
| Glucose (NP) | 1508 | 50304 | 48 | 48 | 51908 |
| Glucose (P) | 1538 | 34806 | 34 | 64 | 36442 |
| G3 | | | | | |
| Amine (NP) | 1092 | 44979 | 42 | 42 | 46155 |
| Amine (P) | 1124 | 39495 | 40 | 72 | 40731 |
| Amine (DP) | 1154 | 38709 | 37 | 99 | 39999 |
| $COO^-$ (DeP) | 1604 | 56586 | 118 | 54 | 58362 |
| COOH (NP) | 1668 | 55230 | 53 | 53 | 57004 |
| COOH (P) | 1730 | 29622 | 29 | 91 | 31472 |
| Glucose (NP) | 3076 | 76017 | 74 | 74 | 79241 |
| Glucose (P) | 3138 | 50682 | 50 | 112 | 53982 |
| G4 | | | | | |
| Amine (NP) | 2244 | 70128 | 67 | 67 | 72506 |
| Amine (P) | 2308 | 66810 | 64 | 128 | 69310 |
| Amine (DP) | 2370 | 67623 | 65 | 191 | 70249 |
| $COO^-$ (DeP) | 3268 | 85659 | 211 | 83 | 89221 |
| COOH (NP) | 3396 | 80115 | 78 | 78 | 83667 |
| COOH (P) | 3522 | 49158 | 49 | 175 | 52904 |
| Glucose (NP) | 6212 | 117129 | 114 | 114 | 123569 |
| Glucose (P) | 6338 | 75990 | 78 | 204 | 82610 |
| G5 | | | | | |
| Amine (NP) | 4548 | 106227 | 102 | 102 | 110979 |
| Amine (P) | 4676 | 101187 | 98 | 226 | 106187 |
| Amine (DP) | 4802 | 103398 | 101 | 355 | 108656 |
| $COO^-$ (DeP) | 6596 | 123471 | 379 | 123 | 130569 |
| COOH (NP) | 6852 | 123309 | 121 | 121 | 130403 |
| COOH (P) | 7106 | 72171 | 73 | 327 | 79677 |
| Glucose (NP) | 12484 | 159213 | 158 | 158 | 172013 |
| Glucose (P) | 12738 | 100764 | 103 | 357 | 113962 |

**Table 1:** Atoms info for PAMAM dendrimers

| PETIM (O-core) |
| --- |

| | **Dend_atom** | **Water_atoms** | **Na** | **Cl** | **Total** |
|---|---|---|---|---|---|
| G2 | | | | | |
| Amine (NP) | 277 | 22644 | 21 | 21 | 22963 |
| Amine (P) | 285 | 22614 | 21 | 29 | 22949 |
| Amine (DP) | 291 | 24927 | 24 | 38 | 25280 |
| $COO^-$ (DeP) | 405 | 30759 | 45 | 29 | 31238 |
| COOH (NP) | 421 | 25260 | 24 | 24 | 25729 |
| COOH (P) | 435 | 32721 | 31 | 45 | 33232 |
| β-galactose (NP) | 773 | 45903 | 43 | 43 | 46762 |
| β-galactose (P) | 787 | 49335 | 46 | 60 | 50228 |
| G3 | | | | | |
| Amine (NP) | 613 | 42060 | 41 | 41 | 42755 |
| Amine (P) | 629 | 43065 | 41 | 57 | 43792 |
| Amine (DP) | 643 | 45414 | 42 | 72 | 46171 |
| $COO^-$ (DeP) | 869 | 56163 | 85 | 53 | 57170 |
| COOH (NP) | 901 | 49752 | 47 | 47 | 50747 |
| COOH (P) | 931 | 53802 | 51 | 81 | 54865 |
| β-galactose (NP) | 1605 | 77625 | 74 | 74 | 79378 |
| β-galactose (P) | 1635 | 77418 | 73 | 103 | 79229 |
| G4 | | | | | |
| Amine (NP) | 1285 | 74139 | 71 | 71 | 75566 |
| Amine (P) | 1317 | 73890 | 71 | 103 | 75381 |
| Amine (DP) | 1347 | 67974 | 64 | 126 | 69511 |
| $COO^-$ (DeP) | 1797 | 92667 | 151 | 87 | 94702 |
| COOH (NP) | 1861 | 77436 | 77 | 77 | 79451 |
| COOH (P) | 1923 | 85686 | 81 | 143 | 87833 |
| β-galactose (NP) | 3269 | 121815 | 116 | 116 | 125316 |
| β-galactose (P) | 3331 | 117969 | 112 | 174 | 121586 |
| G5 | | | | | |
| Amine (NP) | 2629 | 117009 | 110 | 110 | 119858 |
| Amine (P) | 2693 | 115086 | 110 | 174 | 118063 |
| Amine (DP) | 2755 | 122268 | 116 | 242 | 125381 |
| $COO^-$ (DeP) | 3653 | 143133 | 263 | 135 | 147184 |
| COOH (NP) | 3781 | 127101 | 122 | 122 | 131126 |
| COOH (P) | 3907 | 132303 | 126 | 252 | 136588 |
| β-galactose (NP) | 6597 | 179481 | 172 | 172 | 186422 |
| β-galactose (P) | 6723 | 187245 | 180 | 306 | 194454 |
| G6 | | | | | |
| Amine (NP) | 5317 | 171318 | 164 | 164 | 176963 |
| Amine (P) | 5445 | 165582 | 159 | 287 | 171473 |
| Amine (DP) | 5571 | 195057 | 187 | 441 | 201256 |
| $COO^-$ (DeP) | 7365 | 200259 | 450 | 194 | 208268 |
| COOH (NP) | 7621 | 185370 | 179 | 179 | 193349 |
| COOH (P) | 7875 | 208509 | 206 | 460 | 217050 |
| β-galactose (NP) | 13253 | 245748 | 240 | 240 | 259481 |
| β-galactose (P) | 13507 | 262539 | 255 | 509 | 276810 |

**Table 2:** Atoms info for PETIM (O-core) dendrimers

| PETIM (N-core) | | | | | |
|---|---|---|---|---|---|
| | **Dend_atom** | **Water_atoms** | **Na** | **Cl** | **Total** |
| G2 | | | | | |
| Amine (NP) | 445 | 31098 | 29 | 29 | 31601 |
| Amine (P) | 457 | 31110 | 30 | 42 | 31639 |
| Amine (DP) | 467 | 30249 | 28 | 50 | 30794 |
| $COO^-$ (DeP) | 637 | 39996 | 64 | 40 | 40737 |
| COOH (NP) | 661 | 37380 | 36 | 36 | 38113 |
| COOH (P) | 683 | 37449 | 36 | 58 | 38226 |
| β-galactose (NP) | 1189 | 55005 | 52 | 52 | 56298 |
| β-galactose (P) | 1211 | 52986 | 50 | 72 | 54319 |
| G3 | | | | | |
| Amine (NP) | 949 | 49773 | 47 | 47 | 50816 |
| Amine (P) | 973 | 48621 | 46 | 70 | 49710 |
| Amine (DP) | 995 | 49626 | 47 | 93 | 50761 |
| $COO^-$ (DeP) | 1333 | 61854 | 107 | 59 | 63353 |
| COOH (NP) | 1381 | 55488 | 53 | 53 | 56975 |
| COOH (P) | 1427 | 64221 | 61 | 107 | 65816 |
| β-galactose (NP) | 2437 | 86109 | 82 | 82 | 88710 |
| β-galactose (P) | 2483 | 84675 | 80 | 126 | 87364 |
| G4 | | | | | |
| Amine (NP) | 1957 | 87762 | 83 | 83 | 89885 |
| Amine (P) | 2005 | 88467 | 84 | 132 | 90688 |
| Amine (DP) | 2051 | 92484 | 87 | 181 | 94803 |
| $COO^-$ (DeP) | 2725 | 111654 | 202 | 106 | 114687 |
| COOH (NP) | 2821 | 102462 | 98 | 98 | 105479 |
| COOH (P) | 2915 | 107034 | 101 | 195 | 110245 |
| β-galactose (NP) | 4933 | 131736 | 128 | 128 | 136925 |
| β-galactose (P) | 5027 | 130074 | 125 | 219 | 135445 |
| G5 | | | | | |
| Amine (NP) | 3973 | 125010 | 120 | 120 | 129223 |
| Amine (P) | 4069 | 127659 | 123 | 219 | 132070 |
| Amine (DP) | 4163 | 131955 | 128 | 318 | 136564 |
| $COO^-$ (DeP) | 5509 | 152850 | 339 | 147 | 158845 |
| COOH (NP) | 5701 | 139101 | 134 | 134 | 145070 |
| COOH (P) | 5891 | 158235 | 152 | 342 | 164620 |
| β-galactose (NP) | 9925 | 189552 | 185 | 185 | 199847 |
| β-galactose (P) | 10115 | 189813 | 186 | 376 | 200490 |
| G6 | | | | | |
| Amine (NP) | 4069 | 127659 | 123 | 219 | 132070 |
| Amine (P) | 8197 | 190098 | 184 | 376 | 198855 |
| Amine (DP) | 8387 | 195918 | 192 | 574 | 205071 |
| $COO^-$ (DeP) | 11077 | 224061 | 603 | 219 | 235960 |
| COOH (NP) | 11461 | 205821 | 207 | 207 | 217696 |
| COOH (P) | 11843 | 219672 | 213 | 595 | 232323 |
| β-galactose (NP) | 19909 | 261033 | 259 | 259 | 281460 |
| β-galactose (P) | 20291 | 264507 | 265 | 647 | 285710 |

**Table 3:** Atoms info for PETIM (N-core) dendrimers

| PAMAM EDA-core ‘aaa’ (NP) | | | PAMAM EDA-core ‘aaa’ (P) | | |
|---|---|---|---|---|---|
| **Atom** | **GAFF atom type** | **Charge** | **Atom** | **GAFF atom type** | **Charge** |
| C1 | c3 | 0.0289 | N1 | n4 | -0.0182 |
| H1 | h1 | 0.0846 | C1 | c3 | 0.0124 |
| H2 | h1 | 0.0846 | C2 | c3 | 0.0124 |
| C2 | c3 | 0.0289 | N2 | n4 | -0.0182 |
| H3 | h1 | 0.0846 | C3 | c3 | -0.0143 |
| H4 | h1 | 0.0846 | C4 | c3 | -0.0143 |
| N1 | n3 | -0.6577 | C5 | c3 | -0.0143 |
| N2 | n3 | -0.6577 | C6 | c3 | -0.0143 |
| C3 | c3 | 0.3694 | C7 | c3 | -0.2676 |
| H5 | h1 | 0.0065 | C8 | c | 0.7134 |
| H6 | h1 | 0.0065 | C9 | c3 | -0.2676 |
| C4 | c3 | 0.3694 | C10 | c | 0.7134 |
| H7 | h1 | 0.0065 | C11 | c3 | -0.2676 |
| H8 | h1 | 0.0065 | C12 | c | 0.7134 |
| C5 | c3 | 0.3694 | C13 | c3 | -0.2676 |
| H9 | h1 | 0.0065 | C14 | c | 0.7134 |
| H10 | h1 | 0.0065 | O1 | o | -0.5553 |
| C6 | c3 | 0.3694 | O2 | o | -0.5553 |
| H11 | h1 | 0.0065 | O3 | o | -0.5553 |
| H12 | h1 | 0.0065 | O4 | o | -0.5553 |
| C7 | c3 | -0.5435 | H1 | hx | 0.0828 |
| H13 | hc | 0.1589 | H2 | hx | 0.0828 |
| H14 | hc | 0.1589 | H3 | hx | 0.0828 |
| C8 | c3 | -0.5435 | H4 | hx | 0.0828 |
| H15 | hc | 0.1589 | H5 | hx | 0.1047 |
| H16 | hc | 0.1589 | H6 | hx | 0.1047 |
| C9 | c3 | -0.5435 | H7 | hx | 0.1047 |
| H17 | hc | 0.1589 | H8 | hx | 0.1047 |
| H18 | hc | 0.1589 | H9 | hx | 0.1047 |
| C10 | c3 | -0.5435 | H10 | hx | 0.1047 |
| H19 | hc | 0.1589 | H11 | hx | 0.1047 |
| H20 | hc | 0.1589 | H12 | hx | 0.1047 |
| C11 | c | 0.6780 | H13 | hc | 0.1183 |
| C12 | c | 0.6780 | H14 | hc | 0.1183 |
| C13 | c | 0.6780 | H15 | hc | 0.1183 |
| C14 | c | 0.6780 | H16 | hc | 0.1183 |
| O1 | o | -0.6049 | H17 | hc | 0.1183 |
| O2 | o | -0.6049 | H18 | hc | 0.1183 |
| O3 | o | -0.6049 | H19 | hc | 0.1183 |
| O4 | o | -0.6049 | H20 | hc | 0.1183 |
| | | | H37 | hn | 0.1960 |
| | | | H38 | hn | 0.1960 |

**Table 4:** Charge information of PAMAM (EDA-core)

| **PAMAM Repeating Residue ‘bbb’ (NP)** | | | **PAMAM Repeating Residue ‘bbb’ (P)** | | |
|---|---|---|---|---|---|
| **Atom** | **GAFF atom type** | **Charge** | **Atom** | **GAFF atom type** | **Charge** |
| C1 | c3 | 0.4418 | N1 | n4 | -0.0275 |
| H1 | h1 | -0.0353 | C1 | c3 | -0.0510 |
| H2 | h1 | -0.0353 | C2 | c3 | -0.0510 |
| C2 | c3 | -0.2825 | C3 | c3 | -0.1846 |
| H3 | h1 | 0.1571 | C4 | c | 0.6725 |
| H4 | h1 | 0.1571 | C5 | c3 | -0.1846 |
| N1 | n3 | -0.7409 | C6 | c | 0.6725 |
| C3 | c3 | 0.5168 | C7 | c3 | -0.0208 |
| H5 | h1 | -0.0523 | C8 | c3 | -0.0333 |
| H6 | h1 | -0.0523 | N2 | n | -0.3330 |
| C4 | c3 | 0.5168 | O1 | o | -0.5292 |
| H7 | h1 | -0.0523 | O2 | o | -0.5292 |
| H8 | h1 | -0.0523 | H1 | hx | 0.1156 |
| C5 | c3 | -0.6039 | H2 | hx | 0.1156 |
| H9 | hc | 0.1673 | H3 | hx | 0.1156 |
| H10 | hc | 0.1673 | H4 | hx | 0.1156 |
| C6 | c3 | -0.6039 | H5 | hc | 0.0835 |
| H11 | hc | 0.1673 | H6 | hc | 0.0835 |
| H12 | hc | 0.1673 | H7 | hc | 0.0835 |
| C7 | c | 0.7211 | H8 | hc | 0.0835 |
| C8 | c | 0.7211 | H9 | hx | 0.0799 |
| O1 | o | -0.6195 | H10 | hx | 0.0799 |
| O2 | o | -0.6195 | H11 | h1 | 0.1062 |
| N4 | n | -0.4658 | H12 | h1 | 0.1062 |
| H21 | hn | 0.3148 | H13 | hn | 0.2592 |
| | | | H25 | hn | 0.1713 |

**Table 5:** Charge information of PAMAM (repeating unit)

| **PAMAM Terminal Residue ‘ccc’ Amine (NP)** | | | **PAMAM Terminal Residue ‘ccc’ Amine (P)** | | |
|---|---|---|---|---|---|
| Atom | GAFF atom type | Charge | Atom | GAFF atom type | Charge |
| C1 | c3 | -0.0240 | N1 | n | -0.2641 |
| H1 | h1 | 0.1017 | H4 | hn | 0.2763 |
| H2 | h1 | 0.1017 | C3 | c3 | -0.1790 |
| C2 | c3 | 0.5257 | C4 | c3 | 0.0800 |
| H3 | h1 | -0.0249 | H5 | h1 | 0.1387 |
| H4 | h1 | -0.0249 | H6 | h1 | 0.1387 |
| N1 | n3 | -1.3377 | H7 | hx | 0.0779 |
| H5 | hn | 0.4721 | H8 | hx | 0.0779 |
| H6 | hn | 0.4721 | N2 | n4 | -0.1678 |
| N2 | n | -0.5918 | H9 | hn | 0.2738 |
| H7 | hn | 0.3300 | H10 | hn | 0.2738 |
| | | | H11 | hn | 0.2738 |

**Table 6:** Charge information of PAMAM (amine functionalised group)

| PAMAM Terminal Residue 'ccc' Carboxylic (DeP) | | | PAMAM Terminal Residue 'ccc' Carboxylic (NP) | | |
|---|---|---|---|---|---|
| **Atom** | **GAFF atom type** | **Charge** | **Atom** | **GAFF atom type** | **Charge** |
| N1 | n | -0.6370 | N1 | n | -0.8725 |
| H4 | hn | 0.3423 | H1 | hn | 0.4604 |
| C3 | c3 | 0.0109 | C1 | c3 | 0.6548 |
| H5 | h1 | 0.1119 | H2 | h1 | -0.0896 |
| C4 | c3 | -0.0616 | C2 | c3 | 0.0362 |
| H6 | h1 | 0.1119 | H3 | h1 | -0.0896 |
| H7 | h1 | 0.1125 | H4 | h1 | 0.0693 |
| H8 | h1 | 0.1125 | H5 | h1 | 0.0693 |
| N2 | n3 | -0.4929 | N2 | n3 | -0.6136 |
| C5 | c3 | 0.0012 | C3 | c3 | 0.1794 |
| C6 | c3 | 0.0012 | C4 | c3 | 0.1794 |
| H9 | h1 | 0.0535 | H6 | h1 | 0.0734 |
| C7 | c3 | 0.0310 | C5 | c3 | -0.4908 |
| H10 | h1 | 0.0535 | H7 | h1 | 0.0734 |
| H11 | hc | -0.0338 | H8 | hc | 0.1510 |
| H12 | hc | -0.0338 | H9 | hc | 0.1510 |
| C8 | c | 0.9307 | C6 | c | 1.0022 |
| O2 | o | -0.9038 | O1 | oh | -0.7731 |
| O3 | o | -0.9038 | O2 | o | -0.6643 |
| H13 | h1 | 0.0535 | H10 | h1 | 0.0734 |
| H14 | h1 | 0.0535 | H11 | h1 | 0.0734 |
| C9 | c3 | 0.0310 | C7 | c3 | -0.4908 |
| H15 | hc | -0.0338 | H12 | hc | 0.1510 |
| H16 | hc | -0.0338 | H13 | hc | 0.1510 |
| C10 | c | 0.9307 | C8 | c | 1.0022 |
| O4 | o | -0.9038 | O3 | o | -0.6643 |
| O5 | o | -0.9038 | O4 | oh | -0.7731 |
| | | | H14 | ho | 0.4855 |
| | | | H15 | ho | 0.4855 |

**Table 7:** Charge information of PAMAM (carboxylic functionalised group)

| PAMAM Terminal Residue 'ccc' D-glucose (NP) | | | PAMAM Terminal Residue 'ccc' D-glucose (P) | | |
|---|---|---|---|---|---|
| **Atom** | **GAFF atom type** | **Charge** | **Atom** | **GAFF atom type** | **Charge** |
| N1 | n | -0.4658 | N1 | n | -0.33303 |
| H4 | hn | 0.3148 | H4 | hn | 0.259161 |
| C3 | c3 | -0.2825 | C3 | c3 | -0.033281 |
| H5 | h1 | 0.1571 | H5 | h1 | 0.106181 |
| C4 | c3 | 0.4418 | C4 | c3 | -0.02083 |
| H6 | h1 | 0.1571 | H6 | h1 | 0.106181 |
| H7 | h1 | -0.0353 | H7 | h1 | 0.07988 |
| H8 | h1 | -0.0353 | H8 | h1 | 0.07988 |
| N2 | n3 | -0.7409 | N2 | n4 | -0.027479 |
| C5 | c3 | 0.5168 | C5 | c3 | -0.050976 |
| C6 | c3 | 0.5168 | C6 | c3 | -0.050976 |
| H9 | h1 | -0.0523 | H9 | h1 | 0.115633 |
| C7 | c3 | -0.6039 | C7 | c3 | -0.184579 |
| H10 | h1 | -0.0523 | H10 | h1 | 0.115633 |
| H11 | hc | 0.1673 | H11 | hc | 0.083479 |
| H12 | hc | 0.1673 | H12 | hc | 0.083479 |
| C8 | c | 0.7211 | C8 | c | 0.672544 |
| O2 | o | -0.6195 | O2 | o | -0.529172 |
| H13 | h1 | -0.0523 | H13 | h1 | 0.115633 |
| H14 | h1 | -0.0523 | H14 | h1 | 0.115633 |
| C9 | c3 | -0.6039 | C9 | c3 | -0.184579 |
| H15 | hc | 0.1673 | H15 | hc | 0.083479 |

| | | | | | |
|---|---|---|---|---|---|
| H16 | hc | 0.1673 | H16 | hc | 0.083479 |
| C10 | c | 0.7211 | C10 | c | 0.672544 |
| O4 | o | -0.6195 | O4 | o | -0.529172 |
| C11 | c3 | 0.0852 | C11 | c3 | 0.08518 |
| C12 | c3 | 0.0792 | C12 | c3 | 0.07918 |
| H17 | h1 | 0.0698 | H17 | h1 | 0.06978 |
| C13 | c3 | 0.0792 | C13 | c3 | 0.07918 |
| C14 | c3 | 0.0358 | C14 | c3 | 0.03578 |
| C15 | c3 | 0.3350 | C15 | c3 | 0.33498 |
| H18 | h1 | 0.0618 | H18 | h1 | 0.06178 |
| H19 | h1 | 0.1128 | H19 | h1 | 0.11278 |
| H20 | h2 | 0.1218 | H20 | h2 | 0.12178 |
| O1 | oh | -0.5697 | O1 | oh | -0.56972 |
| O6 | oh | -0.5777 | O6 | oh | -0.57772 |
| C16 | c3 | 0.1455 | C16 | c3 | 0.14548 |
| H21 | ho | 0.4051 | H21 | ho | 0.40508 |
| H22 | hn | 0.3176 | H22 | hn | 0.31758 |
| H23 | h1 | 0.0628 | H23 | h1 | 0.06278 |
| H24 | h1 | 0.0628 | H24 | h1 | 0.06278 |
| O7 | oh | -0.5947 | O7 | oh | -0.59472 |
| H25 | ho | 0.4191 | H25 | ho | 0.41908 |
| O8 | os | -0.4455 | O8 | os | -0.44552 |
| H26 | ho | 0.4191 | H26 | ho | 0.41908 |
| H27 | h1 | 0.0858 | H27 | h1 | 0.08578 |
| O9 | oh | -0.5867 | O9 | oh | -0.58672 |
| H28 | ho | 0.4191 | H28 | ho | 0.41908 |
| N3 | n3 | -0.5428 | N3 | n3 | -0.54282 |
| C17 | c3 | 0.0852 | C17 | c3 | 0.08518 |
| C18 | c3 | 0.0792 | C18 | c3 | 0.07918 |
| H29 | h1 | 0.0698 | H29 | h1 | 0.06978 |
| C19 | c3 | 0.0792 | C19 | c3 | 0.07918 |
| C20 | c3 | 0.0358 | C20 | c3 | 0.03578 |
| C21 | c3 | 0.3350 | C21 | c3 | 0.33498 |
| H30 | h1 | 0.0618 | H30 | h1 | 0.06178 |
| H31 | h1 | 0.1128 | H31 | h1 | 0.11278 |
| H32 | h2 | 0.1218 | H32 | h2 | 0.12178 |
| O10 | oh | -0.5697 | O10 | oh | -0.56972 |
| O11 | oh | -0.5777 | O11 | oh | -0.57772 |
| C22 | c3 | 0.1455 | C22 | c3 | 0.14548 |
| H33 | ho | 0.4051 | H33 | ho | 0.40508 |
| H34 | hn | 0.3176 | H34 | hn | 0.31758 |
| H35 | h1 | 0.0628 | H35 | h1 | 0.06278 |
| H36 | h1 | 0.0628 | H36 | h1 | 0.06278 |
| O12 | oh | -0.5947 | O12 | oh | -0.59472 |
| H37 | ho | 0.4191 | H37 | ho | 0.41908 |
| O13 | os | -0.4455 | O13 | os | -0.44552 |
| H38 | ho | 0.4191 | H38 | ho | 0.41908 |
| H39 | h1 | 0.0858 | H39 | h1 | 0.08578 |
| O5 | oh | -0.5867 | O5 | oh | -0.58672 |
| H40 | ho | 0.4191 | H40 | ho | 0.41908 |
| N4 | n3 | -0.5428 | N4 | n3 | -0.54282 |
| | | | H41 | hn | 0.171255 |

**Table 8:** Charge information of PAMAM (D-Glucose functionalised group)

| O-core 'aaa' (NP) | | | O-core 'aaa' (P) | | |
|---|---|---|---|---|---|
| **Atom** | **GAFF atom type** | **Charge** | **Atom** | **GAFF atom type** | **Charge** |
| O1 | os | -0.4381 | O1 | os | -0.5811 |
| C1 | c3 | 0.0451 | C1 | c3 | 0.1771 |
| H1 | h1 | 0.0477 | H1 | h1 | 0.0457 |
| H2 | h1 | 0.0477 | H2 | h1 | 0.0457 |
| C2 | c3 | 0.0451 | C2 | c3 | 0.1771 |
| H3 | h1 | 0.0477 | H3 | h1 | 0.0457 |
| H4 | h1 | 0.0477 | H4 | h1 | 0.0457 |
| C3 | c3 | 0.0566 | C3 | c3 | 0.1510 |
| H5 | hc | 0.0493 | H5 | hc | 0.0254 |
| H6 | hc | 0.0493 | H6 | hc | 0.0254 |
| C4 | c3 | 0.0566 | C4 | c3 | 0.1510 |
| H7 | hc | 0.0493 | H7 | hc | 0.0254 |
| H8 | hc | 0.0493 | H8 | hc | 0.0254 |
| C5 | c3 | 0.0099 | C5 | c3 | -0.2947 |
| H9 | h1 | 0.0485 | H9 | hx | 0.1253 |
| H10 | h1 | 0.0485 | H10 | hx | 0.1253 |
| C6 | c3 | 0.0099 | C6 | c3 | -0.2947 |
| H11 | h1 | 0.0485 | H11 | hx | 0.1253 |
| H12 | h1 | 0.0485 | H12 | hx | 0.1253 |
| N1 | n3 | -0.5994 | N1 | n4 | -0.1145 |
| N2 | n3 | -0.5994 | N2 | n4 | -0.1145 |
| C7 | c3 | 0.0803 | C7 | c3 | -0.0343 |
| H13 | h1 | 0.0507 | H13 | hx | 0.1052 |
| H14 | h1 | 0.0507 | H14 | hx | 0.1052 |
| C8 | c3 | 0.0803 | C8 | c3 | -0.0343 |
| H15 | h1 | 0.0507 | H15 | hx | 0.1052 |
| H16 | h1 | 0.0507 | H16 | hx | 0.1052 |
| C9 | c3 | 0.0803 | C9 | c3 | -0.0343 |
| H17 | h1 | 0.0507 | H17 | hx | 0.1052 |
| H18 | h1 | 0.0507 | H18 | hx | 0.1052 |
| C10 | c3 | 0.0803 | C10 | c3 | -0.0343 |
| H19 | h1 | 0.0507 | H19 | hx | 0.1052 |
| H20 | h1 | 0.0507 | H20 | hx | 0.1052 |
| C11 | c3 | 0.1029 | C11 | c3 | -0.0618 |
| H21 | hc | 0.0679 | H21 | hc | 0.0887 |
| H22 | hc | 0.0679 | H22 | hc | 0.0887 |
| C12 | c3 | 0.1029 | C12 | c3 | -0.0618 |
| H23 | hc | 0.0679 | H23 | hc | 0.0887 |
| H24 | hc | 0.0679 | H24 | hc | 0.0887 |
| C13 | c3 | 0.1029 | C13 | c3 | -0.0618 |
| H25 | hc | 0.0679 | H25 | hc | 0.0887 |
| H26 | hc | 0.0679 | H26 | hc | 0.0887 |
| C14 | c3 | 0.1029 | C14 | c3 | -0.0618 |
| H27 | hc | 0.0679 | H27 | hc | 0.0887 |
| H28 | hc | 0.0679 | H28 | hc | 0.0887 |
| C15 | c3 | -0.7259 | C15 | c3 | -0.2222 |
| H29 | h1 | 0.2567 | H29 | hc | 0.1644 |
| H30 | h1 | 0.2567 | H30 | hc | 0.1644 |
| C16 | c3 | -0.7259 | C16 | c3 | -0.2222 |
| H31 | h1 | 0.2567 | H31 | hc | 0.1644 |
| H32 | h1 | 0.2567 | H32 | hc | 0.1644 |
| C17 | c3 | -0.7259 | C17 | c3 | -0.2222 |
| H33 | h1 | 0.2567 | H33 | hc | 0.1644 |
| H34 | h1 | 0.2567 | H34 | hc | 0.1644 |
| C18 | c3 | -0.7259 | C18 | c3 | -0.2222 |
| H35 | h1 | 0.2567 | H35 | hc | 0.1644 |
| H36 | h1 | 0.2567 | H36 | hc | 0.1644 |
| | | | H38 | hn | 0.1827 |
| | | | H37 | hn | 0.1827 |

**Table 9:** Charge information of PETIM O-core

| N-core 'aaa' (NP) | | | N-core 'aaa' (P) | | |
|---|---|---|---|---|---|
| **Atom** | **GAFF atom type** | **Charge** | **Atom** | **GAFF atom type** | **Charge** |
| N1 | n3 | -0.6542 | N1 | n4 | -0.2480 |
| C1 | c3 | 0.0518 | C1 | c3 | 0.1573 |
| H1 | h1 | 0.0648 | H1 | hx | 0.0507 |
| H2 | h1 | 0.0648 | H2 | hx | 0.0507 |
| C2 | c3 | 0.0518 | C2 | c3 | 0.1573 |
| H3 | h1 | 0.0648 | H3 | hx | 0.0507 |
| H4 | h1 | 0.0648 | H4 | hx | 0.0507 |
| C3 | c3 | 0.0518 | C3 | c3 | 0.1573 |
| H5 | h1 | 0.0648 | H5 | hx | 0.0507 |
| H6 | h1 | 0.0648 | H6 | hx | 0.0507 |
| C4 | c3 | 0.0787 | C4 | c3 | 0.0544 |
| H7 | hc | 0.0868 | H7 | hc | 0.0771 |
| H8 | hc | 0.0868 | H8 | hc | 0.0771 |
| C5 | c3 | 0.0787 | C5 | c3 | 0.0544 |
| H9 | hc | 0.0868 | H9 | hc | 0.0771 |
| H10 | hc | 0.0868 | H10 | hc | 0.0771 |
| C6 | c3 | 0.0787 | C6 | c3 | 0.0544 |
| H11 | hc | 0.0868 | H11 | hc | 0.0771 |
| H12 | hc | 0.0868 | H12 | hc | 0.0771 |
| C7 | c3 | -0.7494 | C7 | c3 | -0.7457 |
| H13 | h1 | 0.2669 | H13 | h1 | 0.3084 |
| H14 | h1 | 0.2669 | H14 | h1 | 0.3084 |
| C8 | c3 | -0.7494 | C8 | c3 | -0.7457 |
| H15 | h1 | 0.2669 | H15 | h1 | 0.3084 |
| H16 | h1 | 0.2669 | H16 | h1 | 0.3084 |
| C9 | c3 | -0.7494 | C9 | c3 | -0.7457 |
| H17 | h1 | 0.2669 | H17 | h1 | 0.3084 |
| H18 | h1 | 0.2669 | H18 | h1 | 0.3084 |
| | | | H19 | hn | 0.2330 |

**Table 10:** Charge information of PETIM N-core

| PETIM Repeating Residue 'bbb' (NP) | | | PETIM Repeating Residue 'bbb' (P) | | |
|---|---|---|---|---|---|
| **Atom** | **GAFF atom type** | **Charge** | **Atom** | **GAFF atom type** | **Charge** |
| N1 | n3 | -0.5969 | N1 | n4 | 0.0373 |
| C1 | c3 | 0.1014 | C1 | c3 | -0.0702 |
| H1 | h1 | 0.0441 | H1 | hx | 0.1222 |
| H2 | h1 | 0.0441 | H2 | hx | 0.1222 |
| C2 | c3 | 0.1014 | C2 | c3 | -0.0702 |
| H3 | h1 | 0.0441 | H3 | hx | 0.1222 |
| H4 | h1 | 0.0441 | H4 | hx | 0.1222 |
| C3 | c3 | -0.0020 | C3 | c3 | -0.3901 |
| H5 | h1 | 0.0506 | H5 | hx | 0.1494 |
| H6 | h1 | 0.0506 | H6 | hx | 0.1494 |
| C4 | c3 | -0.0161 | C4 | c3 | 0.1806 |
| H7 | hc | 0.0494 | H7 | hc | 0.0179 |
| H8 | hc | 0.0494 | H8 | hc | 0.0179 |
| C5 | c3 | 0.1136 | C5 | c3 | -0.1134 |
| H9 | hc | 0.0627 | H9 | hc | 0.0952 |
| H10 | hc | 0.0627 | H10 | hc | 0.0952 |
| C6 | c3 | 0.1136 | C6 | c3 | -0.1134 |
| H11 | hc | 0.0627 | H11 | hc | 0.0952 |
| H12 | hc | 0.0627 | H12 | hc | 0.0952 |
| C7 | c3 | 0.0586 | C7 | c3 | 0.2184 |
| H13 | h1 | 0.0612 | H13 | h1 | 0.0450 |
| H14 | h1 | 0.0612 | H14 | h1 | 0.0450 |
| C8 | c3 | -0.7242 | C8 | c3 | -0.0646 |
| H15 | h1 | 0.2564 | H15 | h1 | 0.1355 |
| H16 | h1 | 0.2564 | H16 | h1 | 0.1355 |
| C9 | c3 | -0.7242 | C9 | c3 | -0.0646 |

| | | | | | |
|---|---|---|---|---|---|
| H17 | h1 | 0.2564 | H17 | h1 | 0.1355 |
| H18 | h1 | 0.2564 | H18 | h1 | 0.1355 |
| O1 | os | -0.2003 | O1 | os | -0.6259 |
| | | | H19 | hn | 0.2398 |

**Table 11:** Charge information of PETIM (repeating unit)

| **PETIM Terminal Residue 'ccc' Amine (NP)** | | | **PETIM Terminal Residue 'ccc' Amine (P)** | | |
|---|---|---|---|---|---|
| **Atom** | **GAFF atom type** | **Charge** | **Atom** | **GAFF atom type** | **Charge** |
| C1 | c3 | 0.6021 | C1 | c3 | 0.1706 |
| H1 | h1 | -0.0925 | H1 | hx | 0.0421 |
| H2 | h1 | -0.0925 | H2 | hx | 0.0421 |
| C2 | c3 | -0.0254 | C2 | c3 | -0.0450 |
| H3 | hc | 0.0085 | H3 | hc | 0.0265 |
| H4 | hc | 0.0085 | H4 | hc | 0.0265 |
| C3 | c3 | -0.0407 | C3 | c3 | 0.0473 |
| H5 | h1 | 0.0461 | H5 | h1 | 0.0703 |
| H6 | h1 | 0.0461 | H6 | h1 | 0.0703 |
| N1 | n3 | -1.1180 | N1 | n4 | -0.2108 |
| H7 | hn | 0.3842 | H7 | hn | 0.2840 |
| H8 | hn | 0.3842 | H8 | hn | 0.2840 |
| O1 | os | -0.1106 | O1 | os | -0.0920 |
| | | | H17 | hn | 0.2840 |

**Table 12:** Charge information of PETIM (amine functionalised group)

| **PETIM Terminal Residue 'ccc' Carboxylic (DeP)** | | | **PETIM Terminal Residue 'ccc' Carboxylic (NP)** | | |
|---|---|---|---|---|---|
| **Atom** | **GAFF atom type** | **Charge** | **Atom** | **GAFF atom type** | **Charge** |
| C3 | c3 | -0.0407 | C1 | c3 | 0.5075 |
| H5 | h1 | 0.0461 | H1 | h1 | -0.0281 |
| H6 | h1 | 0.0461 | H2 | h1 | -0.0281 |
| C2 | c3 | -0.0254 | C2 | c3 | -0.1549 |
| H3 | hc | 0.0085 | H3 | hc | 0.1027 |
| H4 | hc | 0.0085 | H4 | hc | 0.1027 |
| C1 | c3 | 0.6021 | C3 | c3 | -0.1569 |
| H1 | h1 | -0.0925 | H5 | h1 | 0.0850 |
| H2 | h1 | -0.0925 | H6 | h1 | 0.0850 |
| N1 | n3 | -0.5970 | N1 | n3 | -0.2995 |
| C4 | c3 | 0.3280 | C4 | c3 | -0.0132 |
| H7 | h1 | -0.0712 | H7 | h1 | 0.0817 |
| H8 | h1 | -0.0712 | H8 | h1 | 0.0817 |
| C5 | c3 | -0.3933 | C5 | c3 | -0.2218 |
| H9 | hc | 0.0813 | H9 | hc | 0.0967 |
| H10 | hc | 0.0813 | H10 | hc | 0.0967 |
| C6 | c | 1.0897 | C6 | c | 0.9965 |
| O2 | o | -0.9605 | O1 | o | -0.6965 |
| O3 | o | -0.9605 | O2 | oh | -0.8408 |
| C7 | c3 | 0.3280 | H11 | ho | 0.5261 |
| H11 | h1 | -0.0712 | C7 | c3 | -0.0132 |
| H12 | h1 | -0.0712 | H12 | h1 | 0.0817 |
| C8 | c3 | -0.3933 | H13 | h1 | 0.0817 |
| H13 | hc | 0.0813 | C8 | c3 | -0.2218 |
| H14 | hc | 0.0813 | H14 | hc | 0.0967 |
| C9 | c | 1.0897 | H15 | hc | 0.0967 |
| O4 | o | -0.9605 | C9 | c | 0.9965 |
| O5 | o | -0.9605 | O3 | o | -0.6965 |
| O1 | os | -0.1106 | O4 | oh | -0.8408 |
| | | | H16 | ho | 0.5261 |
| | | | O5 | os | -0.4297 |

**Table 13:** Charge information of PETIM (carboxylic functionalised group)

| PETIM Terminal Residue 'ccc' β-galactose (NP) | | |
|---|---|---|
| **Atom** | **GAFF atom type** | **Charge** |
| C1 | c3 | -0.3318 |
| C2 | c3 | 0.3061 |
| H1 | h1 | 0.1166 |
| H2 | h1 | 0.1153 |
| C3 | c3 | 0.1559 |
| C4 | c3 | 0.2838 |
| C5 | c3 | 0.3240 |
| H3 | h1 | 0.1339 |
| H4 | h1 | 0.1015 |
| H5 | h2 | 0.1096 |
| O1 | os | -0.2865 |
| O2 | oh | -0.8601 |
| O3 | oh | -0.8377 |
| O4 | oh | -0.7745 |
| C6 | c3 | 0.3758 |
| H6 | ho | 0.5366 |
| H7 | ho | 0.4949 |
| H8 | ho | 0.5130 |
| H9 | h1 | 0.0565 |
| H10 | h1 | 0.0565 |
| O5 | oh | -0.8411 |
| H11 | ho | 0.5120 |
| O6 | os | -0.2601 |

**Table 14:** Charge information of PETIM (β-galactose functionalised group)

| Radius of Gyration (O-core) | | | | | |
|---|---|---|---|---|---|
| **gen** | **G2** | **G3** | **G4** | **G5** | **G6** |
| **Amine (NP)** | 7.13 ± 0.32 | 9.07 ± 0.18 | 11.55 ± 0.15 | 15.42 ± 0.18 | 18.72 ± 0.13 |
| **Amine (P)** | 8.61 ± 0.89 | 12.28 ± 1.04 | 16.33 ± 1.41 | 21.3 ± 0.56 | 25.65 ± 0.31 |
| **Amine (DP)** | 10.91 ± 0.58 | 15.52 ± 0.84 | 20.12 ± 0.54 | 25.83 ± 0.46 | 32.27 ± 0.51 |
| **$COO^-$ (DeP)** | 8.36 ± 0.09 | 11.19 ± 0.14 | 14.26 ± 0.14 | 19.64 ± 0.12 | 23.69 ± 0.08 |
| **COOH (NP)** | 9.69 ± 0.6 | 12.04 ± 0.37 | 15.33 ± 0.33 | 18.44 ± 0.2 | 21.63 ± 0.11 |
| **COOH (P)** | 12.69 ± 0.93 | 17.64 ± 0.65 | 22.24 ± 0.67 | 27.84 ± 0.54 | 34 ± 0.4 |
| **β-galactose (NP)** | 10.85 ± 0.23 | 14.18 ± 0.19 | 18.68 ± 0.14 | 22.69 ± 0.15 | 27.57 ± 0.07 |
| **β-galactose (P)** | 15.32 ± 0.79 | 19.78 ± 0.57 | 24.61 ± 0.57 | 30.49 ± 0.37 | 37.33 ± 0.27 |
| **Asphericity (O-core)** | | | | | |
| **gen** | **G2** | **G3** | **G4** | **G5** | **G6** |
| **Amine (NP)** | 0.08 ± 0.06 | 0.06 ± 0.02 | 0.05 ± 0.01 | 0.12 ± 0.02 | 0.06 ± 0.01 |
| **Amine (P)** | 0.19 ± 0.2 | 0.2 ± 0.22 | 0.41 ± 0.17 | 0.31 ± 0.07 | 0.09 ± 0.02 |
| **Amine (DP)** | 0.18 ± 0.08 | 0.11 ± 0.06 | 0.04 ± 0.03 | 0.05 ± 0.02 | 0.01 ± 0.01 |
| **$COO^-$ (DeP)** | 0.11 ± 0.02 | 0.09 ± 0.01 | 0.08 ± 0.01 | 0.09 ± 0.01 | 0.02 ± 0.01 |
| **COOH (NP)** | 0.09 ± 0.09 | 0.07 ± 0.03 | 0.06 ± 0.02 | 0.02 ± 0.01 | 0.06 ± 0.01 |
| **COOH (P)** | 0.11 ± 0.1 | 0.13 ± 0.06 | 0.05 ± 0.04 | 0.01 ± 0.01 | 0.01 ± 0.01 |
| **β-galactose (NP)** | 0.13 ± 0.05 | 0.14 ± 0.02 | 0.1 ± 0.01 | 0.06 ± 0.01 | 0.03 ± 0.01 |
| **β-galactose (P)** | 0.12 ± 0.08 | 0.08 ± 0.04 | 0.03 ± 0.02 | 0.02 ± 0.01 | 0.01 ± 0.01 |
| **Aspect ratio ZY (O-core)** | | | | | |
| **gen** | **G2** | **G3** | **G4** | **G5** | **G6** |
| **Amine (NP)** | 1.55 ± 0.42 | 1.42 ± 0.21 | 1.61 ± 0.11 | 2.05 ± 0.2 | 1.74 ± 0.07 |
| **Amine (P)** | 1.99 ± 1.24 | 2.51 ± 0.79 | 4.93 ± 1.41 | 3.84 ± 1 | 1.59 ± 0.14 |
| **Amine (DP)** | 1.71 ± 0.48 | 1.61 ± 0.28 | 1.4 ± 0.24 | 1.55 ± 0.19 | 1.19 ± 0.09 |
| **$COO^-$ (DeP)** | 1.63 ± 0.17 | 1.73 ± 0.1 | 2.01 ± 0.21 | 1.48 ± 0.07 | 1.15 ± 0.04 |
| **COOH (NP)** | 1.49 ± 0.47 | 1.51 ± 0.29 | 1.44 ± 0.39 | 1.34 ± 0.08 | 1.64 ± 0.04 |
| **COOH (P)** | 1.81 ± 0.61 | 1.71 ± 0.27 | 1.43 ± 0.25 | 1.3 ± 0.11 | 1.25 ± 0.08 |
| **β-galactose (NP)** | 2.28 ± 0.32 | 1.69 ± 0.09 | 1.78 ± 0.15 | 1.61 ± 0.08 | 1.25 ± 0.04 |
| **β-galactose (P)** | 1.82 ± 0.35 | 1.37 ± 0.32 | 1.34 ± 0.12 | 1.2 ± 0.17 | 1.17 ± 0.08 |
| **Aspect ratio ZX (O-core)** | | | | | |
| **gen** | **G2** | **G3** | **G4** | **G5** | **G6** |
| **Amine (NP)** | 2.62 ± 0.73 | 2.37 ± 0.31 | 2.17 ± 0.41 | 3.27 ± 0.29 | 2.15 ± 0.07 |
| **Amine (P)** | 4.94 ± 2.17 | 4.92 ± 1.98 | 8.25 ± 2.07 | 6.43 ± 1.44 | 3.15 ± 0.48 |
| **Amine (DP)** | 7.08 ± 2.55 | 3.69 ± 1.02 | 1.94 ± 0.55 | 2.01 ± 0.31 | 1.34 ± 0.18 |
| **$COO^-$ (DeP)** | 3.59 ± 0.83 | 2.98 ± 0.48 | 2.29 ± 0.19 | 3.13 ± 0.09 | 1.61 ± 0.03 |
| **COOH (NP)** | 2.86 ± 0.73 | 2.61 ± 0.38 | 2.46 ± 0.85 | 1.57 ± 0.09 | 2.41 ± 0.06 |
| **COOH (P)** | 3.57 ± 2.31 | 4.22 ± 1.09 | 2.04 ± 0.52 | 1.47 ± 0.14 | 1.47 ± 0.09 |
| **β-galactose (NP)** | 3.24 ± 0.29 | 4.41 ± 0.29 | 3.06 ± 0.19 | 2.38 ± 0.1 | 1.8 ± 0.02 |
| **β-galactose (P)** | 3.92 ± 1.79 | 2.92 ± 0.76 | 1.81 ± 0.27 | 1.63 ± 0.27 | 1.42 ± 0.07 |

**Table 15:** Radius of gyration, asphericity and aspect ratios for PETIM (O-core) dendrimers

| Radius of Gyration (N-core) | | | | | |
|---|---|---|---|---|---|
| **gen** | **G2** | **G3** | **G4** | **G5** | **G6** |
| **Amine (NP)** | 8.77 ± 0.44 | 11.11 ± 0.31 | 14.18 ± 0.2 | 19.24 ± 0.17 | 24.43 ± 0.1 |
| **Amine (P)** | 8.91 ± 0.4 | 13.74 ± 0.53 | 19.01 ± 0.67 | 23.24 ± 0.53 | 28.38 ± 0.3 |
| **Amine (DP)** | 13.52 ± 0.65 | 18.18 ± 0.57 | 23.33 ± 0.33 | 29.51 ± 0.37 | 36.36 ± 0.37 |
| **$COO^-$ (DeP)** | 9.83 ± 0.1 | 13.95 ± 0.23 | 17.04 ± 0.12 | 23.07 ± 0.15 | 26.6 ± 0.08 |
| **COOH (NP)** | 11.72 ± 0.53 | 13.51 ± 0.3 | 17.81 ± 0.18 | 22.13 ± 0.17 | 24.61 ± 0.06 |
| **COOH (P)** | 15.04 ± 0.85 | 19.63 ± 0.88 | 25.09 ± 0.61 | 31.17 ± 0.39 | 38.13 ± 0.37 |
| **β-galactose (NP)** | 12.18 ± 0.21 | 16.7 ± 0.19 | 21.35 ± 0.12 | 25.18 ± 0.08 | 31.49 ± 0.07 |
| **β-galactose (P)** | 16.91 ± 0.75 | 22.8 ± 0.78 | 28.65 ± 0.62 | 34.69 ± 0.34 | 40.9 ± 0.26 |
| **Asphericity (N-core)** | | | | | |
| **gen** | **G2** | **G3** | **G4** | **G5** | **G6** |
| **Amine (NP)** | 0.15 ± 0.1 | 0.14 ± 0.04 | 0.18 ± 0.02 | 0.14 ± 0.02 | 0.03 ± 0 |
| **Amine (P)** | 0.12 ± 0.06 | 0.19 ± 0.09 | 0.23 ± 0.09 | 0.14 ± 0.04 | 0.04 ± 0.01 |
| **Amine (DP)** | 0.13 ± 0.07 | 0.07 ± 0.08 | 0.03 ± 0.01 | 0.02 ± 0.01 | 0.01 ± 0.01 |
| **$COO^-$ (DeP)** | 0.08 ± 0.01 | 0.19 ± 0.02 | 0.08 ± 0.01 | 0.09 ± 0.01 | 0.03 ± 0 |
| **COOH (NP)** | 0.15 ± 0.06 | 0.08 ± 0.03 | 0.12 ± 0.02 | 0.07 ± 0.11 | 0.02 ± 0 |
| **COOH (P)** | 0.09 ± 0.07 | 0.05 ± 0.04 | 0.07 ± 0.03 | 0.02 ± 0.01 | 0.01 ± 0.01 |
| **β-galactose (NP)** | 0.05 ± 0.02 | 0.15 ± 0.02 | 0.08 ± 0.01 | 0.02 ± 0 | 0.01 ± 0 |
| **β-galactose (P)** | 0.11 ± 0.05 | 0.05 ± 0.03 | 0.02 ± 0.02 | 0.01 ± 0.01 | 0.01 ± 0 |
| **Aspect ratio ZY (N-core)** | | | | | |
| **gen** | **G2** | **G3** | **G4** | **G5** | **G6** |
| **Amine (NP)** | 2.43 ± 0.65 | 2.18 ± 0.31 | 2.86 ± 0.35 | 1.58 ± 0.1 | 1.23 ± 0.05 |
| **Amine (P)** | 1.67 ± 0.27 | 2.39 ± 0.66 | 2.37 ± 0.71 | 1.58 ± 0.22 | 1.39 ± 0.09 |
| **Amine (DP)** | 1.48 ± 0.39 | 1.27 ± 0.27 | 1.19 ± 0.14 | 1.13 ± 0.09 | 1.14 ± 0.05 |
| **$COO^-$ (DeP)** | 1.45 ± 0.1 | 3.06 ± 0.29 | 1.7 ± 0.09 | 1.32 ± 0.09 | 1.27 ± 0.02 |
| **COOH (NP)** | 1.9 ± 0.39 | 1.79 ± 0.24 | 1.69 ± 0.13 | 1.75 ± 0.25 | 1.07 ± 0.02 |
| **COOH (P)** | 1.64 ± 0.3 | 1.42 ± 0.17 | 1.5 ± 0.2 | 1.34 ± 0.17 | 1.16 ± 0.06 |
| **β-galactose (NP)** | 1.24 ± 0.08 | 2 ± 0.12 | 1.6 ± 0.05 | 1.27 ± 0.04 | 1.05 ± 0.02 |
| **β-galactose (P)** | 1.48 ± 0.21 | 1.33 ± 0.16 | 1.26 ± 0.11 | 1.09 ± 0.05 | 1.09 ± 0.03 |
| **Aspect ratio ZX (N-core)** | | | | | |
| **gen** | **G2** | **G3** | **G4** | **G5** | **G6** |
| **Amine (NP)** | 3.32 ± 1.21 | 3.54 ± 0.54 | 3.54 ± 0.38 | 4.77 ± 0.46 | 1.86 ± 0.1 |
| **Amine (P)** | 3.81 ± 1.22 | 4.53 ± 1.02 | 6.79 ± 1.06 | 4.88 ± 0.66 | 1.97 ± 0.17 |
| **Amine (DP)** | 4.51 ± 1.25 | 2.83 ± 0.57 | 1.92 ± 0.32 | 1.54 ± 0.12 | 1.49 ± 0.08 |
| **$COO^-$ (DeP)** | 3.06 ± 0.23 | 3.68 ± 0.38 | 2.66 ± 0.12 | 3.3 ± 0.17 | 1.78 ± 0.03 |
| **COOH (NP)** | 4.45 ± 1.24 | 2.54 ± 0.37 | 3.7 ± 0.3 | 2.48 ± 0.39 | 1.58 ± 0.05 |
| **COOH (P)** | 3.07 ± 0.79 | 2.12 ± 0.5 | 2.51 ± 0.53 | 1.57 ± 0.14 | 1.42 ± 0.08 |
| **β-galactose (NP)** | 2.4 ± 0.17 | 4.4 ± 0.27 | 2.92 ± 0.1 | 1.74 ± 0.04 | 1.32 ± 0.03 |
| **β-galactose (P)** | 3.67 ± 1.03 | 2.27 ± 0.37 | 1.59 ± 0.16 | 1.48 ± 0.1 | 1.35 ± 0.12 |

**Table 16:** Radius of gyration, asphericity and aspect ratios for PETIM (N-core) dendrimers.

| O-core | | | | | | | | | |
|---|---|---|---|---|---|---|---|---|---|
| **Gen** | **Inner Gen** | **Amine (NP)** | **Amine (P)** | **Amine (DP)** | **COO- (DeP)** | **COOH (NP)** | **COOH (P)** | **β-galactose (NP)** | **β-galactose (P)** |
| G2 | **g1** | 2 ± 0.36 | 4.11 ± 0.45 | 3.61 ± 0.99 | 1.04 ± 0.31 | 2.98 ± 0.4 | 4.27 ± 1.08 | 1.1 ± 0.11 | 3.96 ± 0.17 |
| | **g2** | 6.02 ± 0.78 | 8.15 ± 0.71 | 8.54 ± 1.08 | 0.99 ± 0.23 | 5.71 ± 1.08 | 8.61 ± 0.86 | 1.08 ± 0.45 | 6.34 ± 0.94 |
| | **g2.5** | | | | 1.89 ± 1.04 | 6.8 ± 2.09 | 9.48 ± 1.9 | 1.73 ± 1.07 | 7.9 ± 1.67 |
| G3 | **g1** | 1.74 ± 0.36 | 3.25 ± 0.35 | 3.36 ± 0.49 | 1.01 ± 0.2 | 1.86 ± 0.26 | 3.6 ± 0.71 | 0.84 ± 0.22 | 3.27 ± 0.54 |
| | **g2** | 2.09 ± 0.69 | 4.65 ± 1.75 | 6.49 ± 1.58 | 1.26 ± 0.34 | 2.02 ± 0.93 | 6.56 ± 1.24 | 0.85 ± 0.29 | 4.65 ± 0.9 |
| | **g3** | 5.47 ± 1.63 | 8.54 ± 2.22 | 10.96 ± 1.51 | 1.5 ± 0.45 | 4.54 ± 1.48 | 9.87 ± 1.88 | 0.94 ± 0.23 | 6.25 ± 1.53 |
| | **g3.5** | | | | 2.03 ± 0.7 | 5.66 ± 2.28 | 10.55 ± 2.29 | 1.26 ± 0.64 | 7.41 ± 2.18 |
| G4 | **g1** | 1.28 ± 0.31 | 4.24 ± 0.95 | 3.06 ± 0.25 | 1.16 ± 0.38 | 1.51 ± 0.3 | 3.46 ± 0.29 | 0.83 ± 0.28 | 3.16 ± 0.21 |
| | **g2** | 1.37 ± 0.42 | 2.92 ± 0.19 | 5.16 ± 0.99 | 1.13 ± 0.44 | 1.92 ± 0.46 | 5.07 ± 0.77 | 0.96 ± 0.26 | 4.15 ± 0.45 |
| | **g3** | 1.61 ± 0.5 | 4.69 ± 1.12 | 8.47 ± 1.35 | 1.2 ± 0.49 | 2.52 ± 1.02 | 7.6 ± 1 | 1.09 ± 0.44 | 5.92 ± 0.91 |
| | **g4** | 4.61 ± 1.39 | 8.76 ± 1.39 | 12.59 ± 1.47 | 1.16 ± 0.35 | 4.65 ± 1.79 | 10.49 ± 1.19 | 1.28 ± 0.55 | 7.53 ± 1.73 |
| | **g4.5** | | | | 1.48 ± 0.59 | 5.72 ± 2.29 | 11.15 ± 1.84 | 1.82 ± 1.14 | 8.86 ± 2.15 |
| G5 | **g1** | 0.99 ± 0.17 | 2.73 ± 0.35 | 2.95 ± 0.38 | 0.75 ± 0.1 | 0.92 ± 0.08 | 3.1 ± 0.41 | 0.89 ± 0.15 | 2.94 ± 0.51 |
| | **g2** | 0.99 ± 0.19 | 2.67 ± 0.62 | 4.63 ± 0.52 | 0.81 ± 0.14 | 1.08 ± 0.29 | 4.07 ± 0.33 | 1.01 ± 0.34 | 3.91 ± 0.5 |
| | **g3** | 1.15 ± 0.37 | 3.34 ± 0.91 | 6.52 ± 0.91 | 0.86 ± 0.17 | 1.06 ± 0.26 | 5.37 ± 0.5 | 0.92 ± 0.2 | 4.71 ± 0.61 |
| | **g4** | 1.83 ± 1.27 | 4.99 ± 1.98 | 9.42 ± 1.33 | 1.04 ± 0.34 | 1.7 ± 0.65 | 7.84 ± 1.36 | 0.99 ± 0.31 | 5.85 ± 0.98 |
| | **g5** | 4.72 ± 2.1 | 8.59 ± 2.4 | 13.08 ± 1.54 | 1.2 ± 0.55 | 3.62 ± 2.02 | 10.42 ± 1.75 | 1.18 ± 0.73 | 7.16 ± 1.71 |
| | **g5.5** | | | | 1.56 ± 0.89 | 4.58 ± 2.58 | 11.11 ± 2.13 | 1.6 ± 1.24 | 8.21 ± 2.1 |
| G6 | **g1** | 1.06 ± 0.34 | 1.86 ± 0.18 | 2.64 ± 0.39 | 0.68 ± 0.09 | 0.69 ± 0.15 | 2.51 ± 0.19 | 0.83 ± 0.19 | 2.01 ± 0.22 |
| | **g2** | 0.9 ± 0.22 | 2.16 ± 0.38 | 4.12 ± 0.44 | 0.73 ± 0.16 | 0.64 ± 0.11 | 3.52 ± 0.29 | 0.73 ± 0.18 | 2.79 ± 0.15 |
| | **g3** | 0.96 ± 0.24 | 2.57 ± 0.75 | 5.68 ± 1.08 | 0.79 ± 0.18 | 0.67 ± 0.1 | 4.51 ± 0.46 | 0.74 ± 0.14 | 3.57 ± 0.4 |
| | **g4** | 1.25 ± 0.4 | 3.6 ± 1.27 | 8.05 ± 1.89 | 0.82 ± 0.23 | 0.82 ± 0.22 | 6.16 ± 0.77 | 0.76 ± 0.18 | 4.28 ± 0.69 |
| | **g5** | 1.66 ± 0.84 | 5.19 ± 2.01 | 10.78 ± 2.46 | 1.02 ± 0.37 | 1.09 ± 0.46 | 8.09 ± 1.48 | 0.88 ± 0.27 | 5.08 ± 1.14 |
| | **g6** | 4.14 ± 1.92 | 8.47 ± 2.5 | 14.17 ± 2.56 | 1.09 ± 0.42 | 1.53 ± 0.99 | 10.44 ± 1.85 | 1.06 ± 0.6 | 6.17 ± 1.66 |
| | **g6.5** | | | | 1.37 ± 0.63 | 2.31 ± 1.62 | 11.13 ± 2.17 | 1.36 ± 1.09 | 7.12 ± 2.22 |
| **N-core** | | | | | | | | | |
| **Gen** | | **Amine (NP)** | **Amine (P)** | **Amine (DP)** | **COO- (DeP)** | **COOH (NP)** | **COOH (P)** | **β-galactose (NP)** | **β-galactose (P)** |
| G2 | **g1** | 2.72 ± 0.87 | 2.13 ± 0.56 | 5.2 ± 1.76 | 0.74 ± 0.19 | 3.08 ± 0.79 | 5.21 ± 1.45 | 0.83 ± 0.22 | 4.03 ± 0.92 |
| | **g2** | 6.25 ± 1.77 | 6.45 ± 1.12 | 11.05 ± 0.92 | 0.89 ± 0.31 | 6.3 ± 1.31 | 9.42 ± 1.41 | 1.18 ± 0.54 | 7.35 ± 1.12 |
| | **g2.5** | | | | 1.47 ± 0.87 | 7.27 ± 2.04 | 10.13 ± 2 | 1.91 ± 1.04 | 8.66 ± 1.69 |
| G3 | **g1** | 1.83 ± 0.32 | 2.56 ± 0.51 | 4.66 ± 1.4 | 1.15 ± 0.37 | 1.33 ± 0.2 | 4.1 ± 0.8 | 0.91 ± 0.18 | 3.7 ± 0.54 |
| | **g2** | 2.01 ± 0.53 | 4.67 ± 1.97 | 8.68 ± 1.44 | 1.06 ± 0.27 | 1.92 ± 0.5 | 6.49 ± 0.71 | 0.88 ± 0.2 | 5.21 ± 0.71 |
| | **g3** | 5.4 ± 1.58 | 8.57 ± 2.61 | 12.61 ± 1.64 | 1.36 ± 0.45 | 4.49 ± 1.64 | 9.21 ± 1.01 | 1.09 ± 0.32 | 7.49 ± 1.39 |
| | **g3.5** | | | | 1.85 ± 0.81 | 5.4 ± 2.22 | 10.04 ± 1.7 | 1.71 ± 0.99 | 9.04 ± 1.91 |
| G4 | **g1** | 1.56 ± 0.41 | 2.38 ± 0.83 | 3.62 ± 0.7 | 1.03 ± 0.39 | 1.33 ± 0.3 | 3.76 ± 0.6 | 1.04 ± 0.19 | 3.22 ± 0.36 |
| | **g2** | 1.55 ± 0.13 | 2.94 ± 1.36 | 6.26 ± 0.79 | 0.91 ± 0.37 | 1.53 ± 0.39 | 5.44 ± 0.41 | 1.02 ± 0.21 | 4.2 ± 0.58 |
| | **g3** | 2.23 ± 0.66 | 4.43 ± 2.1 | 9.22 ± 1.39 | 1.05 ± 0.51 | 1.87 ± 0.6 | 7.56 ± 0.88 | 1.09 ± 0.28 | 5.22 ± 0.83 |
| | **g4** | 5.65 ± 1.74 | 8.12 ± 2.3 | 12.89 ± 1.55 | 1.22 ± 0.63 | 3.84 ± 1.85 | 10.4 ± 1.23 | 1.35 ± 0.55 | 6.8 ± 1.47 |
| | **g4.5** | | | | 1.6 ± 0.94 | 4.75 ± 2.47 | 11.1 ± 1.84 | 1.85 ± 1.15 | 7.96 ± 1.98 |
| G5 | **g1** | 1.13 ± 0.35 | 2.15 ± 0.44 | 2.96 ± 0.52 | 1.13 ± 0.33 | 0.84 ± 0.11 | 3.01 ± 0.47 | 0.66 ± 0.09 | 2.71 ± 0.32 |
| | **g2** | 1.19 ± 0.23 | 2.32 ± 0.5 | 4.95 ± 0.62 | 1.12 ± 0.33 | 1.03 ± 0.23 | 4.47 ± 0.61 | 0.66 ± 0.09 | 3.73 ± 0.37 |
| | **g3** | 1.36 ± 0.42 | 2.87 ± 0.75 | 7.11 ± 1.09 | 1.15 ± 0.3 | 1.25 ± 0.3 | 5.92 ± 1.02 | 0.74 ± 0.17 | 4.63 ± 0.69 |
| | **g4** | 1.98 ± 0.89 | 4.29 ± 1.73 | 10.18 ± 1.64 | 1.33 ± 0.48 | 1.69 ± 0.76 | 8.28 ± 1.4 | 0.83 ± 0.18 | 5.64 ± 1.16 |
| | **g5** | 4.61 ± 2 | 7.81 ± 2.32 | 13.79 ± 1.78 | 1.32 ± 0.45 | 3.12 ± 1.94 | 10.91 ± 1.69 | 0.92 ± 0.33 | 6.79 ± 1.76 |
| | **g5.5** | | | | 1.56 ± 0.71 | 3.94 ± 2.46 | 11.59 ± 2.05 | 1.2 ± 0.72 | 7.64 ± 2.18 |
| G6 | **g1** | 0.87 ± 0.12 | 1.89 ± 0.23 | 2.46 ± 0.42 | 0.76 ± 0.16 | 0.83 ± 0.31 | 2.36 ± 0.38 | 0.89 ± 0.18 | 2.07 ± 0.24 |
| | **g2** | 0.86 ± 0.13 | 2.01 ± 0.62 | 3.62 ± 0.29 | 0.71 ± 0.12 | 0.75 ± 0.21 | 3.48 ± 0.25 | 0.86 ± 0.23 | 2.78 ± 0.3 |
| | **g3** | 1 ± 0.2 | 2.76 ± 0.86 | 4.89 ± 0.79 | 0.74 ± 0.14 | 0.74 ± 0.17 | 4.59 ± 0.57 | 0.84 ± 0.21 | 3.58 ± 0.57 |
| | **g4** | 1.17 ± 0.33 | 3.77 ± 1.26 | 6.89 ± 1.14 | 0.83 ± 0.2 | 0.83 ± 0.2 | 6.23 ± 0.97 | 0.8 ± 0.18 | 4.04 ± 0.66 |
| | **g5** | 1.57 ± 0.74 | 5.08 ± 1.98 | 9.63 ± 1.6 | 0.9 ± 0.25 | 1.03 ± 0.42 | 8.14 ± 1.6 | 0.86 ± 0.25 | 4.76 ± 0.95 |
| | **g6** | 3.83 ± 1.79 | 8.2 ± 2.46 | 13.12 ± 1.75 | 0.9 ± 0.3 | 1.64 ± 1.36 | 10.4 ± 2.07 | 0.92 ± 0.39 | 5.65 ± 1.39 |
| | **g6.5** | | | | 1.13 ± 0.51 | 2.35 ± 1.85 | 11.11 ± 2.32 | 1.16 ± 0.76 | 6.51 ± 1.93 |

**Table 17:** RMSF values of PETIM (O-core and N-core) dendrimers

| Surface water calculation (O-core) | | | | | |
|---|---|---|---|---|---|
| **gen** | **G2** | **G3** | **G4** | **G5** | **G6** |
| **Amine (NP)** | 147.67 ± 7.63 | 257.54 ± 10.03 | 392.93 ± 11.49 | 680.82 ± 18.8 | 1037.2 ± 18.74 |
| **Amine (P)** | 175.14 ± 8.9 | 323.58 ± 11.52 | 606.92 ± 15.57 | 1043.92 ± 23.32 | 1950.52 ± 47.7 |
| **Amine (DP)** | 245.2 ± 9.73 | 535.16 ± 19.52 | 979.91 ± 30.19 | 1973.88 ± 44.5 | 3856.46 ± 58.95 |
| **$COO^-$ (DeP)** | 191.33 ± 6.1 | 333.84 ± 8.83 | 532.81 ± 12.15 | 967.42 ± 15.18 | 1576.23 ± 23.83 |
| **COOH (NP)** | 213.35 ± 11.8 | 384.17 ± 16.71 | 646.78 ± 18.02 | 998.76 ± 22.65 | 1238.05 ± 23.59 |
| **COOH (P)** | 312.53 ± 14.29 | 651.1 ± 22.56 | 1171.96 ± 58.98 | 2272.54 ± 44.45 | 4162.17 ± 68.06 |
| **β-galactose (NP)** | 333.02 ± 9.17 | 533.11 ± 11.82 | 970.93 ± 17.94 | 1473.31 ± 20.39 | 2300.73 ± 27 |
| **β-galactose (P)** | 494.05 ± 19.37 | 917.2 ± 24.74 | 1739.43 ± 40.38 | 3029.85 ± 85.67 | 5219 ± 116.95 |
| **Bulk water calculation (O-core)** | | | | | |
| **gen** | **G2** | **G3** | **G4** | **G5** | **G6** |
| **Amine (NP)** | 246.94 ± 11.87 | 393.71 ± 16.22 | 557.79 ± 17.12 | 920.95 ± 26.74 | 1349.34 ± 27.49 |
| **Amine (P)** | 294.52 ± 14.87 | 507.23 ± 20.02 | 901.93 ± 29.02 | 1535 ± 42.39 | 2781.44 ± 59.23 |
| **Amine (DP)** | 405.75 ± 19.33 | 857.27 ± 37.51 | 1498.27 ± 51.95 | 2890.35 ± 67.78 | 5508.54 ± 91.23 |
| **$COO^-$ (DeP)** | 305.08 ± 9.15 | 476.72 ± 15.62 | 721.03 ± 14.22 | 1282.28 ± 20.13 | 2040.84 ± 26.52 |
| **COOH (NP)** | 348.32 ± 19.51 | 583.98 ± 24.23 | 961.8 ± 27.58 | 1417.54 ± 40.38 | 1612.1 ± 31.96 |
| **COOH (P)** | 507.73 ± 23.22 | 996.11 ± 43.44 | 1748.39 ± 96.4 | 3239.52 ± 58.13 | 5672.14 ± 75.32 |
| **β-galactose (NP)** | 491.46 ± 13.81 | 716.5 ± 15.23 | 1320.76 ± 25.47 | 1884.52 ± 30.21 | 2962.28 ± 40.44 |
| **β-galactose (P)** | 748.28 ± 37.42 | 1309.56 ± 54.21 | 2386.16 ± 65.41 | 3965.78 ± 88.83 | 6670.12 ± 114.91 |
| **Bound water calculation (O-core)** | | | | | |
| **gen** | **G2** | **G3** | **G4** | **G5** | **G6** |
| **Amine (NP)** | 0.18 ± 0.45 | 1.71 ± 1.44 | 3.97 ± 2.17 | 17.61 ± 5.04 | 56.01 ± 6.11 |
| **Amine (P)** | 1.56 ± 1.5 | 12.07 ± 3.98 | 25.48 ± 7.76 | 91.37 ± 14.28 | 260.9 ± 21.39 |
| **Amine (DP)** | 3.01 ± 5.51 | 16.73 ± 8.1 | 82.32 ± 16.6 | 175.87 ± 27.38 | 413.41 ± 32.67 |
| **$COO^-$ (DeP)** | 0.83 ± 0.78 | 5.3 ± 2.03 | 51.24 ± 5.45 | 132.92 ± 7.72 | 437.02 ± 12.82 |
| **COOH (NP)** | 6.49 ± 2.58 | 21.54 ± 4.8 | 59.37 ± 8.2 | 171.6 ± 10.65 | 177.29 ± 6.47 |
| **COOH (P)** | 13.6 ± 5.06 | 35.65 ± 8.71 | 143.99 ± 27.97 | 310.6 ± 31.37 | 794.99 ± 54.59 |
| **β-galactose (NP)** | 17.1 ± 2.85 | 36.24 ± 4.33 | 123.02 ± 7.36 | 397.05 ± 10.5 | 1010.24 ± 17.22 |
| **β-galactose (P)** | 28.35 ± 8.31 | 87.78 ± 11.99 | 262.63 ± 22.56 | 633.45 ± 56.07 | 1606.61 ± 73.52 |

**Table 18:** Number of bound water molecules in PETIM (O-core) dendrimers.

| Surface water calculation (N-core) | | | | | |
|---|---|---|---|---|---|
| **gen** | **G2** | **G3** | **G4** | **G5** | **G6** |
| **Amine (NP)** | 211.05 ± 9.69 | 338.93 ± 11.7 | 546.08 ± 15.88 | 950.04 ± 20.44 | 1323.09 ± 19.4 |
| **Amine (P)** | 233.98 ± 11.02 | 498.63 ± 17.72 | 832.32 ± 23.66 | 1354.81 ± 29.82 | 2545.38 ± 48.57 |
| **Amine (DP)** | 384.63 ± 15.92 | 745.44 ± 32.08 | 1516.05 ± 33.52 | 2868.06 ± 71.99 | 5227.96 ± 77.33 |
| **$COO^-$ (DeP)** | 262.63 ± 7.09 | 458.39 ± 10.01 | 749.46 ± 12.98 | 1380.3 ± 17.86 | 1963.71 ± 24.6 |
| **COOH (NP)** | 341.37 ± 13.22 | 510.03 ± 14.06 | 862.23 ± 25.4 | 1530 ± 29 | 1667.57 ± 23.72 |
| **COOH (P)** | 443.79 ± 19.22 | 937.19 ± 28.11 | 1698.46 ± 68.13 | 3054.93 ± 55.4 | 5412.67 ± 139.84 |
| **β-galactose (NP)** | 429.37 ± 11.91 | 721.76 ± 15.19 | 1289.34 ± 27.67 | 1845.03 ± 24.18 | 3047 ± 28.47 |
| **β-galactose (P)** | 673.83 ± 24.45 | 1237.93 ± 49.88 | 2558.34 ± 53.42 | 4045.87 ± 83.71 | 5959.72 ± 194.04 |
| **Bulk water calculation (N-core)** | | | | | |
| **gen** | **G2** | **G3** | **G4** | **G5** | **G6** |
| **Amine (NP)** | 335.55 ± 14.65 | 493.9 ± 16.91 | 749.07 ± 26.42 | 1258.66 ± 28.55 | 1649.68 ± 31.71 |
| **Amine (P)** | 377.04 ± 17.55 | 773.5 ± 31.34 | 1232.51 ± 41.97 | 1962.93 ± 45.04 | 3635.31 ± 55.34 |
| **Amine (DP)** | 620.78 ± 33.61 | 1151.47 ± 50.79 | 2265.51 ± 52.58 | 4125.87 ± 98.47 | 7396.23 ± 110.54 |
| **$COO^-$ (DeP)** | 387.5 ± 10.13 | 639.74 ± 13.2 | 998.95 ± 17.34 | 1783.54 ± 23.42 | 2440.33 ± 27.28 |
| **COOH (NP)** | 544.1 ± 23.48 | 765.44 ± 20.99 | 1241.73 ± 50.54 | 1910 ± 70 | 2159.14 ± 34.51 |
| **COOH (P)** | 686.55 ± 33.74 | 1407.17 ± 40.92 | 2470.04 ± 70.22 | 4244.59 ± 57.81 | 7412.34 ± 135.66 |
| **β-galactose (NP)** | 598.41 ± 22.25 | 952 ± 23.56 | 1699.66 ± 38.81 | 2335.65 ± 28.81 | 3902.67 ± 28.16 |
| **β-galactose (P)** | 987.27 ± 34.69 | 1734.59 ± 62.58 | 3437.82 ± 70.19 | 5362.77 ± 96.21 | 7892.32 ± 188.28 |
| **Bound water calculation (N-core)** | | | | | |
| **gen** | **G2** | **G3** | **G4** | **G5** | **G6** |
| **Amine (NP)** | 0.9 ± 1.1 | 4.9 ± 2.63 | 9.44 ± 4.03 | 53.93 ± 6.75 | 113.43 ± 6.83 |
| **Amine (P)** | 1.75 ± 1.53 | 20.9 ± 6.31 | 51.53 ± 10.43 | 167.73 ± 17.15 | 578.69 ± 25.96 |
| **Amine (DP)** | 15.23 ± 7.78 | 48.73 ± 14.01 | 125.22 ± 21.02 | 309.88 ± 44.73 | 920.91 ± 53.78 |
| **$COO^-$ (DeP)** | 5.16 ± 1.77 | 20.43 ± 4.1 | 74.62 ± 6.19 | 223.66 ± 9.08 | 724.46 ± 15.76 |
| **COOH (NP)** | 10.29 ± 4.46 | 38.88 ± 6.47 | 103.8 ± 9.12 | 270 ± 13 | 329.1 ± 8.92 |
| **COOH (P)** | 32.71 ± 8.83 | 75.76 ± 16.49 | 238.88 ± 32.6 | 632.7 ± 26.89 | 1560.79 ± 65.37 |
| **β-galactose (NP)** | 23.74 ± 3.95 | 79.28 ± 6.34 | 269.46 ± 13.48 | 710.51 ± 12.55 | 1740.78 ± 16.87 |
| **β-galactose (P)** | 69.85 ± 14.22 | 197.96 ± 29.98 | 397.55 ± 37.23 | 1270.23 ± 42.05 | 3528.62 ± 155.4 |

**Table 19:** Number of bound water molecules in PETIM (N-core) dendrimers.

| Radius of Gyration (PAMAM) | | | | | |
|---|---|---|---|---|---|
| **gen** | **G1** | **G2** | **G3** | **G4** | **G5** |
| **Amine (NP)** | 6.98 ± 0.37 | 9.32 ± 0.3 | 12.47 ± 0.24 | 15.45 ± 0.15 | 20 ± 0.1 |
| **Amine (P)** | 8.93 ± 0.61 | 11.23 ± 0.56 | 15.18 ± 0.54 | 19.43 ± 0.4 | 24.32 ± 0.29 |
| **Amine (DP)** | 10.12 ± 0.4 | 14.64 ± 0.52 | 19.37 ± 0.55 | 24.53 ± 0.41 | 30.02 ± 0.33 |
| **$COO^-$ (DeP)** | 8.66 ± 0.36 | 11.86 ± 0.17 | 14.85 ± 0.09 | 18.85 ± 0.12 | 24.02 ± 0.09 |
| **COOH (NP)** | 8.84 ± 0.37 | 10.7 ± 0.28 | 13.72 ± 0.16 | 17.28 ± 0.15 | 22.2 ± 0.09 |
| **COOH (P)** | 11.26 ± 0.57 | 16.4 ± 0.61 | 20.51 ± 0.53 | 25.65 ± 0.48 | 30.32 ± 0.36 |
| **D-Glucose (NP)** | 11.29 ± 0.27 | 13.21 ± 0.14 | 16.65 ± 0.1 | 21.05 ± 0.07 | 25.96 ± 0.05 |
| **D-Glucose (P)** | 11.73 ± 0.41 | 17.12 ± 0.4 | 21.19 ± 0.42 | 25.84 ± 0.44 | 31.07 ± 0.2 |
| **Asphericity (PAMAM)** | | | | | |
| **gen** | **G1** | **G2** | **G3** | **G4** | **G5** |
| **Amine (NP)** | 0.16 ± 0.11 | 0.12 ± 0.05 | 0.05 ± 0.02 | 0.04 ± 0.01 | 0.03 ± 0 |
| **Amine (P)** | 0.16 ± 0.14 | 0.17 ± 0.11 | 0.1 ± 0.09 | 0.07 ± 0.02 | 0.04 ± 0.01 |
| **Amine (DP)** | 0.15 ± 0.09 | 0.05 ± 0.03 | 0.05 ± 0.02 | 0.02 ± 0.01 | 0.01 ± 0.01 |
| **$COO^-$ (DeP)** | 0.08 ± 0.07 | 0.15 ± 0.02 | 0.05 ± 0.01 | 0.03 ± 0 | 0.02 ± 0.01 |
| **COOH (NP)** | 0.12 ± 0.08 | 0.06 ± 0.03 | 0.05 ± 0.01 | 0.04 ± 0.01 | 0.08 ± 0.01 |
| **COOH (P)** | 0.15 ± 0.08 | 0.07 ± 0.04 | 0.06 ± 0.04 | 0.01 ± 0.01 | 0.01 ± 0 |
| **D-Glucose (NP)** | 0.14 ± 0.05 | 0.11 ± 0.02 | 0.08 ± 0.01 | 0.03 ± 0 | 0.03 ± 0.01 |
| **D-Glucose (P)** | 0.14 ± 0.1 | 0.14 ± 0.04 | 0.03 ± 0.01 | 0.02 ± 0.02 | 0.02 ± 0.01 |
| **Aspect ratio ZY (PAMAM)** | | | | | |
| **gen** | **G1** | **G2** | **G3** | **G4** | **G5** |
| **Amine (NP)** | 2.01 ± 0.56 | 2.13 ± 0.23 | 1.44 ± 0.27 | 1.43 ± 0.06 | 1.52 ± 0.06 |
| **Amine (P)** | 2.09 ± 0.48 | 2.1 ± 0.42 | 1.46 ± 0.36 | 1.53 ± 0.27 | 1.37 ± 0.11 |
| **Amine (DP)** | 1.4 ± 0.31 | 1.23 ± 0.23 | 1.31 ± 0.12 | 1.19 ± 0.07 | 1.19 ± 0.07 |
| **$COO^-$ (DeP)** | 1.63 ± 0.26 | 2.13 ± 0.25 | 1.66 ± 0.06 | 1.43 ± 0.06 | 1.23 ± 0.02 |
| **COOH (NP)** | 1.93 ± 0.62 | 1.29 ± 0.12 | 1.4 ± 0.08 | 1.13 ± 0.05 | 1.53 ± 0.04 |
| **COOH (P)** | 2.06 ± 0.5 | 1.68 ± 0.3 | 1.26 ± 0.16 | 1.21 ± 0.11 | 1.19 ± 0.07 |
| **D-Glucose (NP)** | 1.6 ± 0.2 | 2.11 ± 0.1 | 1.93 ± 0.08 | 1.14 ± 0.03 | 1.17 ± 0.02 |
| **D-Glucose (P)** | 2.36 ± 0.51 | 1.92 ± 0.35 | 1.24 ± 0.09 | 1.15 ± 0.05 | 1.33 ± 0.05 |
| **Aspect ratio ZX (PAMAM)** | | | | | |
| **gen** | **G1** | **G2** | **G3** | **G4** | **G5** |
| **Amine (NP)** | 4.72 ± 0.89 | 3.19 ± 0.4 | 2.08 ± 0.51 | 2.07 ± 0.13 | 1.69 ± 0.05 |
| **Amine (P)** | 4.72 ± 2.04 | 4.93 ± 1.19 | 3.31 ± 0.54 | 2.65 ± 0.48 | 2.06 ± 0.17 |
| **Amine (DP)** | 6.46 ± 2.6 | 2.23 ± 0.87 | 2.2 ± 0.39 | 1.55 ± 0.15 | 1.35 ± 0.08 |
| **$COO^-$ (DeP)** | 2.61 ± 0.64 | 4.1 ± 0.27 | 2.12 ± 0.07 | 1.69 ± 0.05 | 1.55 ± 0.03 |
| **COOH (NP)** | 3.15 ± 1.26 | 2.53 ± 0.33 | 2.24 ± 0.26 | 1.99 ± 0.07 | 2.99 ± 0.08 |
| **COOH (P)** | 4.25 ± 1.34 | 2.53 ± 0.46 | 2.42 ± 0.42 | 1.36 ± 0.21 | 1.43 ± 0.13 |
| **D-Glucose (NP)** | 4.88 ± 0.65 | 2.94 ± 0.2 | 2.38 ± 0.1 | 1.91 ± 0.04 | 1.86 ± 0.02 |
| **D-Glucose (P)** | 3.46 ± 1.05 | 4.23 ± 0.76 | 1.9 ± 0.24 | 1.73 ± 0.12 | 1.62 ± 0.05 |

**Table 20:** Radius of gyration, asphericity and aspect ratios for PAMAM dendrimers

| PAMAM | | | | | | | | | |
|---|---|---|---|---|---|---|---|---|---|
| **Gen** | **Inner Gen** | **Amine (NP)** | **Amine (P)** | **Amine (DP)** | **COO- (DeP)** | **COOH (NP)** | **COOH (P)** | **D-Glucose (NP)** | **D-Glucose (P)** |
| G1 | **g0** | 1.61 ± 0.26 | 3.3 ± 0.92 | 2.85 ± 0.78 | 0.84 ± 0.09 | 1.43 ± 0.23 | 4.48 ± 1.77 | 1.34 ± 0.35 | 1.27 ± 0.47 |
| | **g1** | 3.77 ± 0.65 | 5.81 ± 0.94 | 5.6 ± 0.67 | 1.01 ± 0.14 | 2.85 ± 1 | 7.57 ± 1.14 | 1.82 ± 0.47 | 1.75 ± 0.41 |
| | | | | | 2.02 ± 1.08 | 5.79 ± 1.48 | 10.56 ± 1.1 | 2.87 ± 1.69 | 2.7 ± 1.66 |
| G2 | **g0** | 0.87 ± 0.07 | 3.13 ± 0.64 | 2.73 ± 0.41 | 0.99 ± 0.21 | 0.57 ± 0.08 | 4.75 ± 1.37 | 0.65 ± 0.22 | 2.26 ± 0.33 |
| | **g1** | 1.19 ± 0.39 | 5.85 ± 1.54 | 6.89 ± 0.64 | 1.19 ± 0.34 | 0.75 ± 0.12 | 8.92 ± 1.44 | 0.9 ± 0.31 | 2.66 ± 0.57 |
| | **g2** | 2.36 ± 1.16 | 7.91 ± 1.63 | 9.19 ± 0.83 | 1.55 ± 1.03 | 1.09 ± 0.27 | 10.5 ± 1.5 | 1.04 ± 0.38 | 2.98 ± 1.01 |
| | | | | | 2.34 ± 1.95 | 3.06 ± 1.35 | 13 ± 1.57 | 1.73 ± 1.33 | 4.3 ± 2.13 |
| G3 | **g0** | 1.18 ± 0.21 | 2.25 ± 0.37 | 2.78 ± 0.46 | 0.62 ± 0.05 | 0.9 ± 0.41 | 2.64 ± 0.33 | 0.62 ± 0.07 | 2.45 ± 0.32 |
| | **g1** | 1.51 ± 0.42 | 3.16 ± 0.71 | 5.25 ± 1.19 | 0.8 ± 0.19 | 1.17 ± 0.42 | 4.31 ± 0.37 | 0.74 ± 0.23 | 3.06 ± 0.54 |
| | **g2** | 2.26 ± 1.09 | 5.02 ± 1.77 | 8.93 ± 1.85 | 0.89 ± 0.31 | 1.26 ± 0.28 | 7.05 ± 1.13 | 0.78 ± 0.24 | 4 ± 1.02 |
| | **g3** | 3.74 ± 2.1 | 6.83 ± 1.9 | 10.92 ± 2.01 | 0.95 ± 0.32 | 1.72 ± 0.87 | 8.54 ± 1.65 | 0.95 ± 0.34 | 4.48 ± 1.09 |
| | | | | | 1.4 ± 0.81 | 3.36 ± 2 | 11.23 ± 2.22 | 1.87 ± 1.36 | 5.41 ± 2.31 |
| G4 | **g0** | 0.68 ± 0.13 | 1.98 ± 0.26 | 2.31 ± 0.31 | 0.77 ± 0.3 | 0.6 ± 0.12 | 2.57 ± 0.31 | 0.47 ± 0.06 | 1.78 ± 0.42 |
| | **g1** | 0.84 ± 0.14 | 2.93 ± 1.24 | 4.71 ± 1.27 | 0.82 ± 0.31 | 0.63 ± 0.11 | 3.93 ± 0.7 | 0.49 ± 0.06 | 2.04 ± 0.48 |
| | **g2** | 0.9 ± 0.25 | 4.51 ± 2.4 | 7.18 ± 2.26 | 0.95 ± 0.39 | 0.79 ± 0.3 | 5.41 ± 1.02 | 0.57 ± 0.13 | 2.45 ± 0.74 |
| | **g3** | 1.37 ± 0.66 | 6.23 ± 3.2 | 10.53 ± 2.56 | 1.03 ± 0.33 | 1.07 ± 0.54 | 7.66 ± 1.2 | 0.66 ± 0.22 | 2.8 ± 0.84 |
| | **g4** | 2.83 ± 1.69 | 7.63 ± 3.43 | 12.42 ± 2.62 | 1.12 ± 0.33 | 1.41 ± 0.96 | 9.01 ± 1.42 | 0.8 ± 0.27 | 3.04 ± 0.97 |
| | | | | | 1.53 ± 0.8 | 2.95 ± 1.9 | 11.48 ± 1.87 | 1.43 ± 1.05 | 3.71 ± 1.77 |
| G5 | **g0** | 0.69 ± 0.13 | 1.69 ± 0.26 | 1.96 ± 0.3 | 0.59 ± 0.12 | 0.56 ± 0.12 | 1.95 ± 0.21 | 0.42 ± 0.06 | 1.36 ± 0.14 |
| | **g1** | 0.87 ± 0.3 | 1.88 ± 0.42 | 3.54 ± 0.25 | 0.68 ± 0.14 | 0.57 ± 0.06 | 2.73 ± 0.15 | 0.44 ± 0.05 | 1.58 ± 0.36 |
| | **g2** | 0.9 ± 0.26 | 2.39 ± 0.98 | 5.44 ± 0.92 | 0.7 ± 0.18 | 0.57 ± 0.12 | 3.76 ± 0.58 | 0.5 ± 0.07 | 1.61 ± 0.38 |
| | **g3** | 1.03 ± 0.37 | 3.46 ± 1.7 | 7.38 ± 1.5 | 0.85 ± 0.35 | 0.65 ± 0.17 | 4.94 ± 1.01 | 0.53 ± 0.11 | 1.79 ± 0.39 |
| | **g4** | 1.55 ± 0.75 | 5.22 ± 2.27 | 10.14 ± 1.96 | 1 ± 0.42 | 0.86 ± 0.38 | 6.52 ± 1.67 | 0.6 ± 0.18 | 1.88 ± 0.5 |
| | **g5** | 2.82 ± 1.56 | 6.63 ± 2.49 | 11.84 ± 2.06 | 1.09 ± 0.39 | 1.1 ± 0.58 | 7.58 ± 2.02 | 0.74 ± 0.26 | 2.04 ± 0.68 |
| | | | | | 1.49 ± 0.71 | 2.11 ± 1.47 | 10.11 ± 2.83 | 1.32 ± 1.12 | 2.78 ± 1.74 |

**Table 21:** RMSF values of PAMAM dendrimers

| Surface water calculation (PAMAM) | | | | | |
|---|---|---|---|---|---|
| **gen** | **G1** | **G2** | **G3** | **G4** | **G5** |
| **Amine (NP)** | 151.87 ± 7.22 | 238.95 ± 7.51 | 446.93 ± 13.48 | 722.64 ± 14.51 | 1270.56 ± 22.99 |
| **Amine (P)** | 193.48 ± 11.87 | 326.12 ± 14.43 | 631.04 ± 27.09 | 1100.97 ± 34.93 | 1858.53 ± 47.1 |
| **Amine (DP)** | 224.37 ± 7.94 | 478.38 ± 16.33 | 943.06 ± 26.6 | 1836.43 ± 51.53 | 3322.5 ± 93.37 |
| **$COO^-$ (DeP)** | 188.71 ± 6.12 | 338.65 ± 9.08 | 549.42 ± 10.86 | 977.95 ± 16.15 | 1643.26 ± 22.97 |
| **COOH (NP)** | 195.46 ± 10.32 | 310.47 ± 9.36 | 526.7 ± 13.37 | 866.35 ± 20.99 | 1277.48 ± 23.97 |
| **COOH (P)** | 272.69 ± 10.49 | 550.66 ± 15.59 | 1015.32 ± 49 | 2056.37 ± 62.73 | 3197.65 ± 82.73 |
| **D-Glucose (NP)** | 313.01 ± 11.06 | 476.81 ± 10.69 | 780.37 ± 15.88 | 1274.53 ± 20.26 | 1981.63 ± 24.69 |
| **D-Glucose (P)** | 320.5 ± 10.06 | 663.8 ± 14.61 | 1229.44 ± 27.31 | 2099.39 ± 48.35 | 2952.11 ± 64.63 |
| **Bulk water calculation (PAMAM)** | | | | | |
| **gen** | **G1** | **G2** | **G3** | **G4** | **G5** |
| **Amine (NP)** | 257.94 ± 13.7 | 363.38 ± 11.65 | 649.56 ± 20.61 | 991.09 ± 25.39 | 1687.73 ± 29.52 |
| **Amine (P)** | 336.79 ± 26.88 | 520.64 ± 25.9 | 977.36 ± 38.32 | 1662.75 ± 64.93 | 2774.47 ± 79.77 |
| **Amine (DP)** | 397.92 ± 17.06 | 808.45 ± 34.96 | 1497.68 ± 42.01 | 2787.68 ± 64.27 | 4804.57 ± 126.49 |
| **$COO^-$ (DeP)** | 298.91 ± 8.59 | 506.76 ± 13.06 | 769.46 ± 13.92 | 1287.82 ± 22.2 | 2100.19 ± 26.15 |
| **COOH (NP)** | 324.05 ± 20.05 | 465.28 ± 13.75 | 770.23 ± 20.27 | 1197.09 ± 43.57 | 1677.48 ± 27.16 |
| **COOH (P)** | 442 ± 18.24 | 854.51 ± 27.44 | 1483.44 ± 63.45 | 2939.23 ± 73.87 | 4413.72 ± 77.3 |
| **D-Glucose (NP)** | 469.65 ± 18.03 | 664.3 ± 17.4 | 1055.63 ± 24.35 | 1670.74 ± 23.48 | 2519.29 ± 43.44 |
| **D-Glucose (P)** | 459.69 ± 16.78 | 952.2 ± 25.59 | 1683.24 ± 38.47 | 2826.18 ± 51.98 | 3900.96 ± 104.21 |
| **Bound water calculation (PAMAM)** | | | | | |
| **gen** | **G1** | **G2** | **G3** | **G4** | **G5** |
| **Amine (NP)** | 1.26 ± 1.26 | 5.83 ± 1.78 | 40.07 ± 5.25 | 92.56 ± 6.58 | 269.2 ± 12.27 |
| **Amine (P)** | 4 ± 2.41 | 21.96 ± 5.3 | 52.78 ± 10.23 | 161.3 ± 16.13 | 431.66 ± 24.04 |
| **Amine (DP)** | 2.99 ± 2.38 | 18.11 ± 6.93 | 51.52 ± 17.17 | 148.16 ± 28.85 | 440.79 ± 52.9 |
| **$COO^-$ (DeP)** | 3.59 ± 1.61 | 22.26 ± 3.02 | 88.94 ± 5.25 | 250.84 ± 10.26 | 605.33 ± 15.05 |
| **COOH (NP)** | 4.93 ± 2.47 | 9.6 ± 2.65 | 56 ± 5.15 | 153.53 ± 10.1 | 270.9 ± 7.81 |
| **COOH (P)** | 6.78 ± 3.66 | 31.4 ± 7.66 | 127.03 ± 24.53 | 292.43 ± 34.42 | 885.58 ± 48.63 |
| **D-Glucose (NP)** | 14.53 ± 2.81 | 30.2 ± 3.18 | 88.15 ± 7.64 | 248.9 ± 8.68 | 722.74 ± 13.49 |
| **D-Glucose (P)** | 13.92 ± 3.22 | 56.13 ± 7.24 | 199.62 ± 23.26 | 517.38 ± 31.31 | 1393.9 ± 39.86 |

**Table 22:** Number of bound water molecules in PAMAM dendrimers
